# Integrative multi-omics combined with functional pharmacological profiling in patient-derived organoids identifies personalized therapeutic vulnerabilities of adult high-grade gliomas

**DOI:** 10.1101/2025.09.09.675145

**Authors:** Luca Ermini, Anuja Lipsa, Ann-Christin Hau, Iryna Krokhmal, Bakhtiyor Nosirov, Reka Toth, Eliane Klein, Anaïs Oudin, Linsey Houben, Sabrina Fritah, Christel Herold-Mende, Katrin B.M. Frauenknecht, Michel Mittelbronn, Guy Berchem, Frank Hertel, Tathiane M. Malta, Petr V. Nazarov, Simone P. Niclou, Anna Golebiewska

## Abstract

**Background:** Precision medicine has transformed cancer treatment by tailoring therapies to specific molecular aberrations. Integrating high-resolution multi-omics with high-throughput functional profiling in patient-derived organoids offers a powerful strategy to further refine patient stratification. While (epi)genetic profiling has drastically improved the classification in diffuse adult gliomas, these advances have not yet translated into effective therapeutic interventions and precision medicine approaches remain to be established.

**Material and Methods:** We investigated a panel of 48 patient-derived organoid and orthotopic xenograft models of adult high-grade gliomas, comprehensively characterized at genomic, epigenomic and transcriptomic levels. A functional drug screen was performed on 27 organoid models using a 202-compound library targeting cancer-related pathways and epigenetic regulators. Unsupervised multi-omics factor analysis was employed to identify patient-specific therapeutic vulnerabilities. Validation included dose-dependent drug efficacy assessments, as well as biomarker assessment in patient tumors across molecular subgroups.

**Results:** Multi-omics analysis revealed a broad spectrum of molecular profiles capturing the genetic, epigenetic, and transcriptomic diversity of high-grade gliomas. Multi-omics factor analysis, integrating multi-omics and drug response profiles, identified distinct subgroups associated with *IDH1* mutation and *MYCN* amplification. IDH1 mutant grade 4 astrocytomas showed selective sensitivity to histone deacetylase 3 inhibitors, while a *MYCN*-amplified glioblastoma responded preferentially to histone methyltransferase inhibitors. The differential drug responses were linked to specific (epi)genetic and transcriptomic biomarkers. While other glioblastomas exhibited heterogeneous treatment responses, no robust biomarker-defined responder subgroups were identified.

**Conclusion:** Our findings highlight the value of integrating multi-omics and functional profiling to inform precision medicine strategies. This approach enables the stratification of distinct patient subgroups in preclinical models, paving the way for tailored therapeutic interventions. While we observed distinct pharmacogenomic profiles in IDH1 mutant grade 4 astrocytomas and a *MYCN*-amplified glioblastoma, implementing precision medicine in other glioblastoma subtypes remains a substantial challenge.

**Key points:**

- Integrating drug screening in a panel of patient-derived organoids with multi-omics enables pharmacogenomic profiling in adult diffuse high-grade gliomas
- IDH1 mutant grade 4 astrocytomas are sensitive to histone deacetylase 3 inhibitors
- *MYCN*-amplified glioblastoma exhibits distinct DNA methylation pattern and drug responses

**Study importance:** To date, attempts to develop effective precision medicine in adult high-grade gliomas failed. Here, we provide a preclinical framework for identifying personalized therapeutic by integrating multi-omics profiling with functional drug screening in patient-derived organoids. We show that IDH1 mutant high-grade astrocytomas present distinct therapeutic vulnerabilities compared to glioblastomas, linked to sensitivity to histone deacetylase 3 inhibitors. Within glioblastomas, we identified a distinct *MYCN*-amplified tumor, sensitive to histone methyltransferase inhibitors. Applying pharmacogenomic approaches using novel drug libraries holds promise for uncovering additional clinically relevant patient subgroups in the future.

**Graphical abstract:** 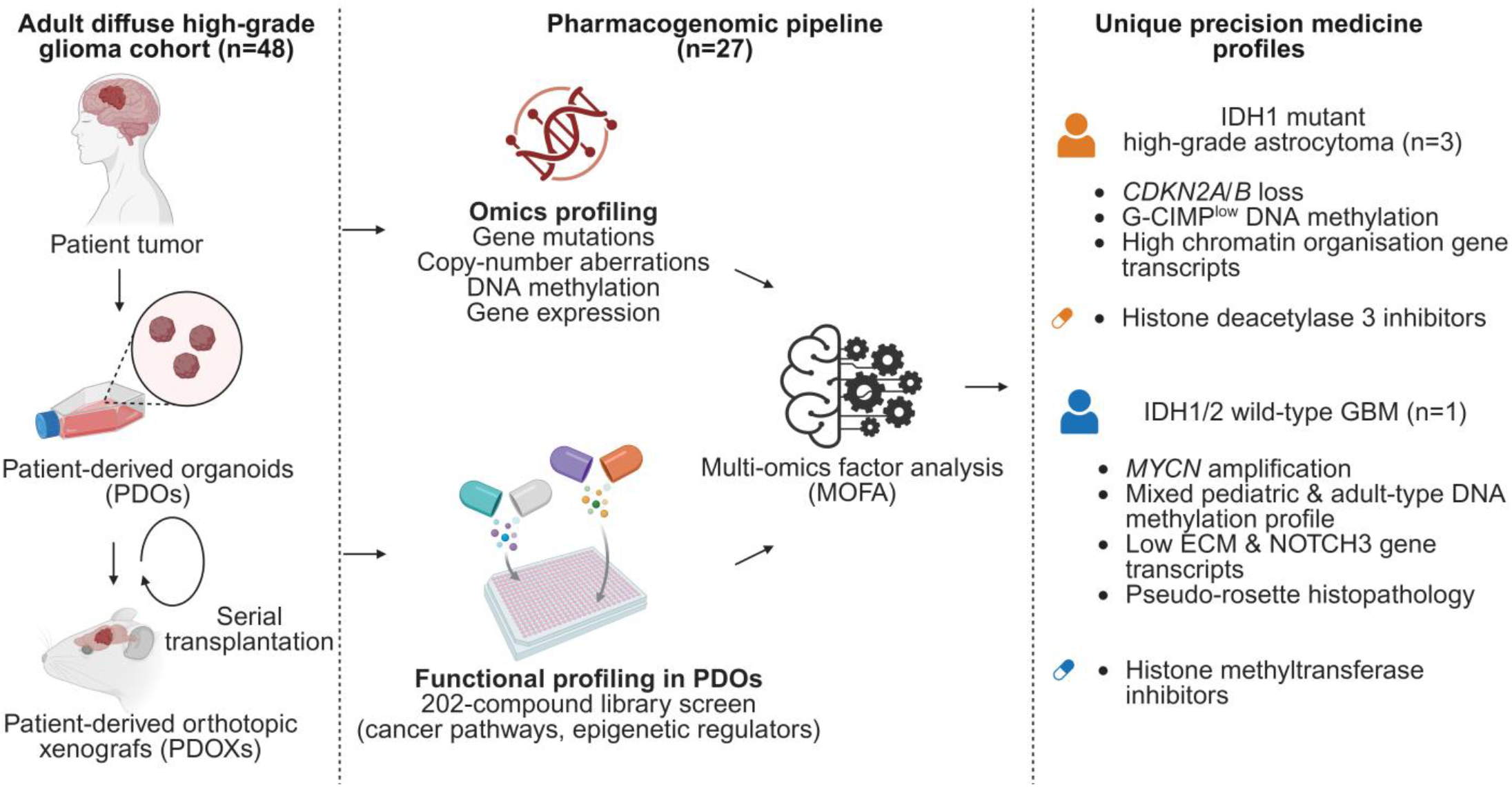

## Introduction

Precision medicine has revolutionized cancer treatment by tailoring therapies to distinct somatic aberrations. It relies on strong predictive and actionable biomarkers that can guide targeted therapies and prognosis, such as *BCR-ABL* fusion kinase in chronic myeloid leukemia, *EGFR* mutations in lung cancer and *BRAF* V600E mutation in melanoma^1^. However, such somatic variants with strong clinical significance are rare and numerous confounding factors within the tumor ecosystem contribute to the resistance to targeted treatment^2^. At the same time, while assessing drug efficacy in patient-derived tumor cells can identify therapeutic hits for the individualized treatment, this approach does not yield biomarkers for stratifying patient groups. To overcome these limitations, an integrated approach combines diverse multi-omics datasets with functional pharmacological profiling, frequently in the framework of drug repurposing^3^. So-called functional precision medicine uniquely enables the identification of actionable vulnerabilities by translating (epi)genetic alterations into functional changes in cellular pathways and phenotypes, while also supporting biomarker-driven drug repurposing strategies. While integrative approaches have demonstrated strong potential in elucidating novel therapeutic targets and biomarkers^3–5^, their clinical implementation remains experimental due to challenges in scalability, standardization of high-throughput screening, and the need for robust validation in diverse patient cohorts. Such approaches rely on the preclinical cohorts with deep molecular characterization suitable for high-throughput drug screening, coupled with advanced machine learning methodologies to integrate multi-omics layers with functional profiling. Among various tumor modeling systems, three-dimensional patient-derived organoids (PDOs) combined with sensitive viability assays represent an appealing compromise between the clinical relevance of conventional cell lines and labor-intensive tissue explants or patient-derived orthotopic xenografts (PDOXs)^6,7^.

Adult-type high-grade gliomas represent a prime candidate for applying functional precision medicine strategies due to their aggressive nature, dismal prognosis and high resistance mechanisms. They constitute roughly 70% of all malignant primary brain tumors, with the large majority diagnosed as glioblastoma (GBM). The 2021 WHO classification characterizes all GBMs as isocitrate dehydrogenase 1/2 (IDH1/2) wild-type (IDH1/2wt), whereas rarer IDH1 and IDH2 mutant (IDH1/2m) high-grade gliomas (grade 3-4) are currently defined as astrocytomas (8-15%) and oligodendrogliomas (3-5%), often as a progression from low-grade tumors^8,9^. At the molecular level, GBMs show frequently +7/-10 chromosome changes, *TERT* promoter mutation and *EGFR* amplification. *IDH1 R132H* mutation is the most common among IDH1 mutant (IDH1m) gliomas. IDH1/2m astrocytomas are distinguished from oligodendrogliomas by lack of 1p/19q-codeletion with frequent *ATRX* and *TP53* mutations. Progression to high-grade is linked to MRI-based contract-enhancing disease and *CDKN2A/B* loss, which also represents classical hallmarks of GBMs^10,11^. These entities differ from pediatric-type high-grade gliomas, which often present mutations in histone 3 (H3) genes^8,9^.

Clinically, GBM and grade 3/4 IDH1/2m adult glioma patients follow similar, largely ineffective, treatment regime comprising maximal safe surgical resection, followed by radio and chemotherapy (astocytomas: temozolomide, oligodendrogliomas: PVC) due to their histological and behavioural similarities, despite molecular differences^12^. While IDH1 inhibitors are ineffective in high-grade IDH1/2m gliomas, it is currently unclear if these tumors show different vulnerabilities compared to GBMs. Rarer high-grade glioma subtypes such as tumors with features of gliosarcoma or primitive neuronal component, may require adapted approaches but often align with core glioma therapies^13^. Despite significant list of molecular aberrations amenable for precision medicine, targeted therapies in gliomas showed very limited success so far^14^, and *MGMT* promoter methylation remains the sole prognostic marker to chemotherapeutic treatment with temozolomide. Indeed, efficacy studies in several preclinical glioma cohorts confirmed the higher susceptibility of *MGMT* promoter methylated models to temozolomide^15,16^. A significant obstacle to improving clinical outcomes is the molecular and functional heterogeneity of these tumors, which encompasses diverse genetic drives, heterogeneous clones, phenotypic states, microenvironmental niches, and adaptive mechanisms of therapy resistance^17^. Expanding personalized treatment strategies to address this complexity remains a critical challenge^18^.

To provide a clinically-relevant platform for testing precision medicine strategies, we applied a large cohort of high-grade gliomas^15^, which currently encompasses 48 molecularly characterized patient-derived models derived from adult patients, including IDH1/2wt GBMs and IDH1m (*R132H)* high-grade astrocytomas. Our protocol allows for indefinite amplification of patient tumor material as PDOs and PDOXs^19^. We have previously demonstrated that these models faithfully replicate the histopathological, genetic, epigenetic, and transcriptomic characteristics of diverse primary and recurrent gliomas and are amenable for drug testing for precision medicine^15,20–25^. In this study, we optimized PDO generation protocol for high-throughput drug screening and evaluated responses to a 202-drug library in 27 PDO models. By applying multi-omics factor analysis (MOFA)^26^, a machine learning approach, we integrated molecular features with drug efficacy profiling to investigate pharmacologically active compounds specific to molecular profiles. We identified distinctive vulnerabilities specific to IDH1m astrocytomas, linked to histone deacetylase 3 (HDAC3) inhibitors. Among GBMs, we identified a unique *MYCN*-amplified model, presenting specific epigenetic features reminiscent of mixed adult- and pediatric-type gliomas linked to distinct drug vulnerabilities. We show that such multi-omics approach, combined with highthroughput functional profiling and drug repurposing, has the potential to refine patient stratification and identify novel molecular subtypes with specific drug vulnerabilities.

## Results

### A preclinical cohort of patient-derived models for precision medicine protocols in diffuse adult high-grade gliomas

To evaluate precision medicine approaches in diffuse high-grade gliomas, we developed a cohort of 48 stable preclinical models derived from 37 individual adult patients, suitable for *ex vivo* and *in vivo* testing **(Table S1)**. This cohort was established using a streamlined protocol involving short-term cultures of primary PDOs, followed by intracranial implantation to generate PDOXs^15,19,23^ (**Fig. 1A**). We have previously shown that this approach minimizes external stressors, such as enzymatic dissociation and extended *in vitro* passaging, preserving the molecular and functional features of patient-derived tumor cells. The cohort represents a diverse patient population with variations in sex, age, and disease stage. A comprehensive characterization of the multi-omics profiling, including targeted DNA sequencing, genome-wide DNA methylation, and bulk RNA sequencing, revealed diverse tumor entities, encompassing 38 IDH1/2wt GBMs and 4 IDH1m (*R132H*) high-grade gliomas (**Fig. 1B**). None of the models presented *IDH2* mutation.

**Figure 1.**
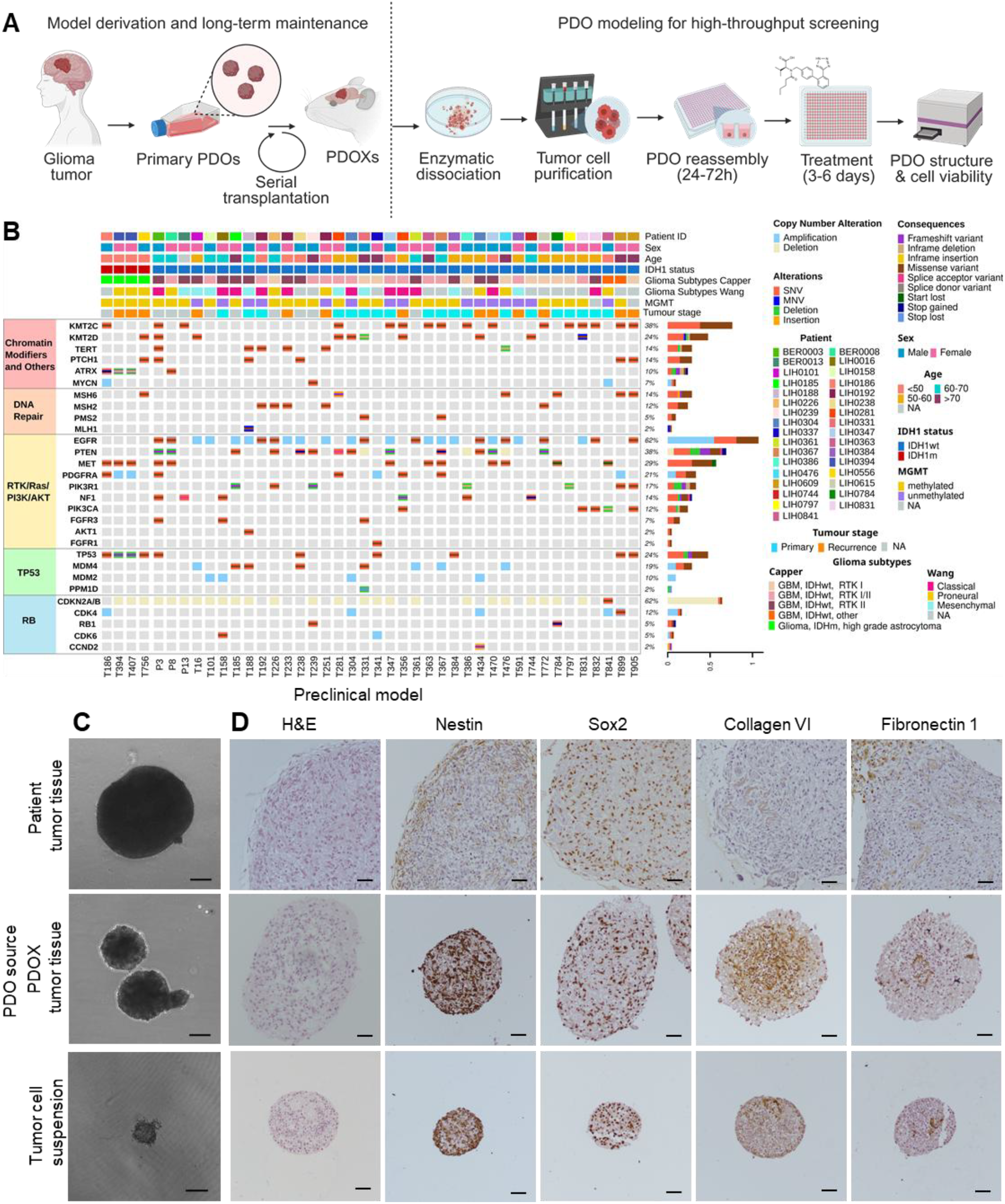
Preclinical cohort of high-grade gliomas for precision medicine studies. **A.** Scheme summarizing a standard protocol of patient-derived organoid (PDO) and patient-derived orthotopic xenograft (PDOX) model derivation and maintenance (left), and uniform PDO establishment for drug screening (right). **B**. Cohort demographics of 42 molecularly characterized preclinical models derived from 33 patients with adult high-grade diffuse gliomas, representing diverse sex, age, tumor stage (primary vs. recurrent tumors) and glioma subtypes (Capper et al.,^27^ Wang et al.,^29^). *MGMT* promoter methylation and *IDH1* mutation status of the models are depicted. See **Table S1** for detailed information; SNV: Single Nucleotide Variant, MNV: Multi Nucleotide Variant, NA: Not Available. **C-D**. Representative images of PDOs derived from patient tumor tissue, PDOX tumor tissue and self-assembled PDOs from PDOX-derived tumor cell suspension; (C) Morphology by bright-field microscopy (magnification: 4x; scale bar: 150μm); (D) IHC (model T1053) showing recapitulation of glioma-specific tumor markers (Nestin, Sox2) and extracellular matrix (Collagen IV, Fibronectin 1); magnification: 10x; scale bar: 50μm.

Epigenetic-based diagnosis assigned the majority of the models to adult-type GBMs with the epigenetic features of the RTK I and II subgroups defined by Capper et al.^27^, although similarly to patient tumors, several models did not show a match for a unique GBM molecular subtype (score ≥0.5, **Table S1**). Lack of DNA methylation-based mesenchymal subtype can be explained by replacement of human tumor microenvironment components upon preclinical modeling^23^, a key contributor to this classifier. Importantly, Malta et al.,^28^ classifier of DNA methylation and Wang et al., classifier^29^ of RNA-seq profiles revealed recapitulation of the tumor-intrinsic subtypes, including proneural, classical and mesenchymal, although mesenchymal subtypes displayed also strong classical-like expression pattern (**Table S1, Fig S1A**). GBM models exhibited diverse molecular alterations known to characterize patient subtypes, such as gene amplifications (e.g., *EGFR, PDGFRA, CDK4, CDK6*), mutations (e.g., *TP53*), and promoter methylation (e.g., *MGMT*, **Fig. 1B)**.

Preclinical modeling of IDH1/2m gliomas is particularly challenging as previously reported by us and several other groups^30,31^, owing to their indolent nature. While short-term PDOs can be derived from IDH1/2m tumors of different grades, the long-term propagation as PDO/PDOX lines is a bottleneck, selecting for the most aggressive tumors. Molecular diagnosis of four IDH1m models revealed features of adult-type high-grade astrocytomas (T186, T394, T407, T756; **Table S1**), which aligned with the molecular classification of the patients’ tumors, regardless differences in the histopathological diagnoses. Three models were derived from recurrent radiotherapy-treated tumors (T394, T407, T756), historically classified as secondary GBMs due to their progression from lower-grade gliomas. *CDKN2A/B* loss classifies them into grade 4 astrocytomas. T394 and T407 represent a unique pair derived from two consecutive surgeries of the same patient, occurring in close temporal proximity and demonstrating consistent molecular features. T186 was derived from recurrent tumor that was not subjected to radiotherapy, histopathologically classified as grade 3 anaplastic oligodendroglioma. While presenting common features of IDH1m gliomas (e.g. *TP53* mutation), models differed at several (epi)genetic modifications. The fifth IDH1m model, T515 was not characterized due to its extremely long growth *in vivo* (survival > 1 year). In summary, our cohort’s diversity in high-grade gliomas provides a strong foundation for evaluating precision medicine approaches.

### Primary PDO cultures are amenable to large-scale drug screening while preserving heterogeneous histological structures

To facilitate high-throughput functional studies, we developed a robust *ex vivo* protocol tailored to primary cultures derived from patient or PDOX tumors (**Fig. 1A**). Our standard protocol for model derivation involves mechanical dissociation of patient and PDOX tumor tissue to generate primary PDOs, maintaining tissue heterogeneity and extracellular matrix integrity (**Fig. 1C-D, Fig. S1B**). However, the resulting variability in PDO sizes hinders scalability and high-throughput screening. To overcome this, we enzymatically digested PDOX-derived tumor tissue, purified human tumor cells, and recreated uniformly sized PDOs (130-150µm) using a 384well plate format and 1000 tumor cells per well, opposed to rather variable size PDOs formed by mechanical dissociation (300-750µm). Tumor cells self-assembled into 3D structures within 24-72h, recapitulating histological complexity **(Fig. 1C-D, Fig. S1B-C)**.

Functional assays optimized for low cell numbers and 3D cultures identified the 3D CellTiter-Glo® luminescent assay as the most reproducible readout, complemented by IncuCyte-based imaging for monitoring PDO size and structure. Despite their primary nature, PDOs exhibited sufficient growth *ex vivo* over 6–9 days, enabling the testing of compounds requiring cell cycle transitions **(Fig. S1D)**. This comprehensive pipeline, integrating patient-derived models with advanced primary PDO culture and screening protocols, provides a robust platform for precision medicine research.

### High-throughput epigenetic drug screening identifies differential vulnerabilities of high-grade gliomas

To investigate the therapeutic vulnerabilities of high-grade gliomas, we performed high-throughput *ex vivo* screening in PDOs using a drug library comprising 202 compounds. The library was enriched for inhibitors targeting epigenetic regulators (n=122), classified into 16 drug families based on their molecular targets (**Fig. S2A, Table S2**), including histone deacetylases (HDACs, n=38), histone methyltransferases (HMTs, n=35), epigenetic reader domains (n=24), alongside kinase inhibitors (n=47), DNA damage/repair enzyme inhibitors (n=28) and a subset of transcription factors (n=5). Each compound was tested at 5µM for 72h and 144h (**Fig. S2B**). Out of 30 models screened, 27 (24 GBMs and 3 grade 4 astrocytomas) met stringent quality control criteria, defined by the coefficient of variation (CoV) ≤20%. The major source of variation in drug sensitivity profiles arose from the diversity across individual patients, followed by *IDH1* status, individual tumor models, time of treatment and tumor stage (**Fig. 2A-B**). Glioma subtypes^27,29^, *MGMT* promoter status, biological (mouse strain) and technical replicates (plate) had a minor impact on the source of variation. These criteria confirmed high assay reproducibility and robust readouts.

**Figure 2.**
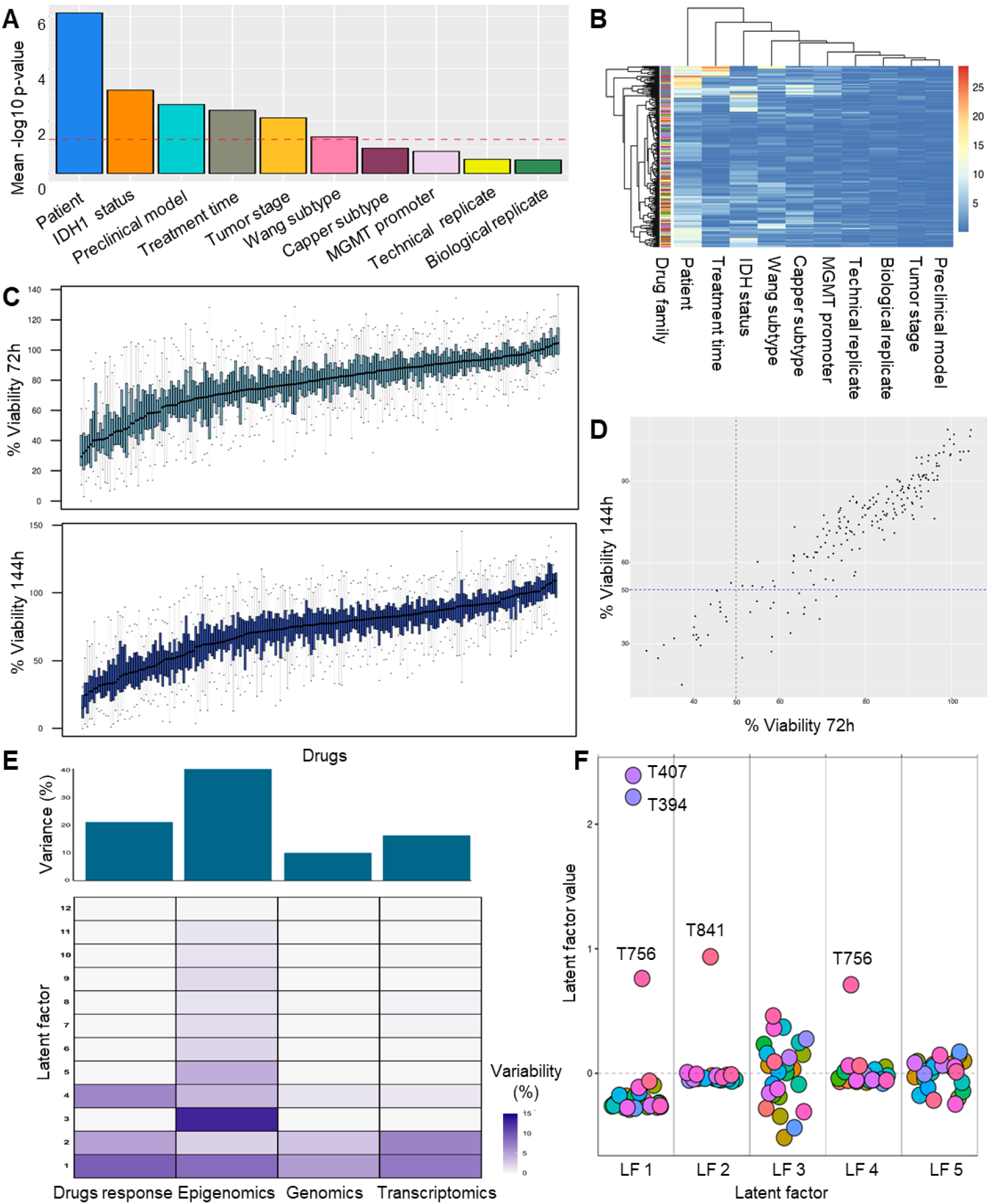
Multi-omics factor analysis of *ex vivo* drug screening in PDOs. **A.** ANOVA showing the mean drug sensitivity p-values associated with ten variables. Technical and biological replicates represent experimental plates and mouse strains respectively. Dashed red line indicates the p-value threshold of 0.05. **B**. Heatmap displays ANOVA p-values (-log10 scale) for each drug, highlighting significant sources of variation across different features. A value of 1.3 corresponds to a significance threshold of p=0.05 **C**. Distribution of % cell viability at two treatment timeponts (72h and 144h). Each box plot displays a single drug’s cell viability across all models assessed (n=27). Boxes represent the interquartile range, with the bottom and top edges indicating the 25th and 75th percentiles, respectively, whiskers represent the full range of data, excluding outliers. The center line within each box denotes the median. **D**. Comparison of drug potency at 72h (x-axis) and 144h (y-axis). Dotted lines indicate the 50% viability threshold. Each dot represents median response across models per drug. **E**. Visualization of the 12 latent factors (LF) identified by MOFA. The heatmap presents the % of variance explained by each individual LF (rows) across different layers. The top panel shows the total % of variance explained by each layer. **F**. LF values across preclinical models. Factor values for the top five latent factors (LF1-LF5) are plotted for each model. Distinct models are color coded.

PDOs displayed variable responses across the cohort, with median cell viability per drug ranging from ∼30-105% for 72h treatment and ∼15-110% for 144h treatment across the models (**Fig. 2C**). The responses at two time points highly correlated to each other with 20 compounds reaching viability <50% at 72h and 144h. Interestingly, extended treatment significantly enhanced the efficacy of several compounds, particularly those targeting epigenetic regulators such as HMTs, HDACs and epigenetic reader domain inhibitors (e.g., AZD2461, Zebularine, RGFP966, CX-6258 HCl, AZD1208; **Fig. 2D, Table S3)**. This observation underscores the advantage of prolonged treatment durations for accurately capturing the therapeutic potential.

Broadly, the most effective compounds were identified as those that reduced cell viability at 144h by ≥50%. None of the compounds showed consistent decrease of viability <50% across all the tested models, while only 4 compounds (TAK-901, UNC0638, UNC0379, Mitomycin C, **Table S3**) showed viability <50% across 90% models. Regrettably, TAK-901, UNC0638 and Mitomycin C have shown limited brain penetration via the blood brain barrier (BBB)^32–34^. While UNC0379’s properties to reach brain parenchyma need further investigation, it shows poor pharmacokinetic properties *in vivo*^35^. 35 compounds that reduced cell viability to <50% in more than half of the tested models belonged to HMT (e.g., MI-503, MI-463, JNJ-64619178), HDAC (e.g., Quisinostat, CUDC-907, Trichostatin A) and kinase (e.g., NVP-BSK805 2HCl (JAK), SGI-1776 (Pim1), TG101209 (FLT3/JAK/c-RET)) drug family inhibitors (**Fig S2C, Table S3**), suggesting inter-patient variability in responses. Similarly, the most effective drugs that showed cell viability <25% in more than half of the tested models were HDAC inhibitors (Mocetinostat (HDAC1), SRT1720 HCl (SIRT1), Dacinostat (general HDACi)). These findings highlight substantial inter-patient variability in drug sensitivity, potentially reflecting distinct molecular dependencies that could be exploited for the precision medicine strategies.

### Unsupervised multi-omics factor analysis reveals high-grade glioma subtypes with differential drug responses

We further leveraged multi-omics profiling to explore whether differential drug responses can be associated with specific molecular features. To identify distinct responder subtypes, we applied MOFA, an unsupervised approach for multi-omics datasets. MOFA identifies major axes of variation, termed latent factors (LFs), which encompass multiple molecular features and enable exploratory analysis across diverse data modalities^26^. We hypothesized that this approach would reveal subgroups characterized by distinct drug response profiles linked to molecular biomarkers.

We selected drug response data at 144h of treatment, given the more pronounced effects observed at this time point **(Fig. 2C-D)**. The drug-response layer was integrated with molecular profiling, incorporating genetic aberrations and copy number aberrations (‘genomics’ layer), genome-wide DNA methylation status (‘epigenomics’ layer) and genome-wide gene expression (‘transcriptomics’ layer). MOFA identified twelve LFs, with the first six factors collectively explaining above 90% of the total variance in the dataset **(Fig. S2D)**. Among these, LF1, LF2, and LF4 were most prominently associated with all three molecular layers as well as the drug-response layer (**Fig. 2E)**, highlighting their relevance in connecting molecular features with functional outcomes. Other LFs were largely associated with the diversity of the DNA methylation profiling, reflecting the strong relevance of DNA methylation to glioma tumor classification. Interestingly, LF1 and LF4 stratified IDH1m models, where LF1 discriminated all three IDH1m astrocytomas (T394, T407, T756) from GBMs, whereas LF4 appeared to be specific for T756 (**Fig. 2F**). LF2 identified T841, a GBM model, showing an unclear molecular classification. Subsequent analyses focused on LF1, LF2, and LF4, to reveal unique drug vulnerabilities and the associated molecular features.

### IDH1m high-grade astrocytomas exhibit distinct drug responses compared to GBMs

To delineate therapeutic vulnerabilities and associated molecular features of IDH1m grade 4 astrocytomas, we focused on latent factors LF1 and LF4, which segregated IDH1m models (T394, T407, and T756) from GBMs. Notably, T756 exhibited reduced variance in LF1 yet demonstrated unique LF4 characteristics, indicative of inter-patient differences within the IDH1m subgroup (**Fig. 2F**). First, we interrogated the drugs with the highest factor loadings, representing compounds that showed unique responses in the associated models (**Fig. 3A, Table S4**). LF1 identified five compounds with loading scores >0.75, including three HMT inhibitors (EPZ011989, CPI-1205, BRD4770) and two HDAC inhibitors (RGFP966, Sirtinol). Assessment of individual responses per preclinical model revealed that, while each compound showed differential responses compared to GBMs, only RGFP966 reached viability <50% in all three IDH1m astrocytomas (**Fig. 3B**). CPI-1205 showed PDO viability <50% in T394 and T407, a response shared rather with T841 GBM PDOs, than T756 IDH1m PDOs **(Fig. 3B)**. Dose-dependent validation in selected models confirmed selective potency of RGFP966 in the three IDH1m PDOs (IC50: T394-3.5µM, T407 3.4µM, T756 2.7µM, **Fig. 3C**) and additional IDH1m glioma stem cell-like cultures (GSCs, **Fig. S3A**) compared to equivalent GBM models. The efficacy of RGFP966 in all IDH1m models assessed suggests a potential broader application, beyond precision medicine strategy. Since RGFP966 (HDAC3 specific inhibitor) is a selective BBB penetrant inhibitor, its potential for clinical application warrants further evaluation.

**Figure 3.**
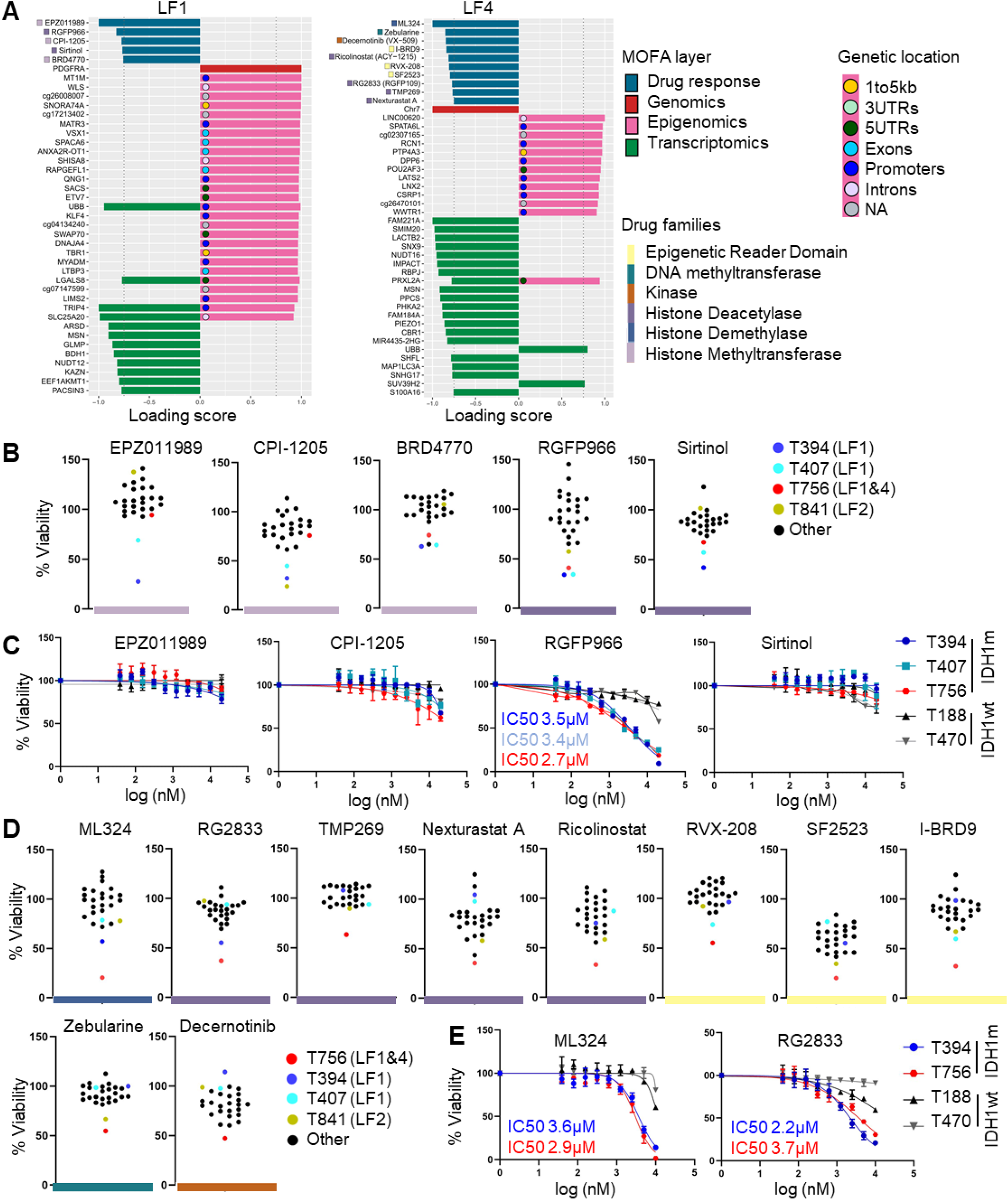
Latent factor analysis reveals differential drug sensitivities in IDH1m high-grade astrocytomas. **A.** The summary of loading scores for the three omics layer and drugs identified for LF1 and LF4, discriminating IDH1m high-grade astrocytomas (threshold >0.75, except for >0.9 for the epigenomics layer). For graphical clarity, only the top 25 epigenomics features are shown. Drugs names are color coded per drug family, genetic locations are specified for the epigenomics probes. **B**. Mean % viability of drugs with loading scores > 0.75 characterizing LF1 across all 27 models. LF1-specific IDH1m models are color coded: T394 dark blue (LF1), T407 light blue (LF1), T756 red (LF1&4). IDH1wt GBM models are presented in black, except for T841 dark yellow (LF2). **C**. Dose-response validation of LF1-identified drugs with mean % viability <50%. RGFP966, a selective HDAC3 inhibitor, demonstrated the highest potency in IDH1m high-grade astrocytoma PDOs, with IC_50_ values below 4µM (mean +/- SD, n=4 per concentration). **D**. Mean % of viability per individual model for ten drugs with LF4 loading score >0.75. LF4-specific T756 model is color coded in red. **E**. Dose-response validation of two LF4-identified drugs, ML324 (JMJD2/KDM4 inhibitor) and RG2833 (HDAC1/3 inhibitor), confirm higher efficacy in IDH1m high-grade astrocytomas (PDOs T756 and T394) with IC_50_ values below 4µM (mean +/- SD, n=6 per concentration), compared to IDH1wt GBM models.

We further interrogated drugs selectively identified in LF4 for T756 (**Fig. 3A**). LF4 analysis revealed ten compounds with loading scores >0.75, including four HDAC inhibitors (RG2833, TMP269, Nexturastat A, and Ricolinostat), two epigenetic reader domain inhibitors (RVX-208, and I-BRD9), a dual epigenetic reader domain and kinase inhibitor SF2523, DNA methyltransferase inhibitor Zebularine, kinase inhibitor Decernotinib, and histone demethylase inhibitor ML324. Among these, six compounds (ML324, RG2833, Nexturastat A, Ricolinostat, SF2523, and I-BRD9) reduced viability below 50% in T756 PDOs, although SF2523 and Nexturastat A showed potency also in other GBM PDOs (**Fig. 3D**). T756 PDOs and GSCs derived thereof confirmed dose-response sensitivity with IC_50_ range of 1.9µM to 6.7µM (**Fig. 3E, Fig. S3B-C)**. However, a similar response was also observed in T394 IDH1m PDOs (2.2µM to 4.5µM) and IDH1m GSCs (1.7µM to 10µM), suggestive of IDH1m-specific responses (**Fig. 3E, Fig. S3B-C)**. While ML324 (JMJD2/KDM4 inhibitor) and RG2833 (HDAC1/3 inhibitor) showed stronger efficacy in other IDH1m models compared to GBMs, four additional inhibitors exhibited similar effects also in GBM PDOs and GSCs, suggesting broader toxicity (**Fig. S3B, Fig. S3C)**. In comparison, Vorasidenib, a clinically approved drug for low-grade IDH1/2m gliomas, was ineffective in high-grade models, aligning with existing data^36^ **(Fig. S3D)**. Overall, MOFA combined with dose-response confirmations support the selection of subgroup- and patient-specific drugs of IDH1m grade 4 astrocytomas. While several drug responses appear specific to individual PDO models, HDAC3 inhibitors appear selectively targeting high-grade IDH1m astrocytomas

### Molecular features associated with differential drug responses in IDH1m high-grade astrocytomas

To elucidate potential biomarkers linked to drug responses in IDH1m astrocytomas, we further interrogated the omics features driving LF1 and LF4. Despite sharing the same *IDH1* mutation status and harboring distinct somatic mutations (**Fig. 1B**), the majority of genetic alterations in the three IDH1m models had minimal impact on LF1 and LF4, as evidenced by low factor loadings (<0.75; **Fig. 3A, Table S4**). MOFA identified *PDGFRA* amplification as the primary driver of LF1 genomic layer. This amplification was exclusive to T394/T407 and absent in T756 model (**Fig. 1B**), explaining overall lower factor loading value of T756 in LF1. LF4 separation was attributed to the absence of gains within chromosome 7 in T756 (**Fig. 3A**).

At the transcriptomic level, MOFA analysis identified 12 gene transcripts with high LF1 loading scores **(Fig. 3A)**. All genes presented lower expression levels in IDH1m models **(Fig. 4A**). Four genes (*SLC25A20, MSN, EEF1AKMT1, LGALS8)* showed similar low expression levels in three IDH1m models, whereas others were differentially expressed between T394/T407 and T756, further highlighting inter-patient differences. Similarly, out of 21 gene transcripts with high loading scores in LF4, 19 were expressed at lower levels in T756 compared to other models, *UBB* showed higher levels compared to T394/T407, equivalent to GBMs, whereas *SUV39H2* was highest among tested models **(Fig. 4A)**. Analysis of TCGA cohort revealed that the majority of the identified genes showed decreased levels in IDH1/2m grade 4 astrocytomas compared to GBMs, further highlighting their broader relevance (**Fig. S4**). Interestingly, higher expression of *SUV39H2*, a H3K9 HMT, was further confirmed in a subset of IDH1/2m grade 4 astrocytomas. We speculate that higher SUV39H2 could potentially create a greater dependency of IDH1m tumors to epigenetic drugs, by further adding to the effect of 2-HG-inhibited H3K9 demethylases, since both enzymes contribute to the suppressed expression of differentiation factors and tumor suppressors. A differential gene expression analysis of the RNA-seq revealed that IDH1m models showed higher levels of genes associated with chromatin organization and DNA binding, which were particularly pronounced in T756 (**Fig. 4B, Table S5**).

**Figure 4.**
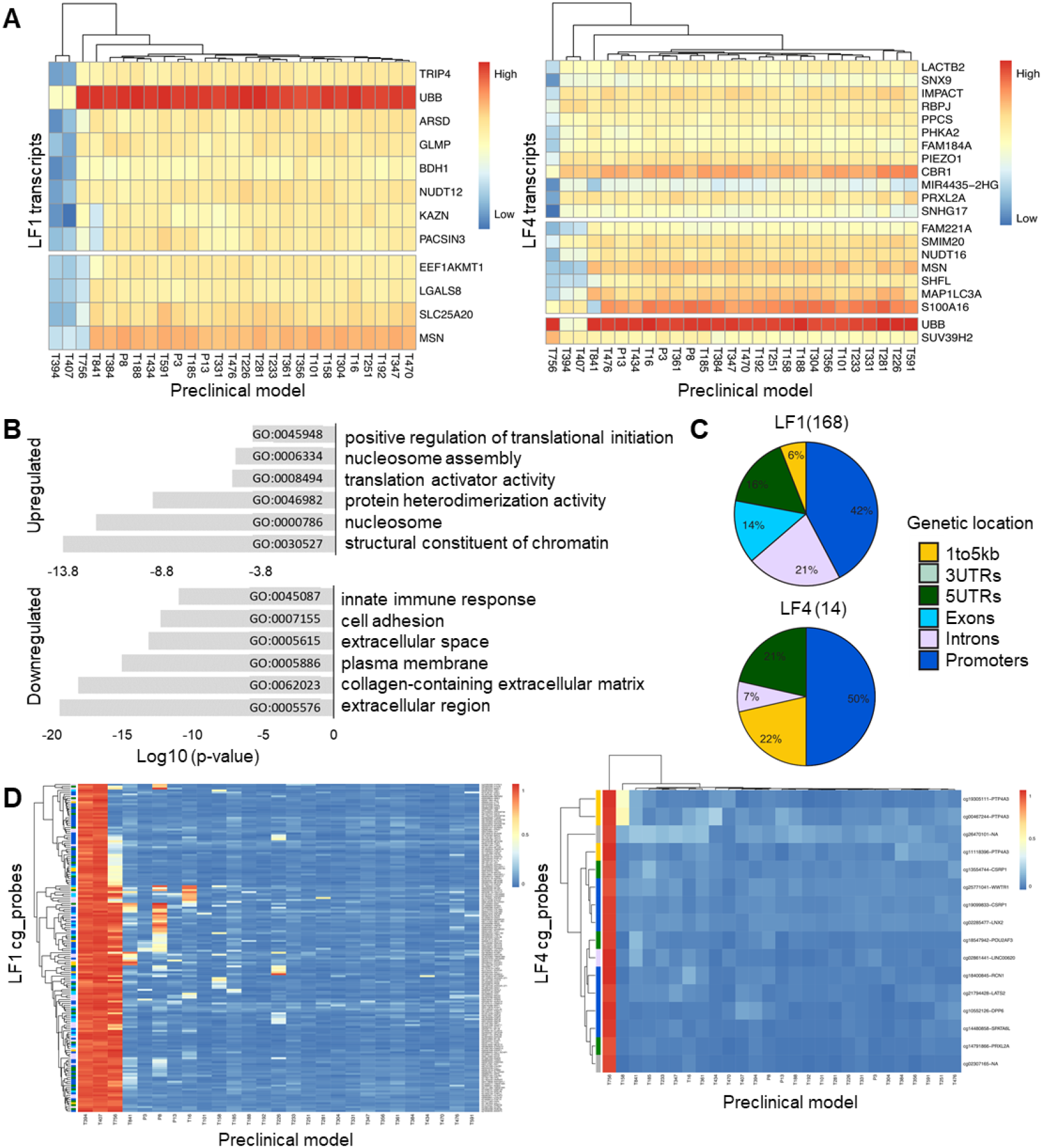
MOFA-driven molecular features discriminating IDH1m high-grade astrocytomas. **A.** Heatmaps showing expression levels of top genes with loading scores > 0.75 identified from LF1 and LF4 across the preclinical cohort. **B**. Differential gene expression analysis between IDH1m astrocytoma vs. IDH1wt GBM models (DEGs: FDR ≤0.01 and |log2FC|≥1), confirming enrichment of genes associated with chromatin organization and DNA binding. Top six GO terms are displayed characterising up- and downregulated genes. **C**. Epigenetic features discriminating IDH1m astrocytoma and GBM models. Distribution of LF1 and LF4-associated DNA methylation cp probes are shown across genetic locations. **D**. Heatmaps depicting top epigenetic features identified in LF1 and LF4, showing differential methylation patterns in IDH1m models. β-values per each DNA methylation probe and associated gene are depicted.

DNA methylation classifier diagnosed our IDH1m models as high-grade astrocytomas, with G-CIMP low phenotype, further confirming their high aggressiveness^28^ (**Table S1**). LF1 further discriminated 191 DNA methylation probes with loading scores >0.9, where 168 probes associated with specific genes **(Fig. 3A, Table S4)**. All probes showed higher methylation in IDH1m models: 126 probes exhibited high methylation in the three IDH1m models, whereas 42 probes showed marked methylation in the T394/T407 only (**Fig. 4C-D**). LF4 distinguished the hypermethylation of 16 probes in T756 uniquely **(Fig. 3A, Fig. 4C-D, Table S4)**. A cross-omics comparison of the transcriptomic and epigenomic features identified four common genes in LF1: *TRIP4, UBB, LGALS8*, and *SLC25A20* and one gene in LF4 (*PRXL2A*), showing probes with increased DNA methylation and lower transcript levels. *SLC25A20* exhibited high DNA methylation and low expression across all three models, whereas *TRIP4, UBB*, and *LGALS8* showed high DNA methylation and low expression in T394/T407 only, and *PRXL2A* showed high methylation and low expression in T756 **(Fig. 3A, Fig. 4A**,**D)**. Out of those, *TRIP4, LGALS8*, and *SLC25A20* show significantly lower levels in IDH1/2m astrocytomas compared to GBMs (**Fig. S4**), while *UBB* and *PRXL2A* did not show statistically significant differences, suggesting that these expression differences in our models are patient-specific.

In summary, MOFA revealed genes characterized by lower gene expression and higher DNA methylation as key features associated with drug responses in IDH1m high-grade astrocytomas, complicating biomarker discovery but highlighting patient-specific molecular profiles.

### A unique drug vulnerability of a MYCN-amplified GBM

Among 24 GBM models tested, only T841 showed unique features discriminated by LF2 (**Fig. 2F**). Although four compounds were identified by LF2 with loading scores >0.75, only two were associated with <50% viability compared to other models (**Fig. 5A-B**). HMT inhibitors EED226 and CPI-1205, demonstrated high efficacy. EED226 was explicitly effective in T841, while CPI-1205 also exhibited activity in T394/T407 IDH1m PDOs, aligning with previous findings by LF1. Dose response assay validated both EED226 and CPI-1205 as potent compounds with IC50 of 5.6µM and 5.8µM **(Fig. 5C)**.

**Figure 5.**
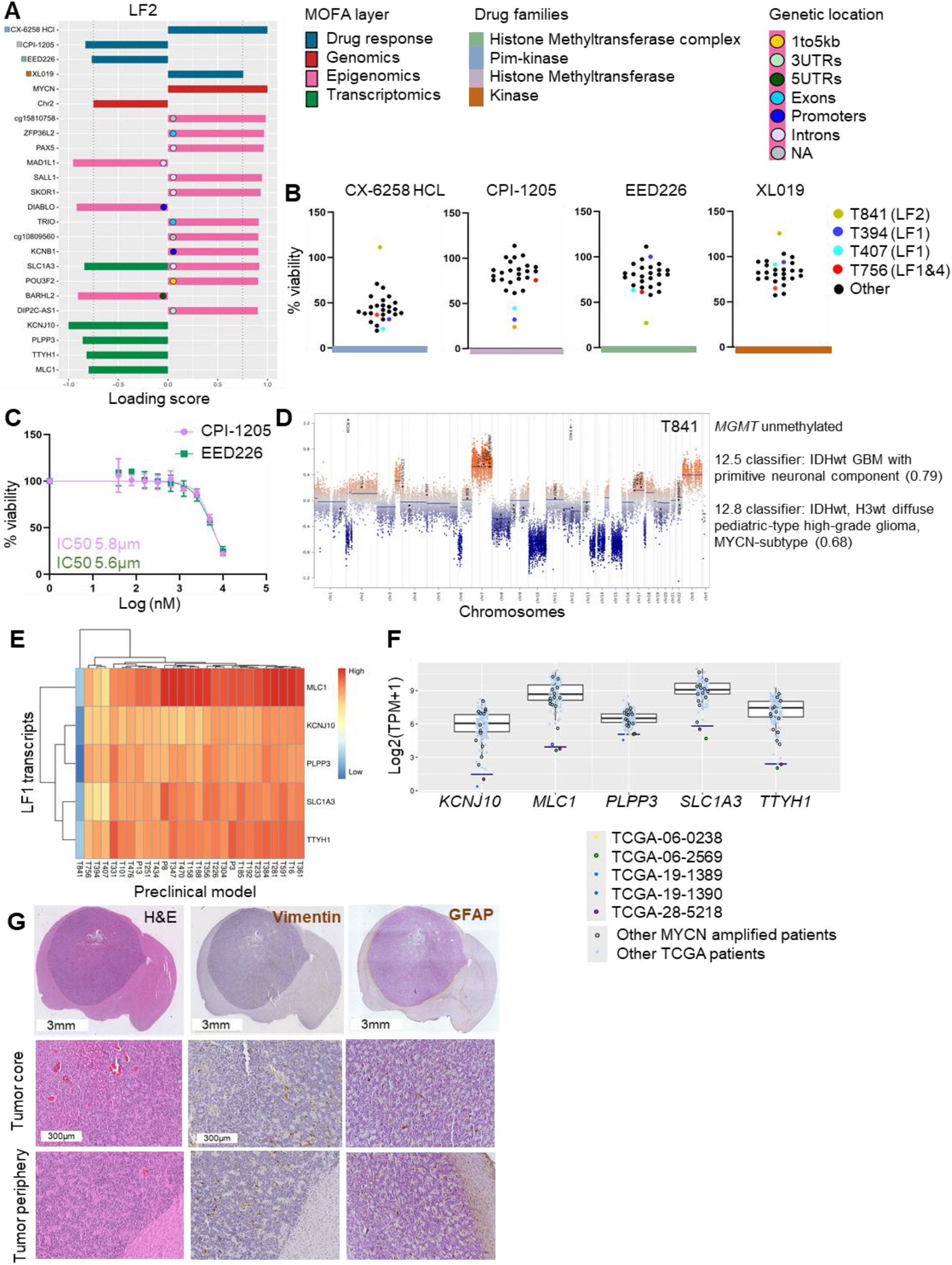
Distinct omics signatures and drug responses for a *MYCN*-amplified GBM model. **A.** The summary of loading scores for the three omics layer and drugs identified for LF2, discriminating T841 model (threshold >0.75, except for >0.9 for the epigenomics layer). For graphical clarity, only the top 25 epigenomics features are shown. Drugs names are color coded per drug family, genetic locations are specified for the epigenomics probes. **B**. Mean % viability of drugs with loading scores >0.75 discriminating LF2-identified T841 model (dark yellow) across all 27 models. **C**. Dose-response validation of LF2-associated drugs in T841 PDOs (mean +/- SD, n=4 per concentration). **D**. DNA methylation-derived CNV profile of T841 model showing *MYCN* and *CDK4* amplification. IDH1/2wt GBM-characteristic +7/-10 Chr feature is present. 12.5v and 12.8v Heidelberg classifier scores show no clear match to molecular glioma subtype. **E**. Expression levels of genes identified by MOFA from LF2 across the preclinical cohort. **F**. Box plots displaying expression levels of the five LF2-identified genes in GBM patient tumors (TCGA cohort, n= 141). Blue line indicates the 1st percentile threshold. Patients are color-coded. Patients with *MYCN* amplification are marked with a black outline around data points. **G**. Representative H&E, Vimentin and GFAP staining in PDOX tumors *in vivo*, showing pseudo-rosette structures, limited tumor cell diffuse growth and low/negative expression of Vimentin and GFAP in tumor cells. Scale bars 3mm for coronal brain sections and 300µm for magnified areas.

The distinct molecular profile of T841 was driven by features across LF2 genetic, transcriptomic, and epigenetic layers. Genetically, MOFA identified *MYCN* amplification, a rare alteration in GBM, accompanied by a partial chromosome 2 loss shared with T356 (**Fig. 5A, D**), which was also confirmed in the original patient tumor (**Fig. S5A**). Additional rare features included *CDK4* amplification and *CDKN2A* mutation (**Table S1**). Transcriptomic layer identified five genes with LF2 loading scores >0.75: *KCNJ10, PLPP3, SLC1A3, TTYH1*, and *MLC1*, showing low expression levels in T841 (**Fig. 5A, E**). These genes are primarily involved in ion and membrane transport processes. Additional analysis of differential genes further indicated lower levels of genes associated cadherin/fibronectin-associated extracellular matrix and NOTCH3-associated nervous system development signaling, while upregulated genes were linked to vesicle trafficking and fusion (**Fig S5B, Table S5**). Interestingly, analysis of the TCGA patient cohort confirmed presence of rare GBMs showing similar low expression patterns (**Fig. S5C**). An outlier analysis of the GBMs, using the first percentile, identified five tumors resembling T841, out of which two displayed *MYCN* amplification. Out of 15 *MYCN*-amplified GBMs in total, TCGA-28-5218 showed markedly low expression in 4/5 LF2 genes (*KCNJ10, MLC1, SLC1A3, TTYH1*), while TCGA-06-2569 was an outlier for *MLC1, SLC1A3, TTYH1*. While additional *MYCN*-amplified tumors also showed lower expression levels for these genes, such low levels were not detectable in all *MYCN*-amplified GBMs, suggesting additional inter-patient differences (**Fig. 5F**).

Epigenetic profiling identified 16 methylated probes with LF2 loading factors >0.9, including 4 hypomethylated and 12 hypermethylated probes linked to 12 genes (**Fig. S5D)**. Only two probes were associated with promoter regions: *cg14094409* (hypomethylated, *DIABLO*) and *cg01663603* (hypermethylated, *KCNB1*), and these differences were not translated to differential expression. Interestingly, DNA methylation-based classification using v12.5 suggested a GBM, IDH1/2wt profile with a primitive neuronal component (score 0.8, **Fig. 5D**), a poorly differentiated phenotype with frequent absence of GFAP expression. However, further analysis with v12.8 indicated a different classification with similarity to pediatric-type diffuse high-grade glioma, H3wt, IDH1/2wt, MYCN subtype (score 0.65). Molecular diagnosis of the patient’s tumor showed ambiguous classification, with features of both GBM and pediatric-type high-grade glioma (scores 0.24–0.26) (**Fig. S5A**). Additional subtyping based on Malta et al.,^28^ also revealed unclassical LGm6-GBM subtype (**Table S1**). The tumor’s temporal lobe localization and histopathological features, including pseudo-rosette-like growth patterns and limited cell diffusion in PDOXs (**Fig. 5G, Fig. S5D**), aligned with GBMs harboring primitive neuronal components. This was confirmed by a lack of pronounced expression for GFAP, Vimentin and Nestin, usually observed in GBM PDOXs^15,37,38^. In summary, we identified a rare *MYCN*-amplified GBM with unique molecular profiles and drug responses.

## Discussion

Understanding the intricate interplay between various molecular layers driving cancer progression and treatment response is crucial for developing personalized therapies. A lack of robust clinically-relevant models is regularly listed as one of the main challenges in establishing successful precision medicine in brain tumors, including high-grade gliomas^18^. Therefore, development of diverse cohorts of well characterized patient-derived models combined with robust functional readouts are of greatest importance to test targeted therapies prior to clinical trials. Our preclinical models stand out as a unique platform allowing for experimental protocols *ex vivo* in PDOs and *in vivo* in PDOXs. Unlike cohorts based on long-term *in vitro* cultures, our models preserve a wide range of patient-specific molecular features. Notably, our PDO/PDOX models retain key characteristics, including tumor cell ploidy, extrachromosomal gene amplifications, and phenotypic heterogeneity^15,21,22^. Previous studies, including ours, have shown that GSCs may undergo genetic evolution *in vitro*, often losing critical features for targeted therapies, such as EGFR amplifications^39–41^. While our cohort is enriched in GBMs, we also derived unique IDH1m models, representing molecularly high-grade IDH1m astrocytomas. Such bias is also visible in GSC cohorts, shown to select for the most aggressive clones^42,43^. Modeling IDH1/2m low-grade gliomas will require alternative protocols, such as assembloids with healthy brain organoids or genetic engineering^44,45^.

To enable robust functional readout, we adapted the PDO derivation protocol for high-throughput functional studies. By reconstructing uniform PDOs of consistent size, we avoided the variability and labor-intensive histology-based readouts^46–48^ or analytically-challenging high-content imaging applied^49,50^ in other protocols. This approach aligns with broader trends in organoid research toward standardization and scalability. Notably, primary glioma cells self-assemble into heterogeneous PDO structures without requiring additional extracellular matrix support, a feature that underscores their ability to recapitulate *in vivo* tumor architecture^51^. In contrast, GSCs often form loose, non-structured 3D spheres, highlighting the superior fidelity of PDOs. Our current protocol depletes tumor microenvironment components to focus on tumor-intrinsic responses. While we acknowledge that presence of supporting microenvironmental cells can influence the responses to treatment, this remains a general challenge also noted in broader organoid research^51^. Reproducible studies incorporating microenvironmental components will require advanced co-culture approach in the future^46,52^. Combining microscopic visualization of PDOs prior and after treatment coupled with sensitive viability readouts has significantly enhanced the reproducibility of our protocol. To our knowledge, it is the first report on high-throughput drug screening in brain tumor PDOs. This methodological refinement is consistent with advancements in organoid-based drug screening, where highthroughput and high-content imaging techniques are becoming standard^7^.

The drug repurposing paradigm has gained increasing interest within the cancer research community in recent years. Although diverse medium and large libraries of approved drugs are available for testing, neuro-oncology community is challenged with the BBB penetration and neurotoxicity. Indeed, our screen revealed several potent compounds, though with limited translational potential due to limited BBB penetration. Since the majority of drugs showed responses dominated by inter-patient differences, we applied precision medicine approach to reveal personalized vulnerabilities. MOFA, a statistically robust generalization of principal component analysis designed to integrate multiple data modalities into a unified framework^26^, allowed us to combine molecular layers with cellular responses to drugs. Traditional statistical methods used in drug testing like ANOVA or linear/logistic regression often struggle with the complexity of multi-omics datasets, while MOFA can effectively integrate multiple data modalities across molecular and functional dimensions and merges different patterns of missing data to uncover tumor subgroups with unique profiles. Importantly, MOFA enabled unbiased approach without requiring predefined drug response categories or molecular subgroups, offering greater flexibility than supervised methods^53,54^. Here, by applying 27 models, three molecular layers and 202 drugs, MOFA revealed three latent factors linking all molecular layers to drug responses. Additional diversity was defined uniquely in the DNA methylation profiles, corroborating importance of this molecular layer in defining subtypes of high-grade gliomas^27^. Interestingly, this approach revealed distinct vulnerabilities of IDH1m high-grade astrocytomas to several epigenetic inhibitors. Among GBMs, only one model stood out as unique. We have previously shown that diverse status of *EGFR* mutations and structural variants has limited impact on responses to EGFR inhibitors^15^, which was further confirmed by MOFA. Assessment of vulnerabilities to other key modifications, such as *PDGFRA, CDK4/6* and *MDM2/4* amplifications may require additional broader drug library. We acknowledge that MOFA may require higher number of diverse models and larger drug library to reveal additional subgroups, as initially employed in 200 chronic lymphocytic leukemia^26^ and more recently in 335 breast cancer patient models^55^ and 170 Alzheimer disease patients^56^. By advancing drug libraries and diversity of the models, integrative multi-modal approaches could reveal individual drug vulnerabilities or patient subgroups.

Epigenetic mechanisms play a critical role in glioma tumorigenesis and therapy resistance. Our approach revealed patient-specific inhibitory effects of multiple drugs on glioma cell viability. For instance, HDAC inhibitors demonstrated selective efficacy against IDH1m models, with the HDAC3 inhibitors RGFP966 and RG2833 eliciting robust responses in all evaluated IDH1m PDOs and GSCs. These findings align with prior reports highlighting the potent effects of HDAC inhibitors on IDH1m glioma cells^57–61^. Although IC_50_ values were in the micromolar range, further investigations are warranted given the limited efficacy of IDH1 inhibitors in hi ghgrade gliomas^12^. While IDH1 mutation-specific peptide vaccines, (NOA-16, NOA-21 clinical trials^62,63^) show promising results in high-grade astrocytomas, long-term efficacy could be potentiated by combining such immunotherapies with tumor-intrinsic modulators. While MOFA revealed several molecular features linked to these drug responses, the majority appeared as downregulated genes. *SUV39H2* showing particularly high levels in a subset of grade 4 astrocytomas, appeared as an exception, with a potential for a biomarker of responses to inhibitors of epigenetic regulators.

Additional studies are required to confirm whether distinct molecularly-defined GBM subgroups suitable for precision medicine truly exist. In our study, the epigenetic drug library did not identify unique responses linked to molecular profiles and GBM subtypes. For biomarker-negative GBMs, future investigations may need to explore rational drug combinations, particularly epigenetic modulators paired with pathway-specific inhibitors, to overcome intratumoral heterogeneity and broaden therapeutic options. The T841 model showed a particularly rare molecular profile, defined by *MYCN* amplification and DNA methylation features between GBM and pediatric-type diffuse high-grade gliomas. This *MYCN*-amplified GBM patient shares characteristics with a rare subset of gliomas described in the literature^13,64^. Suwala et al. identified GBMs with primitive neuronal components as a distinct subgroup characterized by *MYCN* amplification, frequent inactivation of *TP53, PTEN*, and *RB1*, and a unique methylation profile overlapping with pediatric-type high-grade gliomas. While our analysis did not identify *TP53, PTEN*, or *RB1* inactivation, the shared *MYCN* amplification and epigenetic resemblance to pediatric-type gliomas suggest T841 aligns with this subgroup. Similarly, Korshunov et al. reported *MYCN*- amplified IDH1wt gliomas with aggressive behavior and primitive neuronal features, although in younger patients, further supporting the molecular and histopathological overlap with T841. Our models strongly resemble rare GBMs identified by Perry et al. showing pseudo-rosette growth pattern, characteristic for pediatric tumors such as CNS embryonal tumors. The identification of rare T841-like profiles in TCGA patients^65^, particularly those with *MYCN* amplification and *KCNJ10*/*TTYH1* outlier expression, reinforces the existence of this rare subgroup within adult GBM, highlighting potential therapeutic vulnerabilities to HMT inhibitors like EED226, a selective inhibitor of the Polycomb Repressive Complex 2 (PRC2)^66^. While ED226 has not been extensively studied for GBM, its mechanism targeting PRC2, implicated in GBM progression, suggests potential relevance^67^.

In summary, we show the suitability of molecularly characterized PDO models as robust platforms for identifying therapeutic vulnerabilities in high-grade gliomas. While personalized therapy for high-grade gliomas has yet to demonstrate a survival benefit^68^, this study demonstrates the feasibility of identifying potential biomarkers and therapeutic responses based on patients’ unique molecular profiles. Enlarging functional profiling to a larger PDO cohort and broader drug library may provide additional subtypes of high-grade gliomas for precision medicine. A limitation of our study is the lack of investigation into the underlying molecular mechanisms driving the observed drug sensitivities. While these aspects are crucial for developing effective clinical strategies, they fall beyond the scope of this initial exploratory study.

## Acknowledgements

We are grateful to the Neurosurgery Department of the Centre Hospitalier de Luxembourg, Laboratoire National de Santé and Clinical and Epidemiological Investigation Center of the LIH for support in tumor collection. We thank Virginie Baus, Coralie Pulido and Claire Dording for technical assistance. We acknowledge the support from LIH’s core facilities (Animal facility, In vivo imaging facility, LUXGEN Genome Center).

## Author contributions

Conceptualization; LE, AL, A-CH, SPN, AG; Methodology: LE, AL, A-CH, BN, RT, AO, TMM, SPN, AG; Investigation: AL, A-CH, IK, EK, AO, LH; Formal analysis: LE, AL, A-CH, IK, BN, RT, SF, KBMF, MM, TMM, PVN, SPN, AG; Resources: CH-M, KBMF, MM, GB, FH, SPN, AG; Supervision: PVN, SPN, AG; Writing - Original Draft: LE, AL, AG, Writing - Review & Editing: all authors

## Data availability

Molecular information on PDO/PDOX models is available via the PDMC Finder (https://www.cancermodels.org/about/providers/lih).

## Materials and Methods

### Clinical samples and preclinical model derivation

High-grade glioma tissues were collected from the Centre Hospitalier de Luxembourg (CHL, Luxembourg, Luxembourg) or the Haukeland University Hospital (Bergen, Norway). All patients gave informed consent and tumor collection was approved by the local research ethics committees (National Committee for Ethics in Research (CNER) Luxembourg: protocols 200603/07, 200803/12, 201201/06, www.precision-pdx.lu; Haukeland University Hospital: 2009/117). The research protocols conformed to the principles of the Helsinki Declaration.

PDOs were derived based on previously established protocol from mechanically digested tumor tissue^15,19^. Shortly, tissue fragments were cultured for up to 2 weeks in non-adherent conditions at 37°C under 5% CO2 and atmospheric oxygen in DMEM medium, 10% FBS, 2 mM L-Glutamine, 0.4 mM NEAA, and 100 U/ml Pen–Strep (all from Lonza). PDOX derivation and maintenance was mediated by serial transplantation of primary PDOs in NSG (NOD. Cg-Prkdc^scid^ Il2rg^tm1WjI^/SzJ, 6 PDOs/brain). See **Table S1** for preclinical cohort characterization. For the larger amplification of tumor material for ex vivo experiments, PDOXs were derived in female nude mice (athymic nude mice, Charles River Laboratories, France). Animals were sacrificed at the endpoint defined in the scoresheet and tumor growth evaluation by MRI. Animals were housed in a specific pathogen-free (SPF) facility under a controlled environment (temperature, humidity, and light) with free access to water and food. Animal experiments were performed in accordance with the regulations of the European Directive on animal experimentation (2010/63/EU) and were approved by the Animal Welfare Structure of the Luxembourg Institute of Health and by the Luxembourg Ministries of Agriculture and of Health (LRNO-2014-01, LUPA2019/93, LUPA2024/10).

### Patient-derived glioma stem-like cell cultures (GSCs)

Human GSCs BG5 were kindly provided by Prof. Rolf Bjerkvig from University of Bergen. GSCs, NCH551b (RRID: CVCL_A5HU), NCH612 (RRID: CVCL_X913), NCH1681 (RRID: CVCL_A5HV) and NCH660h, were derived by Prof. Christel Herold-Mende at the University of Heidelberg. GSCs T394NS, T407NS, T756NS, P3NS and P13NS were generated from PDOX models as previously described^15^. GSCs were cultured as 3D spheres in serum-free DMEM-F12 medium (Lonza) containing 1 x BIT100 (Provitro), 2 mM L-Glutamine, 30 U/ml Pen-Strep, 1 U/ml Heparin (Sigma), 20 ng/ml bFGF (Miltenyi, 130-093-841) and 20 ng/ml EGF (Provitro, 1325950500) (NCH601, NCH421k, NCH644) or serum-free Neurobasal medium (Thermo Fisher, Cat# 21103049) supplemented with 1X B-27™ Supplement (Thermo Fisher, Cat# 17504044), 2 mM UltraGlutamine (LONZA, Cat#BE17-605E/U1), 0,1 U/mL heparin (Sigma-Aldrich, Cat# H3149), 20 ng/mL bFGF (Miltenyi, Cat# 130-093-839), 20 ng/mL EGF (Provitro, Cat# 1325960500) and 100 U/mL penicillin‒streptomycin (remaining GSCs).

### *Ex vivo* drug screening and dose response validation

PDOX tumors were dissociated using the MACS Neural Dissociation Kit (Miltenyi Biotec) according to the manufacturer’s instructions. Mouse-derived microenvironmental cells were depleted with the Mouse Cell Depletion Kit (Miltenyi Biotec). Purified tumor cells were seeded into 384-well plates (1000/well, PrimeSurface, S-Bio) and cultured for 72h with agitation to allow PDO formation prior to inhibitor treatment. PDOs were maintained at 37°C with 5% CO_2_ and atmospheric oxygen in DMEM medium supplemented with 10% FBS, 2 mM L-glutamine, 0.4 mM NEAA, and 100 U/mL penicillin-streptomycin (all from Lonza). Drug screening included epigenetic compound library from SelleckChem (Cat. L1900, 182 compounds except for S7951) and 21 additional drugs (**Table S2**). Each compound was applied at a concentration of 5µM. To mitigate experimental variability, each experimental plate included a minimum of 38 control PDOs. This setup constituted one technical replicate in 384 well plate. Drug screening was performed in triplicates using tumor cells derived from at least two independent mouse implantations. For validation studies, PDOs (as above) or GSC 3D spheres (500–1,000 cells per well) were subjected to a two-fold, ten-point serial dilution series ranging from 40nM to 20μM. For each compound, 3–5 replicates per concentration were included, and treatments were repeated in 2–3 independent experiments. Each experimental plate included ≥20 control wells.

PDO and GSC sphere integrity was evaluated prior and after treatment, or in real time during the treatment with IncuCyte S3. Viability was assessed after 3 days (72h) and 6 days (144h) of treatment using the CellTiter-Glo® 3D Cell Viability Assay (Promega) according to the manufacturer’s protocol. Luminescence was measured with a ClarioStar plate reader (BMG Labtech). Relative cell viability for each compound was calculated by normalizing the luminescence values to the 0.1% DMSO controls on each plate. Dose-response curves were generated using the best-fit model, and IC_50_ values were calculated using the variable slope (four parameters) function of the “log[inhibitor] versus normalized response” equation in GraphPad Prism (v9.1.2).

### Immunohistochemistry

PDOs were carefully transferred, washed with ice-cold 1X PBS (Thermo Fisher, Cat# 14190144) and fixed in 4% paraformaldehyde for 20-30 min at RT. PDOs were resuspended in 2.5% agar diluted in water and transferred into Tissue-Tek cryomolds (Sakura, Cat# 94-4565) and kept on ice for 5-10 min until agar solidification, followed by automated paraffin embedding. H&E and IHC were carried out using 3μm paraffin sections following standard procedures. Prior to immunostaining with a Dako Omnis automated staining module (Agilent Technologies), sections were heated at 60°C for 30 min in either low (pH6.1) or high pH (pH9) buffer. For primary and secondary antibodies, see **Table S6**. Signal was developed with the EnVision™ FLEX detection DAB+ Substrate Chromogen System (Agilent Technologies). Pictures were acquired with Nikon microscope Eclipse E600 using Nikon imaging software NIS-Elements version 5.30.01.

### DNA isolation

DNA from fresh or flash frozen samples was extracted using the DNeasy Blood & Tissue Kit (Qiagen). DNA was eluted in 50 μl of nuclease-free water and concentrations were measured using either a NanoDrop 1000 (Thermo Fisher Scientific) or a Qubit 4.0 fluorometer (Thermo Fisher Scientific) together with the Qubit dsDNA BR Assay kit (Thermo Fisher Scientific).

### Targeted sequencing and analysis

190 ng of extracted genomic DNA was enzymatically fragmented and used as a template to prepare a custom Agilent SureSelectXT Target Enrichment Library (Cat No. G9612B) according to the manufacturer’s instructions as described in^15,23^. The library was then sequenced using Illumina Paired-End Multiplexed Sequencing on a MiSeq® and NextSeq500 instrument (Illumina). Bioinformatics analysis of sequencing reads was performed using a Snakemake-based workflow in accordance with GATK best practices as described before^23^. Genomic alterations were visualised with ComplexHeatmap^69^ within R version 4.3.3.

### DNA methylation

DNA methylation profiling was performed using the Infinium® MethylationEPIC BeadChip arrays v2.0 (Illumina) as previously described^15^. Bisulfite conversion of DNA was conducted according to the array manufacturer’s protocol. Intensity Data (IDAT) files from EPIC arrays were processed, quality-controlled, and normalized using the RnBeads package from Bioconductor^70^. The resulting normalized probe intensities were then used to calculate β-values, which were subsequently used for downstream analyses. For glioma classification, DNA methylation profiles were referenced against the comprehensive dataset of over 2,800 neuropathological tumors available at molecularneuropathology.org/mnp (v11b4, and v12.5) and the Heidelberg Epignostix platform (https://app.epignostix.com/, v12.8)^27^.

### RNA-seq

Total RNA was isolated from cryopreserved PDOX tumor tissue using direct-zol RNA miniprep kit (Zymo Research, Cat# R2050). RNA quality was checked with the HS RNA (15 nt) Kit (Agilent, Cat# DNF-472). mRNA-seq profiling was carried out with 50 million reads/sample, 2×150bp, using the Illumina Stranded mRNA prep kit on an Illumina NovaSeq 6000 sequencer. RNA-seq quality control was performed using FastQC. Adaptors were trimmed from raw sequencing reads using Cutadapt version 3.7^71^, and the trimmed reads were aligned to both the human reference genome GRCh38 and the mouse reference genome GRCm39 by STAR 2.7.10b aligner^72^. Deconvolution of mouse and human sequencing reads was performed with XenofilterR^73^. Uniquely human-mapped reads were ultimately used to quantify gene expression using FeatureCounts^74^. The R package DGEobj.utils was used to normalise gene expression, controlling for both library size and the gene lengths, by converting raw counts to TPM (transcripts per million). Heatmap was generated using the R package Pheatmap. All analyses were performed with R version 4.3.3. GBM molecular subtyping was based on the gene signatures from Wang et al.,^29^ and the GSVA package^75^. The DESeq2 package in R was used to identify differentially expressed genes. Genes with FDR ≤0.01 and |log2FC|≥1 were included in downstream analyses. Pathway analysis was done using the DAVID Bioinformatics Resources 2021^76,77^.

### Multi-Omics Factor Analysis (MOFA)

We employed the MOFA2 package (version 1.12.1) in R, utilizing default parameters^26^. MOFA infers latent factors (LFs) that are shared across different omics datasets. We allowed the MOFA model to prune inactive latent factors that failed to capture significant variance and to automatically determine the optimal number of latent factors capable of capturing significant variation. The mean percentage viability from three replicates for each drug served as input for the drug-response layer. The epigenomics layer utilized genome-wide DNA methylation status (beta values from 324,858 probes), while the transcriptomics layer incorporated genome-wide gene expression (TPM values for 35,394 expressed genes). The genomic layer used distinct numerical codes for different variant classes: deletions, insertions, multi-nucleotide variants (MNVs), and single nucleotide variants (SNVs), with ‘zero’ signifying no mutation. The genomic layer used distinct numerical codes represented variant classes, including deletions, insertions, multi-nucleotide variants (MNVs), and single nucleotide variants (SNVs), with ‘zero’ indicating no mutation. Each variant class was further categorized numerically based on its mutation consequence, including frameshift, in-frame deletion/insertion, missense, splice acceptor/donor, start/stop lost, and stop gain variants. Copy numbers were also included as the precise copy number for each instance. Significant features were chosen using a threshold for loading factors (range |0-1|), which is a metric for feature importance. A threshold of 0.75 was applied to each omics data type. However, for epigenomics, a stricter threshold of 0.9 was used to select a smaller, more highly important set of features.

### TCGA LGG-GBM cohort analysis

RNA-seq data and corresponding metadata for 677 TCGA GBM-LGG patients^78^ were downloaded from the cBioportal repository^79^. The dataset was filtered to include only patients with complete information on IDH1 and IDH2 mutation status, 1p/19q codeletion status, and histological diagnosis. This filtering step and WHO2021-based reannotation resulted in a final cohort of 564 patients: 190 patients with IDH1/2m astrocytomas (Grades 2, 3, and 4), 147 patients with IDH1/2m oligodendrogliomas (Grades 2 and 3), 86 patients with IDH1/2wt gliomas (Grades 2 and 3), and 141 patients with IDH1/2wt GBM (Grade 4). For the outlier analysis, we calculated the bottom 1st percentile of gene expression per gene within the TCGA cohort entities.

