## Supplementary figures and tables for "Integrative multi-omics combined with functional pharmacological profiling in patient-derived organoids identifies personalized therapeutic vulnerabilities of adult high-grade gliomas"

bioRχiv

[1]NORLUX Neuro-Oncology Laboratory, Department of Cancer Research, Luxembourg Institute of Health, L-1210 Luxembourg, Luxembourg; [2]School of Biosciences and Veterinary Medicine, University of Camerino, 62032 Camerino, Italy; [3]Goethe University Frankfurt, University Hospital, Dr. Senckenberg Institute of Neuroncology, Edinger Institute, Institute of Neurology, Frankfurt Cancer Institute, University Cancer Center Frankfurt, 60528 Frankfurt am Main, Germany; [4]Multiomics Data Science, Department of Cancer Research, Luxembourg Institute of Health, L-1445 Strassen, Luxembourg; [5]Bioinformatics and AI Unit, Department of Medical Informatics, Luxembourg Institute of Health, L-1445 Strassen, Luxembourg; [6]Animal Facility, Department of Cancer Research, Luxembourg Institute of Health, 29 Rue Henri Koch, L-4354 Esch-Sur-Alzette, Luxembourg; [7]Cancer RNAs and Epigenetic group, Department of Cancer Research, Luxembourg Institute of Health, L-1210 Luxembourg, Luxembourg; [8]Division of Experimental Neurosurgery, Department of Neurosurgery, University of Heidelberg, 69120 Heidelberg, Germany; [9]Luxembourg Center of Neuropathology (LCNP) and National Center of Pathology (NCP), Laboratoire National de Santé, Luxembourg Institute of Health, L-3555 Dudelange, Luxembourg; [10]Institute for Neuropathology, University Medical Center of the Johannes Gutenberg University, Mainz, 55131, Germany; [11]Department of Life Science and Medicine (DLSM), Faculty of Science, Technology and Medicine (FSTM), University of Luxembourg, L-4365 Esch-sur-Alzette, Luxembourg; [12]Centre Hospitalier de Luxembourg, Luxembourg L-1526, Luxembourg; [13]Luxembourg Institute of Health, L-1210 Luxembourg, Luxembourg; [14]School of Pharmaceutical Sciences of Ribeirão Preto, University of São Paulo, Ribeirão Preto, São Paulo, Brazil

[#]Equal contribution

### SUPPLEMENTARY FIGURES

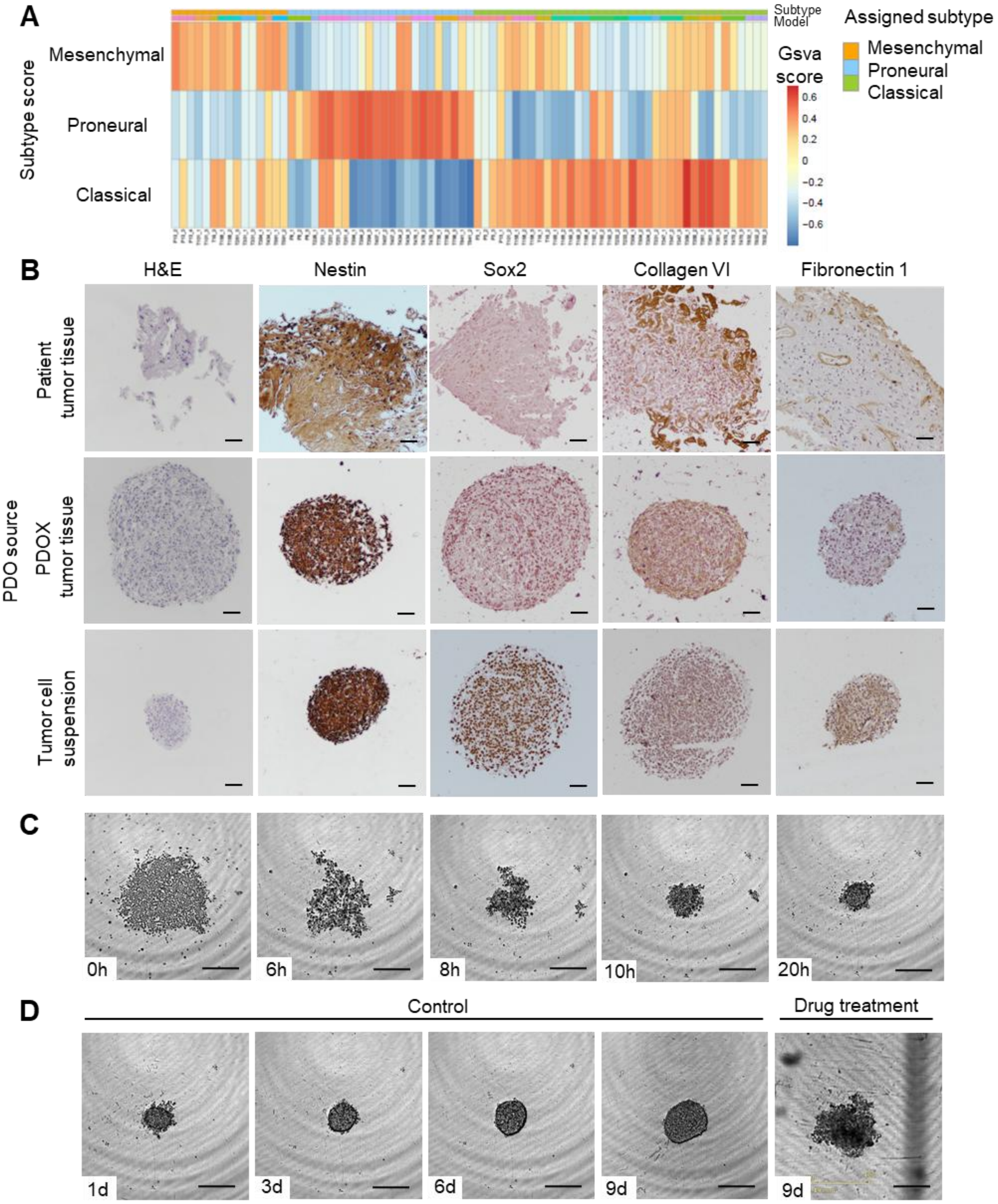

**Fig. S1. Characterisation of molecular and histopathological features of high-grade glioma models. A.** The heatmap showing Wang et al.[1] glioma tumor-intrinsic transcriptional profiling of the preclinical models, gsva score is shown for the three transcriptional subtypes. **B**. Representative IHC of PDOs (model T188) derived from patient tumor tissue (150-750µm), PDOX tumor tissue (200-350µm) and self-assembled PDOs from tumor cell suspension (130-150µm) showing recapitulation of glioma-specific tumor markers (Nestin, Sox2) and extracellular matrix (Collagen IV, Fibronectin 1); magnification: 10x; scale bar: 50µm. **C**. Representative images of self-assembling PDOs (model P3) from MACS-purified tumor cells, scale bar 50µm. **D.** IncuCyte-based monitoring of PDO size and structure upon *ex vivo* culture (day 1-9) and PDO death after drug treatment (representative example at 144h of treatment for 20µM RGFP966). Examples are shown for PDO T407, scale bar: 50µm.

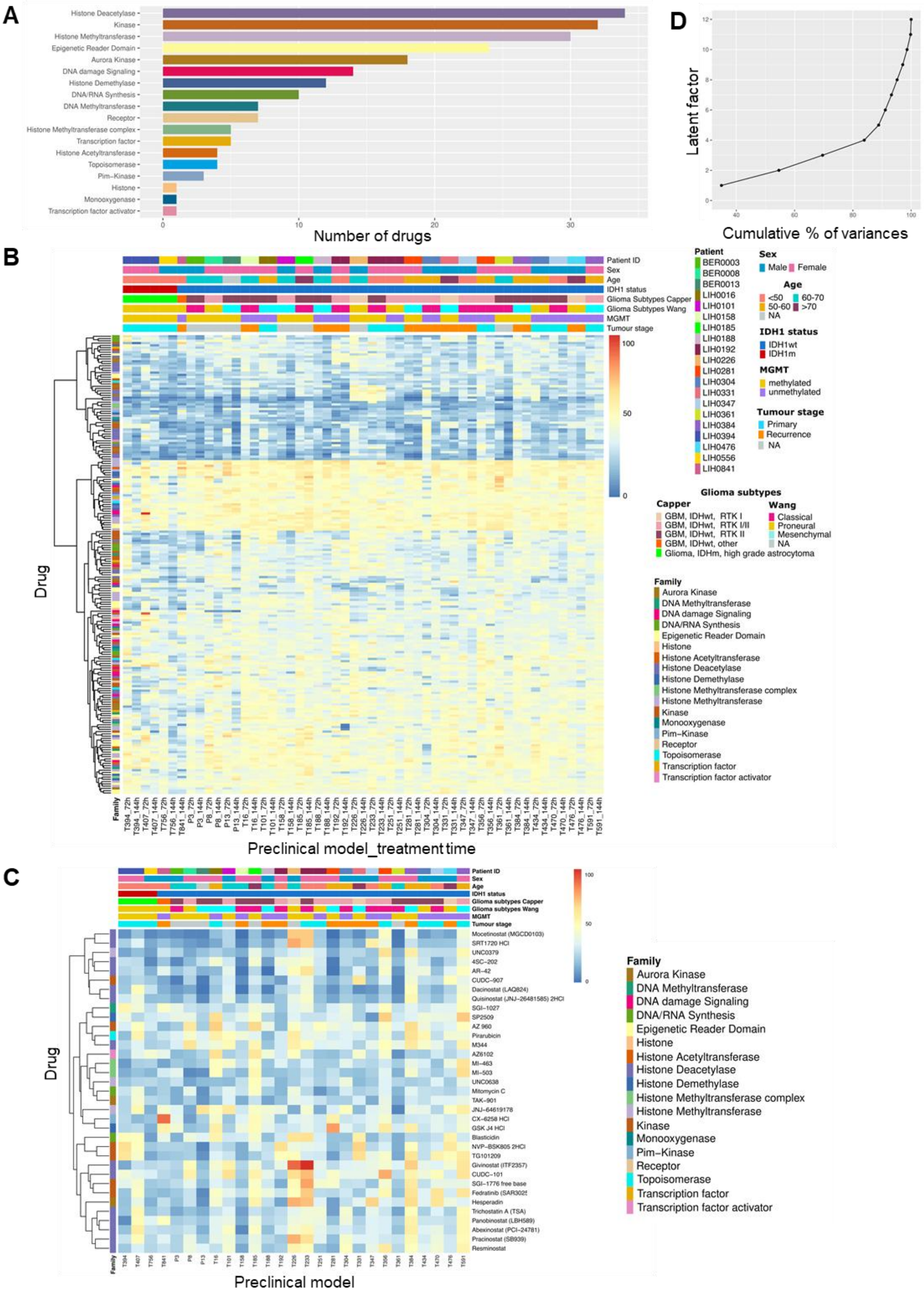

**Fig. S2. *Ex vivo* drug screening in PDOs**. **A**. Main drug target families of compounds applied in the screen. Inhibitors were grouped into 16 drug families, according to their targets**,** indicated by color code**.** The majority of drugs target epigenetic regulators. **B.** Heatmap displaying drug responses in 27 models *ex vivo* at 72h and 144h. Values represent % of viability normalized to DMSO-treated control. Mean values of replicates (n=3) per model per time are shown. Mean of technical controls per plate was used for normalization (='100%'). Glioma subtypes ((Capper et al.[2] Wang et al.[1]), tumor stage (primary vs. recurrent tumors), *MGMT* promoter methylation and *IDH1* mutation status are indicated per preclinical model. Hierarchical clustering was performed for drug responses. **C.** Heatmap of % of viability at 144h clustered by response displayed for the 35 most efficient inhibitors that reduced cell viability to <50% in more than 50% of the tested 27 models. **D.** The cumulative proportion of variance explained by MOFA fittings indicates that first six latent factors explain 90% of the variance.

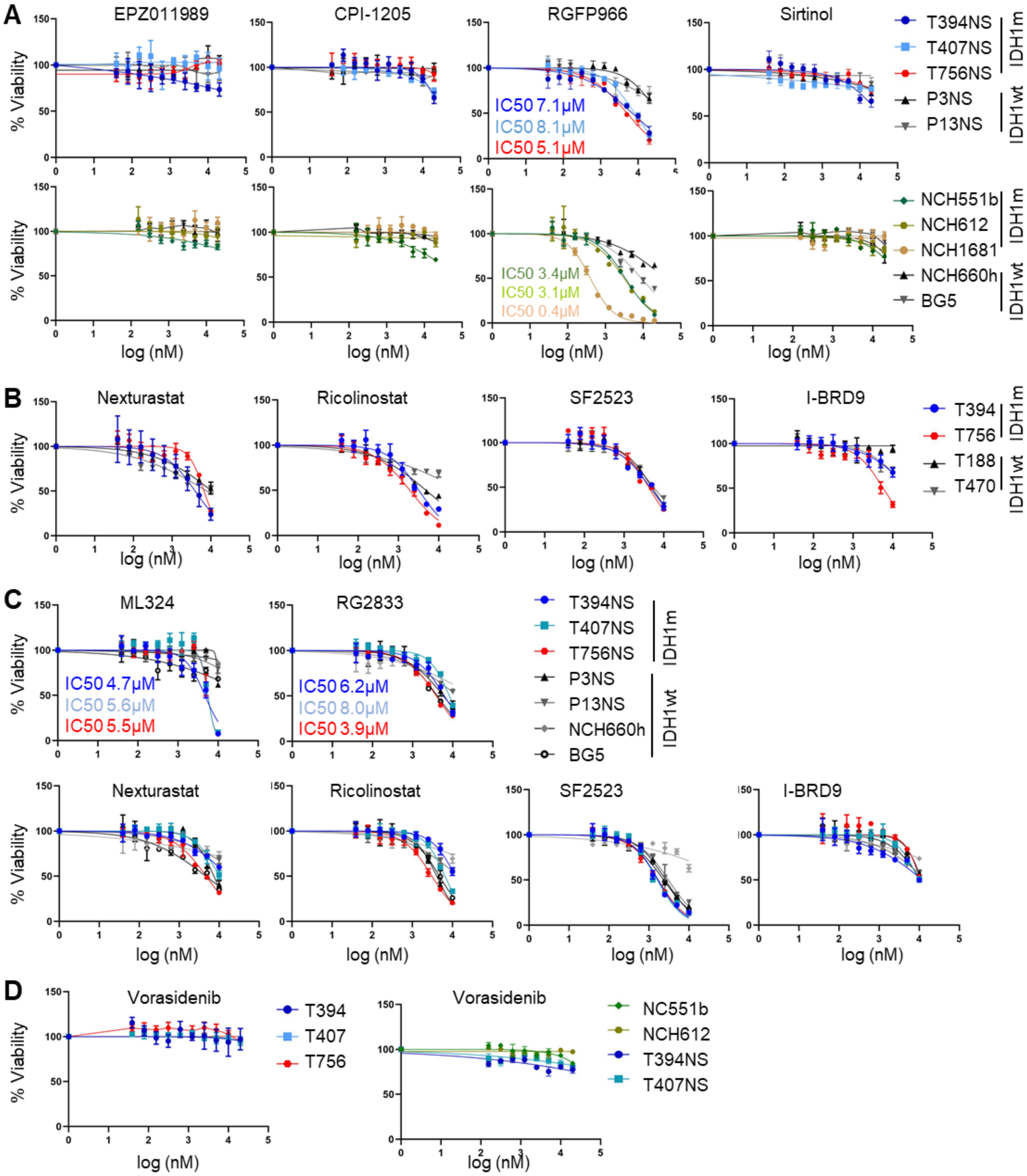

**Figure S3. LF1 and LF4-associated drug sensitivities discriminating IDH1m high-grade astrocytomas.** **A.** Dose-response validation of LF1-identified drugs with mean % viability <50% in IDH1m glioma GSCs compared to IDH1wt GBM counterparts. The top panel shows GSCs derived from the in-house preclinical models, lower panel shows additional GSCs derived from an independent patient cohort. **B.** Dose-response validation of LF4-identified drugs in IDH1m high-grade astrocytoma PDOs (T756 and T394) and IDH1wt GBM PDOs (T188 and T470) with broader toxicity. **C.** Dose-response validation of LF4-identified drugs in IDH1m glioma GSCs compared to GBM counterparts. **D.** Dose-response validation of Vorasedinib (FDA-approved drug for low-grade IDH1/2m gliomas) across IDH1m PDOs (left) and GSCs (right).

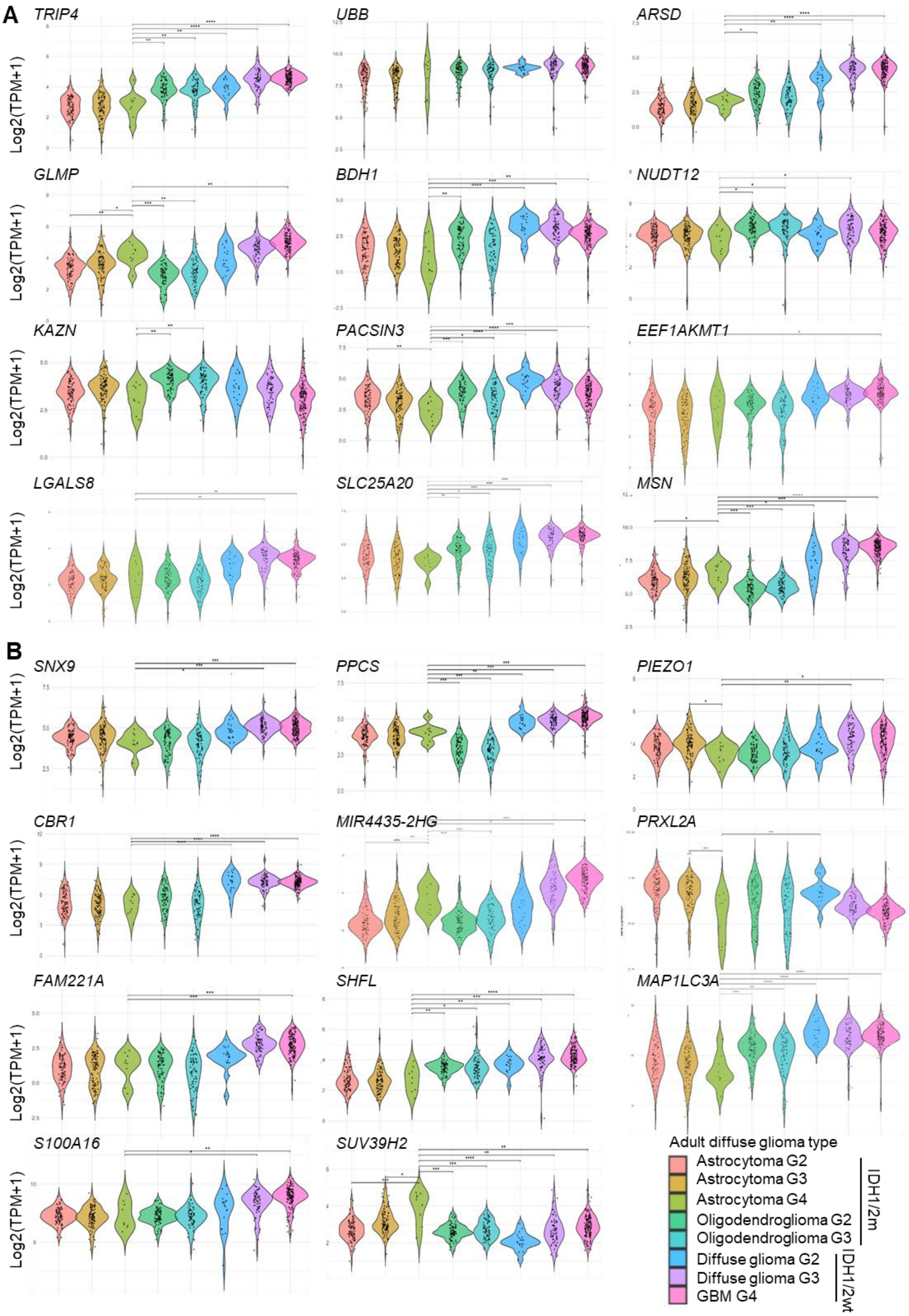

**Figure S4. LF1/LF4-associated gene expression across adult diffuse gliomas.** Violin plots showing the gene expression distribution across a curated LGG-GBM TCGA dataset for LF1 (**A**) and LF4 (**B**) associated gene transcripts. Diffuse glioma tumor entities were curated based on WHO 2021 categories and grades (Gs). Expression profiles are shown for: IDH1/2m 1p/19q intact astrocytomas grade 2 (n=85), grade 3 (n=96) and grade 4 (n= 9); IDH1/2m 1p/19q co-deleted oligodendrogliomas grade 2 (n=79) and grade 3 (n=68); IDH1/2wt gliomas grade 2 (n=19), grade 3 (n=67), and grade 4 GBMs (n=141). The majority of genes show lower expression levels in IDH1/2m grade 4 astrocytomas and/or IDH1/2m gliomas in general, compared to IDH1/2wt GBMs. SUV39H2 shows lower levels in GBMs. Statistical analysis was performed using a fixed pairwise comparison against astrocytoma grade 4. The Wilcoxon test was used, with FDR correction applied for multiple comparisons. Significant p-values are denoted as: *p<0.05, **p<0.01, ***p<0.001, ****p<0.0001.

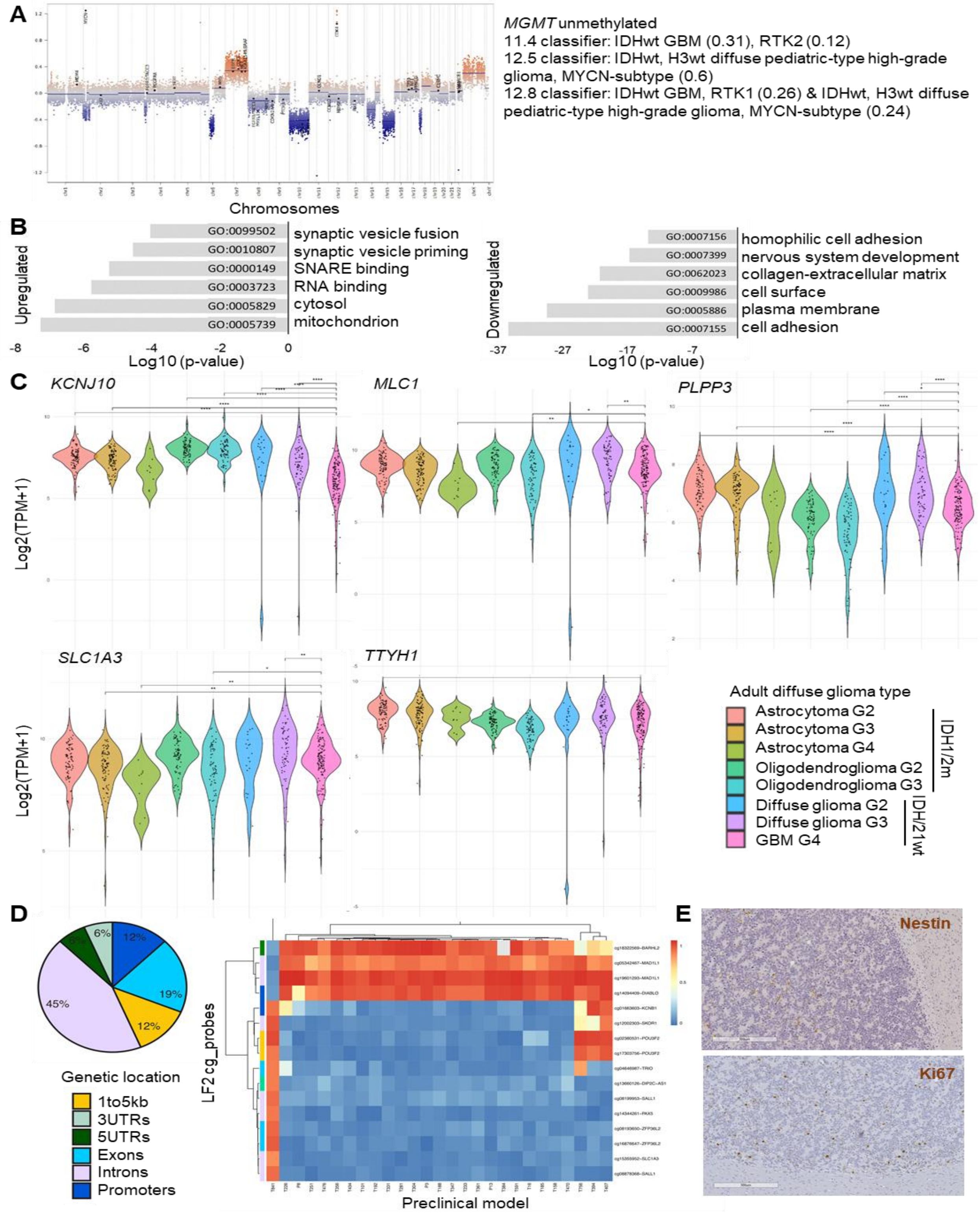

**Figure S6. Assessment of rare *MYCN*-amplified GBM tumor features.** **A.** DNA methylation-based CNV profile of T841 patient tumor showing *MYCN* and *CDK4* amplification. GBM-characteristic +7/-10 Chr feature is present. 11.4, 12.5 and 12.8 Heidelberg classifier scores show no clear match to molecular glioma subtype. **B.** Differential gene expression analysis between T841 vs. other models (DEGs: FDR ≤0.01 and |log2FC|≥1). Top six GO terms are displayed characterising up- and downregulated genes. **C.** Violin plots showing the LF2-associated gene expression distribution across a curated LGG-GBM TCGA dataset. Glioma subgroups were curated based on WHO 2021 categories and grades. Expression profiles are shown for: IDH1/2m 1p/19q intact astrocytomas grade 2 (n=85), grade 3 (n=96) and grade 4 (n= 9); IDH1/2m 1p/19q co-deleted oligodendrogliomas grade 2 (n=79) and grade 3 (n=68); IDH1/2wt gliomas grade 2 (n=19), grade 3 (n=67), and grade 4 GBMs (n=141). A subgroup of patients presents lower gene expression levels compared to the average distribution within molecularly-defined glioma entities. Statistical analysis was performed using a fixed pairwise comparison against GBM. The Wilcoxon test was used, with FDR correction applied for multiple comparisons. Significant p-values are denoted as: *p<0.05, **p<0.01, ***p<0.001, ****p<0.0001. **D.** Epigenetic features discriminating T841 from other models. Pie chart shows distribution of LF2-associated DNA methylation cp probes across genetic locations. Heatmap presenting β-values per each DNA methylation probe identified in LF4 as unique to T841 model. **E.** Representative Nestin (human-specific) and Ki67 staining in PDOX tumors *in vivo*; Scale bar = 300µm for magnified areas. T841 does not show Nestin positivity, common for IDH1wt GBMs. Ki67 depicts proliferative cells.

### SUPPLEMENTARY TABLES

**Table S1. Characteristic of preclinical models applied in the study.** The table depicts key patient characteristics (age, sex, tumor stage and diagnosis). Molecular profiles correspond to features detected in the preclinical models.

**Table S2. List of inhibitors applied in the study.** 202 drugs are numbered in the first column. Each line represents a specific target and associated gene family per each drug.

**Table S3. Drug efficacy across the PDO cohort.** The table depicts median drug viability across the preclinical cohort at 72h and 144h per each individual compound (columns 3-4 respectively) as well as the number of models achieving ≤ 25% and ≤ 50% viability at 144h (columns 5-6 respectively).

**Table S4. Loading scores of the variables identified in the genomic, epigenomic, transcriptomic, and drug layers in the LF1, LF2, and LF4.** The features with loading scores > 0.75 are displayed, except for the epigenomics layer, which shows markers with the highest loading scores >0.9.

**Table S5. Gene ontology functional annotation of differentially expressed genes.** Differential gene expression analysis was performed between (a) IDH1m astrocytoma (T394, T407, T756) versus GBM models; (b) within IDH1m models (T756 versus T394/T407) and (c) T841 *MYCN* amplified GBM model versus other models in the cohort. Differentially expressed genes were defined at threshold: FDR ≤ 0.01 and |log2FC| ≥ 1. Top 30 Gene ontology (GO) terms are shown separately for up and down-regulated genes if p-value ≤ 0.01.

**Table S6. Primary antibodies for Immunohistochemistry (IHC).**

**Table S1. Characteristic of preclinical models applied in the study.**

The table depicts key patient characteristics (age, sex, tumor stage and diagnosis). Molecular profiles correspond to features detected in the preclinical models.

| Model | Patient information | | | | | | | | Molecular features in preclinical models | | | | | |
|---|---|---|---|---|---|---|---|---|---|---|---|---|---|---|
| | Patient code | Age at collection | Sex | Tumor localization | Surgery | Treatment rece4ed prior model der4ation | Histological diagnosis | Molecular diagnosis (Capper et al.) | Copy number variations (CNV) | Gene mutations | MGMT promoter methylation | Molecular diagnosis (Capper et al.) | DNA methylation subtypes (Malta et al.,) | Transcriptional subtype (Wang et al.) |
| P3 | BER0003 | 64 | Male | NA | NA | NA | GBM Grade 4 | NA | +[Chr7, 19p, 20q], -[1p36-1p34.1, 1q21.1-q44, -5p15.33-31, Chr9, Chr10, 11p15-14, 20p] --CDKN2A/B | ALK (chr2:g.29320870A>G, ENSP00000373700.3:p.Val476Ala)<br>BRCA2 (chr13:g.32398489A>T, ENSP00000369497.3:p.Lys3326Ter)<br>COL6A3 (chr2:g.237340908C>T, ENSP00000295550.4:p.Ala2670Thr)<br>GABRA6 (chr5:g.161687516T>C, ENSP00000429943.1:p.Leu13Pro)<br>GABRA6 (chr5:g.161687516T>C, ENSP00000429943.1:p.Leu13Pro)<br>GNAS (chr20:g.58855063C>G, ENSP00000265621.6:p.Arg600Gly)<br>IGF1R (chr15:g.98922429C>T, ENSP00000496919.1:p.Ala828Val)<br>KMT2A (chr11:g.118472663G>A, ENSP00000374157.5:p.Glu502Lys)<br>NUP210L (chr1:g.154100125G>A, ENSP00000271854.3:p.Pro613Leu)<br>PRKAR1A (chr17:g.68530055G>A, ENSP00000466314.2:p.Val343Met)<br>PTCH2 (chr1:g.44829624C>T, ENSP00000361266.3:p.Arg358His)<br>PTEN (chr10:g.87957923del, ENSP00000361021.3:p.Asp236ThrfsTer20)<br>TP53 (chr17:g.7675085C>T, ENSP00000269305.4:p.Cys176Tyr)<br>TSC2 (chr16:g.2087968_2088001del:-:) | methylated | GBM, IDHwt, RTK II | Classic-like | Classical |
| P8 | BER0008 | 64 | Female | NA | NA | NA | GBM Grade 4 | GBM, IDHwt, RTK I | ++EGFR, +[5p15.3-p12, 5q31.1-q35.3, Chr7, 8q24], -[ 6q21-q27, Chr10, 13q13.3-q34, 15q21.2-q23, 18q21.2-q22.4], --CDKN2A/B | ALK (chr2:g.29320870A>G, ENSP00000373700.3:p.Val476Ala)<br>ATR (chr3:g.142555928T>C, ENSP00000343741.4:p.Lys764Glu)<br>ATRX (chrX:g.77663459C>G, ENSP00000362441.4:p.Ser1348Thr)<br>ATRX (chrX:g.77683638T>A, ENSP00000362441.4:p.Ser540Cys)<br>EGFR (chr7:g.55154128G>A, ENSP00000275493.2:p.Ala289Thr)<br>ETS1 (chr11:g.128489374C>T, ENSP00000324578.5:p.Ala107Thr)<br>FLT3 (chr13:g.28100491T>C, ENSP00000241453.7:p.Asp7Gly)<br>MET (chr7:g.116771573G>T, ENSP00000317272.6:p.Ala954Ser)<br>MUC17 (chr7:g.101035233A>G, ENSP00000302716.4:p.Thr1273Ala)<br>NCOR1 (chr17:g.16047086C>T, ENSP00000268712.2:p.Ala2182Thr)<br>PRKAR1A (chr17:g.68551108G>A, ENSP00000467311.2:p.Ser333Asn)<br>PTEN (chr10:g.87961047_87961050del, ENSP00000361021.3:p.Thr319Ter)<br>ZNF292 (chr6:g.87245506A>T, ENSP00000342847.4:p.Glu289Asp) | methylated | GBM, IDHwt, RTK I/II | Classic-like | Proneural |
| P13 | BER0013 | NA | Female | NA | NA | NA | GBM Grade 4 | NA | +(Chr7, Chr19, Chr20), -(1p21.1-p31.2, 6q16.3-q21, Chr10, Chr13, 17q11-12), --CDKN2A/B | ARID2 (chr12:g.45776079A>G:-:)<br>ATR (chr3:g.142562511C>G, ENSP00000343741.4:p.Lys297Asn)<br>CSF1R (chr5:g.150070264C>T, ENSP00000286301.3:p.Gly413Ser)<br>FLT3 (chr13:g.28100491T>C, ENSP00000241453.7:p.Asp7Gly)<br>HMCN1 (chr1:g.186152863_186152864delinsTT, ENSP0000271588.4:p.Thr5004Phe)<br>IDH2 (chr15:g.90087524A>G, ENSP00000331897.4:p.Trp244Arg)<br>KDM1B (chr6:g.18171430A>C, ENSP00000297792.5:p.Gln162Pro)<br>KDM5A (chr12:g.387273A>T, ENSP00000444251.1:p.Phe56Tyr)<br>KMT2C (chr7:g.152252650T>C, ENSP00000262189.6:p.Ile455Met)<br>LRP1B (chr2:g.140274452T>A, ENSP00000374135.3:p.Asn4372Tyr)<br>LRP1B (chr2:g.140358851C>T, ENSP00000374135.3:p.Gly3743Ser)<br>LTBP4 (chr19:g.40610549C>G, ENSP00000204005.10:p.Arg598Gly)<br>MIIP (chr1:g.12031353G>A, ENSP00000235332.4:p.Ala344Thr)<br>MPL (chr1:g.43338669G>A, ENSP00000361548.3:p.Val114Met)<br>NF1 (chr17:g.31221849G>A:-:)- not pathogenic<br>PRKAR1A (chr17:g.68551108G>A, ENSP00000467311.2:p.Ser333Asn)<br>PTPN11 (chr12:g.112489068C>T, ENSP00000340944.3:p.Arg498Trp)<br>SETD7 (chr4:g.139496517G>C, ENSP00000427300.1:p.Gln309Glu)<br>SETD7 (chr4:g.139602701G>A, ENSP00000476302.1:p.His25Tyr)<br>SMARCA4 (chr19:g.10987920T>C, ENSP00000343896.4:p.Tyr372His)<br>SMO (chr7:g.129189225A>G, ENSP00000249373.3:p.Asp25Gly) | methylated | GBM, IDHwt, RTK II | Classical | Mesenchymal |
| T16 | LIH0016 | 52 | Female | Right fronto-parietal | recurrence | TMZ | GBM Grade 4 | GBM, IDHwt, mesenchymal | ++[EGFR, MDM4], +7q, - [6q, Chr10, 13q11-q31.1], --CDKN2A/B | ABL1 (chr9:g.130885205C>T, ENSP00000323315.5:p.Ser972Leu)<br>ABL1 (chr9:g.130885403C>T, ENSP00000323315.5:p.Ala1038Val)<br>AURKB (chr17:g.8206543C>G, ENSP00000313950.6:p.Gly213Arg)<br>BCOR (chrX:g.40052111A>T, ENSP00000345923.4:p.Ter1722ArgextTer34)<br>COL6A3 (chr2:g.237340715C>T, ENSP00000295550.4:p.Arg2734Gln)<br>COL6A3 (chr2:g.237366926T>C, ENSP00000295550.4:p.Lys1754Arg)<br>FGFR4 (chr5:g.177096332C>T, ENSP00000292408.4:p.Arg664Trp)<br>GABRA6 (chr5:g.161687516T>C, ENSP00000429943.1:p.Leu13Pro)<br>GABRA6 (chr5:g.161687516T>C, ENSP00000429943.1:p.Leu13Pro)<br>GSE1 (chr16:g.85657562C>T, ENSP00000253458.6:p.Pro533Leu)<br>KMT2D (chr12:g.49041686C>T, ENSP00000301067.7:p.Arg2068Gln)<br>LZTR1 (chr22:g.20988119G>A:-:)<br>MUC17 (chr7:g.101036559G>A, ENSP00000302716.4:p.Val1715Ile)<br>TSC2 (chr16:g.2053484C>T, ENSP00000387937.3:p.Thr18Met) | unmethylated | GBM, IDHwt, RTK II | Classical | Mesenchymal |
| T101 | LIH0101 | 60 | Male | NA | NA | NA | GBM Grade 4 | NA | ++[EGFR, MDM2] +[Chr1, Chr7, Chr9, Chr13, Chr16, Chr17, Chr19] -[3q, Chr4, Chr6, Chr10, Chr11, 14q11, Chr15], --CDKN2A/B, | AKAP9 (chr7:g.92001968A>G, ENSP00000348573.3:p.Glu684Gly)<br>DICER1 (chr14:g.95124241C>A, ENSP00000343745.3:p.Gly444Val)<br>GABRA6 (chr5:g.161687516T>C, ENSP00000429943.1:p.Leu13Pro)<br>GABRA6 (chr5:g.161687516T>C, ENSP00000429943.1:p.Leu13Pro)<br>KMT2A (chr11:g.118506285A>G, ENSP00000374157.5:p.Lys3462Glu)<br>MUC17 (chr7:g.101031186G>A, ENSP00000302716.4:p.Arg50His)<br>MYB (chr6:g.135192509C>A, ENSP00000326328.8:p.Pro238His)<br>PIK3C2G (chr12:g.18290875C>A, ENSP00000404845.1:p.Ala261Glu)<br>PRKAR1A (chr17:g.68551108G>A, ENSP00000467311.2:p.Ser333Asn)<br>TET2 (chr4:g.105235030C>T, ENSP00000265149.5:p.Pro363Leu) | methylated | GBM, IDHwt, RTK I/II | Classical | Mesenchymal |
| T158 | LIH0158 | 70 | Female | Right frontal | primary | no prior treatment | GBM Grade 4 | GBM, IDHwt, mesenchymal | [++MDM2], +[Chr 7, Chr19, Chr20], -[Chr10, Chr15q], --CDKN2A/B | AKAP9 (chr7:g.92000991A>T, ENSP00000348573.3:p.Leu358Phe)<br>APC (chr5:g.112843416G>A, ENSP00000257430.4:p.Ala2608Thr)<br>ARID1B (chr6:g.156829362A>G, ENSP00000344546.5:p.Ile643Val)<br>ATM (chr11:g.108310190T>C, ENSP00000435524.1:p.Leu73Pro)<br>AURKA (chr20:g.56388155_56388158del:-:)<br>CDK6 (chr7:g.92615172T>A, ENSP00000265734.4:p.Ser317Cys)<br>COL6A3 (chr2:g.237387842A>G, ENSP00000295550.4:p.Val351Ala)<br>EPHA3 (chr3:g.89450214C>G, ENSP00000337451.2:p.Pro845Arg)<br>FGFR3 (chr4:g.1805767G>T, ENSP00000339824.4:p.Val557Leu)<br>GABRA6 (chr5:g.161687516T>C, ENSP00000429943.1:p.Leu13Pro)<br>H2BC9 (chr6:g.26251760_26251761del, ENSP00000479169.1:p.Ser37IlefsTer80)<br>IDO2 (chr8:g.40013579C>T, ENSP00000443432.2:p.Pro245Leu)<br>KDM1B (chr6:g.18201603G>A, ENSP00000405669.2:p.Val493Ile)<br>KDM4B (chr19:g.5151426C>T, ENSP00000159111.3:p.Thr1069Met)<br>KDM5A (chr12:g.387273A>T, ENSP00000444251.1:p.Phe56Tyr)<br>LTBP4 (chr19:g.40610549C>G, ENSP00000204005.10:p.Arg598Gly)<br>MSH4 (chr1:g.75880138A>G, ENSP00000263187.3:p.Tyr589Cys)<br>MYB (chr6:g.135202445T>C, ENSP00000326328.8:p.Leu577Pro)<br>MYB (chr6:g.135202445T>C, ENSP00000326328.8:p.Leu577Pro)<br>NF1 (chr17:g.31163302G>A, ENSP00000491431.1:p.Gly31Glu)- not pathogenic<br>NOTCH1 (chr9:g.136505644C>T, ENSP00000498587.1:p.Ala1418Thr)<br>PLCG1 (chr20:g.41166762_41166764del, ENSP00000244007.3:p.Val735del)<br>SETD7 (chr4:g.139496517G>C, ENSP00000427300.1:p.Gln309Glu)<br>ZNF521 (chr18:g.25224410T>C, ENSP00000354794.3:p.Ser1170Gly) | unmethylated | GBM, IDHwt, RTK II | Classic-like | Classical |
| T185 | LIH0185 | 76 | Female | Right occipital | NA | no prior treatment | GBM Grade 4 | GBM, IDHwt, RTK II | ++EGFR, +[Chr7, Chr20], -[Chr 10q, Chr22q] --CDKN2A/B | ALK (chr2:g.29320870A>G, ENSP00000373700.3:p.Val476Ala)<br>AURKB (chr17:g.8204933G>A, ENSP00000313950.6:p.Pro326Ser)<br>COL6A3 (chr2:g.237344688G>A, ENSP00000295550.4:p.Arg2444Trp)<br>EGFR (chr7:g.55165350G>T, ENSP00000275493.2:p.Gly598Val)<br>GABRA6 (chr5:g.161687516T>C, ENSP00000429943.1:p.Leu13Pro)<br>GNAS (chr20:g.58854392C>T, ENSP00000302237.4:p.Arg378Trp)<br>GSE1 (chr16:g.85648614G>A, ENSP00000253458.6:p.Ala97Thr)<br>MET (chr7:g.116695757A>T, ENSP00000413857.1:p.Met1?)<br>MUC17 (chr7:g.101039562C>G, ENSP00000302716.4:p.Pro2716Ala)<br>NF1 (chr17:g.31350201A>G, ENSP00000348498.3:p.Glu2426Gly)<br>PLCG1 (chr20:g.41172488C>T, ENSP00000468958.1:p.Thr50Met)<br>PRKAR1A (chr17:g.68551108G>A, ENSP00000467311.2:p.Ser333Asn)<br>STAG2 (chrX:g.124090748A>G, ENSP00000218089.9:p.Thr1151Ala)<br>TRAF7 (chr16:g.2173517A>C, ENSP00000318944.6:p.Asp350Ala)<br>ZNF292 (chr6:g.87249373G>T, ENSP00000428073.2:p.Arg344Leu) | methylated | GBM, IDHwt, RTK II | Classic-like | Classical |
| T186 | LIH0186 | 34 | Male | Supratentorial | recurrence | no prior treatment | Anaplastic Oligodendrogliom a Grade 3 | Glioma, IDHm, high-grade astrocytoma | complex genome ++[MYCN, CDK4] | ALK (chr2:g.29320870A>G, ENSP00000373700.3:p.Val476Ala)<br>APC (chr5:g.112707836G>C, ENSP00000481752.1:p.Ser40Thr)<br>APC (chr5:g.112839543G>C, ENSP00000257430.4:p.Glu1317Gln)<br>ARID2 (chr12:g.45891802G>C, ENSP00000335044.6:p.Val1649Leu)<br>ATRX (chrX:g.77652140C>A, ENSP00000362441.4:p.Glu1511Ter)<br>BRPF3 (chr6:g.36225308G>A, ENSP00000345419.7:p.Glu805Lys)<br>DNMT3A (chr2:g.25300211C>G, ENSP00000264709.3:p.Lys35Asn)<br>GABRA6 (chr5:g.161687516T>C, ENSP00000429943.1:p.Leu13Pro)<br>GSE1 (chr16:g.85666291A>G, ENSP00000253458.6:p.His1025Arg)<br>IDH1 (chr2:g.208248388C>T, ENSP00000260985.2:p.Arg132His)<br>KMT2C (chr7:g.152252650T>C, ENSP00000262189.6:p.Ile455Met)<br>MET (chr7:g.116771936C>T, ENSP00000317272.6:p.Thr1010Ile)<br>MPL (chr1:g.43338669G>A, ENSP00000361548.3:p.Val114Met)<br>NOTCH2 (chr1:g.119915647G>C, ENSP00000256646.2:p.Pro2359Ala)<br>PDGFRA (chr4:g.54285926A>T, ENSP00000257290.5:p.Asp842Val)<br>PHC2 (chr1:g.33370563G>A, ENSP00000257118.5:p.Ala145Val)<br>PIK3C2G (chr12:g.18290875C>A, ENSP00000404845.1:p.Ala261Glu)<br>PRKAR1A (chr17:g.68551108G>A, ENSP00000467311.2:p.Ser333Asn)<br>RET (chr10:g.43100551C>A, ENSP00000344798.4:p.Leu56Met)<br>TP53 (chr17:g.7673803G>A, ENSP00000269305.4:p.Arg273Cys) | methylated | Glioma, IDHm, high-grade astrocytoma | K1: G-CIMP-low | NA |
| T188 | LIH0188 | 68 | Male | Right temporal | primary | no prior treatment | GBM Grade 4 | GBM, IDHwt, RTK II | [++EGFR], +[Chr7, Chr 19, Chr 20], -[1p36.23, 6p21.32, 6q27, Chr 10, 11q24.2], --CDKN2A/B | AKT1 (chr14:g.104773520C>T, ENSP00000270202.4:p.Ala255Thr)<br>FLT3 (chr13:g.28100491T>C, ENSP00000241453.7:p.Asp7Gly)<br>GABRA6 (chr5:g.161687516T>C, ENSP00000429943.1:p.Leu13Pro)<br>GABRA6 (chr5:g.161687516T>C, ENSP00000429943.1:p.Leu13Pro)<br>GNAS (chr20:g.58854564A>G, ENSP00000302237.4:p.Gln435Arg)<br>GSE1 (chr16:g.85672270C>T, ENSP00000410812.1:p.Thr405Ile)<br>H2BC12 (chr6:g.27146670G>C, ENSP00000349430.2:p.Tyr43Ter)<br>HMCN1 (chr1:g.186144548C>T, ENSP00000271588.4:p.Ala4704Val)<br>KIT (chr4:g.54707250G>C, ENSP00000288135.6:p.Glu360Gln)<br>LRP1B (chr2:g.140274452T>A, ENSP00000374135.3:p.Asn4372Tyr)<br>LRP1B (chr2:g.140358851C>T, ENSP00000374135.3:p.Gly3743Ser)<br>MDM4 (chr1:g.204549329A>C, ENSP00000356150.3:p.Lys374Gln)<br>MLH1 (chr3:g.37047639_37047640delinsGC, ENSP00000231790.3:p.Lys618Ala)<br>NOTCH1 (chr9:g.136506781C>T, ENSP00000498587.1:p.Arg1279His)<br>NOTCH1 (chr9:g.136511197C>T, ENSP00000498587.1:p.Glu848Lys)<br>NOTCH2 (chr1:g.119915499A>T, ENSP00000256646.2:p.Leu2408His)<br>PTCH1 (chr9:g.95459693C>T, ENSP00000332353.6:p.Val932Ile)<br>SETD7 (chr4:g.139496517G>C, ENSP00000427300.1:p.Gln309Glu)<br>TERT (chr5:g.1279412G>A, ENSP00000309572.5:p.Ala670Val)<br>TET2 (chr4:g.105236541T>C, ENSP00000265149.5:p.Tyr867His)<br>TET2 (chr4:g.105275677C>T, ENSP00000369351.4:p.Pro1723Ser)<br>TSC2 (chr16:g.2083700G>A, ENSP00000219476.3:p.Ala1297Thr) | unmethylated | GBM, IDHwt, RTK II | Classic-like | Mesenchymal |
| T192 | LIH0192 | 42 | Female | Left frontal | primary | no prior treatment | GBM Grade 4 | GBM, IDHwt, mesenchymal | ++[EGFR, 2q34], +[1q21.2-24.2, Chr7, Chr19, Chr20], - [9p24.3 -13.3, Chr10], | CREB1 (chr2:g.207560248C>T, ENSP00000236995.3:p.Ala46Val)<br>EGFR (chr7:g.55154017C>T, ENSP00000275493.2:p.Arg252Cys)<br>GABRA6 (chr5:g.161687516T>C, ENSP00000429943.1:p.Leu13Pro)<br>H2AC17 (chr6:g.27893145G>A, ENSP00000352627.3:p.Ser2Phe)<br>HMCN1 (chr1:g.186093576G>C, ENSP00000271588.4:p.Gly3368Ala)<br>JMJD1C (chr10:g.63193075G>A, ENSP00000335929.5:p.Pro527Leu)<br>KDM5A (chr12:g.387273A>T, ENSP00000444251.1:p.Phe56Tyr)<br>MSH2 (chr2:g.47416318G>T, ENSP00000233146.2:p.Gly322Val)<br>SETD1B (chr12:g.121823309C>T, ENSP00000442924.1:p.Ala1534Val)<br>SETD7 (chr4:g.139496517G>C, ENSP00000427300.1:p.Gln309Glu)<br>TERT (chr5:g.1254479C>T, ENSP00000309572.5:p.Ala1062Thr)<br>TET1 (chr10:g.68574194G>C, ENSP00000362748.4:p.Cys619Ser) | unmethylated | GBM, IDHwt, RTK I/II | Mesenchymal-like | Classical |
| T226 | LIH0226 | 61 | Male | Supratentorial | NA | NA | GBM Grade 4 | NA, low tumor content | ++[EGFR], +[Chr 5p, Chr7, Chr 20p], -[1p21.1-p31.2, 6p22.3, 6q16.3-q21, Chr 10, 14q, 16p12.2, 19q, 22q], , --CDKN2A/B | ARID1B (chr6:g.157186370C>T, ENSP00000313006.7:p.Pro650Leu)<br>ARID2 (chr12:g.45837568G>C, ENSP00000335044.6:p.Gln397His)<br>CREBBP (chr16:g.3731425del, ENSP00000262367.5:p.Ile1649SerfsTer95)<br>EGFR (chr7:g.55173057G>A, ENSP00000275493.2:p.Gly665Asp)<br>GSE1 (chr16:g.85663029A>T, ENSP00000253458.6:p.Glu770Val)<br>KDM1B (chr6:g.18208184G>A, ENSP00000297792.5:p.Gly383Asp)<br>KDM2B (chr12:g.121580856T>C, ENSP00000366271.3:p.His19Arg)<br>KDM5A (chr12:g.387273A>T, ENSP00000444251.1:p.Phe56Tyr)<br>L2HGDH (chr14:g.50237650G>C, ENSP00000261699.4:p.Phe424Leu)<br>MSH2 (chr2:g.47512394G>A, ENSP00000384199.1:p.Arg909Gln)<br>MUC17 (chr7:g.101038011C>A, ENSP00000302716.4:p.Arg2199Ser)<br>PIK3R1 (chr5:g.68296465A>C, ENSP00000513328.1:p.Gln235His)<br>PMS1 (chr2:g.189795810_189795811delinsTA, ENSP00000343888.4:p.Glu59Lys)<br>PRKAR1A (chr17:g.68551108G>A, ENSP00000467311.2:p.Ser333Asn)<br>PTEN (chr10:g.87933154G>A, ENSP00000361021.3:p.Gly132Asp)<br>SETD7 (chr4:g.139496517G>C, ENSP00000427300.1:p.Gln309Glu)<br>SMO (chr7:g.129189225A>G, ENSP00000249373.3:p.Asp25Gly)<br>TSC2 (chr16:g.2070486G>A, ENSP00000219476.3:p.Ala583Thr) | methylated | GBM, IDHwt, RTK I | Classic-like | Proneural |

| | | | | | | | | | | | | |
|---|---|---|---|---|---|---|---|---|---|---|---|---|
| T233 | LIH0192 | 42 | Female | Left frontal | recurrence 1 | radiotherapy + TMZ | GBM Grade 4 | GBM, IDHwt, mesenchymal | ++[EGFR, 2q34], +[1q21.2-24.2, Chr7, Chr19, Chr20], - [9p24.3 -13.3, Chr10, Chr 14q] | AMER1 (chrX:g.64190132A>G, ENSP00000364003.4:p.Leu1052Pro)<br>GABRA6 (chr5:g.161687516T>C, ENSP00000429943.1:p.Leu13Pro)<br>H2AC17 (chr6:g.27893145G>A, ENSP00000352627.3:p.Ser2Phe)<br>HMCN1 (chr1:g.186093576G>C, ENSP00000271588.4:p.Gly3368Ala)<br>JMJD1C (chr10:g.63193075G>A, ENSP00000335929.5:p.Pro527Leu)<br>MSH2 (chr2:g.47416318G>T, ENSP00000233146.2:p.Gly322Val)<br>MTOR (chr1:g.11117022A>G, ENSP00000354558.4:p.Ile2333Thr)<br>MUC17 (chr7:g.101033503A>G, ENSP00000302716.4:p.Asn696Ser)<br>NUP210L (chr1:g.154135949A>G, ENSP00000271854.3:p.Tyr292His)<br>SETD1B (chr12:g.121823309C>T, ENSP00000442924.1:p.Ala1534Val)<br>SETD7 (chr4:g.139496517G>C, ENSP00000427300.1:p.Gln309Glu)<br>TERT (chr5:g.1254479C>T, ENSP00000309572.5:p.Ala1062Thr)<br>TET1 (chr10:g.68574194G>C, ENSP00000362748.4:p.Cys619Ser) | methylated (v. wide range) | GBM, IDHwt, RTK II | Mesenchymal-like | Classical |
| T238 | LIH0238 | 42 | Male | NA | NA | no prior treatment | GBM Grade 4 | NA, inflamatory tissue | complex genome, +[Chr7, Chr19, Chr20] -[4q35.2, 6q, Chr10, 13q, 17q], --CDKN2A/B | BRPF1 (chr3:g.9742228del, ENSP00000507969.1:p.Cys687AlafsTer8)<br>CSF1R (chr5:g.150073459G>C, ENSP00000286301.3:p.Asn308Lys)<br>FLT3 (chr13:g.28100491T>C, ENSP00000241453.7:p.Asp7Gly)<br>GLI1 (chr12:g.57468130G>A, ENSP00000228682.2:p.Arg405Gln)<br>HMCN1 (chr1:g.186019554C>G, ENSP00000271588.4:p.Ile1828Met)<br>HNF1A (chr12:g.120994313_120994314insC, ENSP00000257555.5:p.Pro289AlafsTer28)<br>IDO2 (chr8:g.40005374T>A, ENSP00000443432.2:p.Ser239Thr)<br>KDM5A (chr12:g.387273A>T, ENSP00000444251.1:p.Phe56Tyr)<br>KLF4 (chr9:g.107487535G>A, ENSP00000363804.4:p.His287Tyr)<br>LRP1B (chr2:g.140850171C>G, ENSP00000374135.3:p.Asp1624His)<br>MDM4 (chr1:g.204542937C>G, ENSP00000356147.3:p.Thr107Arg)<br>MPL (chr1:g.43338669G>A, ENSP00000361548.3:p.Val114Met)<br>MUC17 (chr7:g.101036023C>T, ENSP00000302716.4:p.Thr1536Ile)<br>PAX7 (chr1:g.18700783C>T, ENSP00000364524.3:p.Pro306Leu)<br>PTCH1 (chr9:g.95447309T>C, ENSP00000332353.6:p.Tyr1316Cys)<br>PTEN (chr10:g.87961095C>T, ENSP00000361021.3:p.Arg335Ter)<br>SETD7 (chr4:g.139496517G>C, ENSP00000427300.1:p.Gln309Glu)<br>SIRPB1 (chr20:g.1578587C>A, ENSP00000262929.5:p.Val61Leu)<br>SMO (chr7:g.129189225A>G, ENSP00000249373.3:p.Asp25Gly)<br>TP53 (chr17:g.7675157G>A, ENSP00000269305.4:p.Pro152Leu)<br>ZEB1 (chr10:g.31521854A>C, ENSP00000319248.9:p.Gln840Pro) | methylated | GBM, IDHwt, RTK II | Classic-like | NA |
| T239 | LIH0239 | 80 | Male | Parieto-occipital | NA | no prior treatment | GBM Grade 4 | GBM, IDHwt, mesenchymal | +[Chr7, Chr19], -[1p36.32-p34.3, Chr10], --CDKN2A/B | ATM (chr11:g.108227849C>G, ENSP00000278616.4:p.Ser49Cys)<br>AURKB (chr17:g.8206551C>T, ENSP00000313950.6:p.Gly210Glu)<br>DNMT3A (chr2:g.25300227T>G, ENSP00000264709.3:p.Glu30Ala)<br>FOXO3 (chr6:g.108561627C>T, ENSP00000339527.6:p.Ala140Val)<br>HMCN1 (chr1:g.186103595G>A, ENSP00000271588.4:p.Gly3566Asp)<br>KLF7 (chr2:g.207124205C>T, ENSP00000309570.6:p.Arg101Gln)<br>LTBP4 (chr19:g.40629431C>T, ENSP00000204005.10:p.Pro1549Ser)<br>LZTR1 (chr22:g.20982393G>T, ENSP00000397247.1:p.Gly8Trp)<br>MYB (chr6:g.135202445T>C, ENSP00000326328.8:p.Leu577Pro)<br>MYCN (chr2:g.15942237C>T, ENSP00000281043.3:p.Thr58Met)<br>PCDH8 (chr13:g.52846209A>G, ENSP00000341350.4:p.Val743Ala)<br>PCDH8 (chr13:g.52847338T>C, ENSP00000341350.4:p.Thr367Ala)<br>PHF19 (chr9:g.120869321C>T, ENSP00000363003.3:p.Ala159Thr)<br>PIK3C2G (chr12:g.18371180G>T, ENSP00000404845.1:p.Arg542Ser)<br>PIK3R1 (chr5:g.68280986del, ENSP00000513333.1:p.Asn299MetfsTer19)<br>PRKAR1A (chr17:g.68530055G>A, ENSP00000466314.2:p.Val343Met)<br>PTEN (chr10:g.87957955C>T, ENSP00000361021.3:p.Pro246Leu)<br>RB1 (chr13:g.48381415G>A, ENSP00000267163.4:p.Arg556Gln)<br>SETD2 (chr3:g.47121799del, ENSP00000332415.7:p.Glu819ArgfsTer35)<br>SETD7 (chr4:g.139496517G>C, ENSP00000427300.1:p.Gln309Glu)<br>SMARCA4 (chr19:g.10987920T>C, ENSP00000343896.4:p.Tyr372His)<br>SMO (chr7:g.129189225A>G, ENSP00000249373.3:p.Asp25Gly)<br>TSC2 (chr16:g.2060794G>A, ENSP00000219476.3:p.Arg367Gln) | methylated | GBM, IDHwt, RTK I/II | Classic-like | NA |
| T251 | LIH0192 | 43 | Female | Left frontal | recurrence 2 | radiotherapy +TMZ | GBM Grade 4 | GBM, IDHwt, RTK II | ++[EGFR, 2q34], +[1q21.2-24.2, Chr7, Chr19, Chr20], - [9p24.3 -13.3, Chr10], | BCOR (chrX:g.40063031_40063034del, ENSP00000345923.4:p.Leu1262HisfsTer72)<br>GABRA6 (chr5:g.161687516T>C, ENSP00000429943.1:p.Leu13Pro)<br>H2AC17 (chr6:g.27893145G>A, ENSP00000352627.3:p.Ser2Phe)<br>HMCN1 (chr1:g.186093576G>C, ENSP00000271588.4:p.Gly3368Ala)<br>JMJD1C (chr10:g.63193075G>A, ENSP00000335929.5:p.Pro527Leu)<br>MSH2 (chr2:g.47416318G>T, ENSP00000233146.2:p.Gly322Val)<br>SETD7 (chr4:g.139496517G>C, ENSP00000427300.1:p.Gln309Glu)<br>TERT (chr5:g.1254479C>T, ENSP00000309572.5:p.Ala1062Thr)<br>TET1 (chr10:g.68574194G>C, ENSP00000362748.4:p.Cys619Ser) | unmethylated | GBM, IDHwt, RTK I/II | Mesenchymal-like | Proneural |
| T281 | LIH0281 | 51 | Female | Left parietal | primary | no prior treatment | GBM Grade 4 | GBM, IDHwt, mesenchymal | complex genome, +Chr7, - (partial Chr1, 3p, Chr6, Chr9, Chr10, Chr13, Chr14, Chr16, partial Chr17)--CDKN2A/B | FLT3 (chr13:g.28100491T>C, ENSP00000241453.7:p.Asp7Gly)<br>GABRA6 (chr5:g.161687516T>C, ENSP00000429943.1:p.Leu13Pro)<br>H2BC8 (chr6:g.26216619G>A, ENSP00000445633.2:p.Pro9Ser)<br>IDO2 (chr8:g.40015595G>A, ENSP00000443432.2:p.Arg406His)<br>JMJD1C (chr10:g.63215064C>T, ENSP00000382204.2:p.Arg368Gln)<br>KDM2B (chr12:g.121524696G>A, ENSP00000444846.1:p.Ser6Leu)<br>KDM5A (chr12:g.387273A>T, ENSP00000444251.1:p.Phe56Tyr)<br>KMT2C (chr7:g.152149133G>A, ENSP00000262189.6:p.Pro4265Leu)<br>KMT2D (chr12:g.49037164T>C, ENSP00000301067.7:p.Met3398Val)<br>LRP1B (chr2:g.140274452T>A, ENSP00000374135.3:p.Asn4372Tyr)<br>LRP1B (chr2:g.140358851C>T, ENSP00000374135.3:p.Gly3743Ser)<br>LTBP4 (chr19:g.40617142C>T, ENSP00000204005.10:p.Ala1026Val)<br>MSH6 (chr2:g.47799940_47799943dup, ENSP00000234420.5:p.Met654SerfsTer8)<br>PDGFRA (chr4:g.54274864G>C, ENSP00000257290.5:p.Trp559Cys)<br>PTEN (chr10:g.87933012G>A:-:) not pathogenic<br>SETD2 (chr3:g.47122803C>A, ENSP00000332415.7:p.Lys483Asn)<br>SETD7 (chr4:g.139496517G>C, ENSP00000427300.1:p.Gln309Glu) | methylated | GBM, IDHwt, RTK I/II | Classic-like | Mesenchymal |
| T304 | LIH0304 | 53 | Male | Left frontal | primary | no prior treatment | GBM Grade 4 | GBM, IDHwt, RTK II | ++[EGFR, CDK4, MDM2], +[Chr7, Chr20], - (partial Chr1, Chr10, partial Chr12) | ALK (chr2:g.29320870A>G, ENSP00000373700.3:p.Val476Ala)<br>BRPF1 (chr3:g.9742959G>C, ENSP00000373340.2:p.Asp673His)<br>GNAS (chr20:g.58854085C>T, ENSP00000265621.6:p.Pro274Ser)<br>GSE1 (chr16:g.85668334G>T, ENSP00000253458.6:p.Asp1109Tyr)<br>H2BC15 (chr6:g.27839012G>C, ENSP00000475942.1:p.Lys117Asn)<br>KLF7 (chr2:g.207124205C>T, ENSP00000309570.6:p.Arg101Gln)<br>KMT2D (chr12:g.49037164T>C, ENSP00000301067.7:p.Met3398Val)<br>LRP1B (chr2:g.140950336C>A, ENSP00000374135.3:p.Arg1012Ile)<br>MIIP (chr1:g.12022189G>A, ENSP00000235332.4:p.Arg70Gln)<br>PTEN (chr10:g.87933166G>A, ENSP00000361021.3:p.Cys136Tyr)<br>SETD7 (chr4:g.139496517G>C, ENSP00000427300.1:p.Gln309Glu)<br>SMARCD2 (chr17:g.63837555G>A, ENSP00000225742.9:p.Pro21Leu)<br>SMO (chr7:g.129189225A>G, ENSP00000249373.3:p.Asp25Gly)<br>TSC2 (chr16:g.2053484C>T, ENSP00000387937.3:p.Thr18Met) | unmethylated | GBM, IDHwt, RTK I/II | Classic-like | Classical |
| T331 | LIH0331 | 84 | Male | Left fronto-temporo-parietal | primary | no prior treatment | GBM Grade 4 | GBM, IDHwt, mesenchymal | +[Chr 1, Chr 7, Chr 20], -[9p, Chr 10, Chr 22], --[CDKN2A/B, PTEN] | CHD8 (chr14:g.21429308G>A, ENSP00000406288.3:p.Leu12Phe)<br>COL6A3 (chr2:g.237372169_237372171del, ENSP00000295550.4:p.Glu1283del)<br>DNMT1 (chr19:g.10148911G>A, ENSP00000345739.3:p.Thr882Ile)<br>DNMT1 (chr19:g.10175613G>A, ENSP00000345739.3:p.Ala176Val)<br>EGFR (chr7:g.55143387G>A, ENSP00000275493.2:p.Arg108Lys)<br>EGFR (chr7:g.55181309_55181317dup, ENSP00000275493.2:p.Ala767_Val769dup)<br>EZH2 (chr7:g.148836895T>C, ENSP00000417704.1:p.Ile118Val)<br>FGFR3 (chr4:g.1804404T>C, ENSP00000339824.4:p.Phe386Leu)<br>HMCN1 (chr1:g.186000162A>G, ENSP00000271588.4:p.Tyr1331Cys)<br>IDH3B (chr20:g.2658498A>G, ENSP00000370232.5:p.Tyr366His)<br>IDO2 (chr8:g.40005374T>A, ENSP00000443432.2:p.Ser239Thr)<br>KDM4A (chr1:g.43689004C>T, ENSP00000361473.3:p.Pro649Leu)<br>KDM5A (chr12:g.322420T>A, ENSP00000382688.2:p.His808Leu)<br>KMT2D (chr12:g.49051811_49051972del, ENSP00000301067.7:p.Glu578_Glu631del)<br>LTBP4 (chr19:g.40629480G>A, ENSP00000204005.10:p.Cys1565Tyr)<br>MDM4 (chr1:g.204549329A>C, ENSP00000356150.3:p.Lys374Gln)<br>MPL (chr1:g.43338669G>A, ENSP00000361548.3:p.Val114Met)<br>MYB (chr6:g.135202445T>C, ENSP00000326328.8:p.Leu577Pro)<br>MYBL1 (chr8:g.66601730C>T, ENSP00000428265.1:p.Asp56Asn)<br>NOTCH1 (chr9:g.136511217_136511219del, ENSP00000498587.1:p.Gly842del)<br>PIK3C2G (chr12:g.18290875C>A, ENSP00000404845.1:p.Ala261Glu)<br>PMS2 (chr7:g.6006003T>C, ENSP00000265849.7:p.Ile18Val)<br>PPM1D (chr17:g.60656962T>G, ENSP00000376720.3:p.Ter431GlyextTer212)<br>RET (chr10:g.43124887C>T, ENSP00000344798.4:p.Arg982Cys)<br>SETD2 (chr3:g.47057329T>C, ENSP00000386759.3:p.Tyr2152Cys)<br>SETD7 (chr4:g.139496517G>C, ENSP00000427300.1:p.Gln309Glu)<br>TET2 (chr4:g.105234042C>T, ENSP00000265149.5:p.Leu34Phe)<br>ZNF292 (chr6:g.87260932G>A, ENSP00000342847.4:p.Val2430Ile)<br>ZNF814 (chr19:g.57873840G>T, ENSP00000410545.1:p.Ala517Glu) | methylated | GBM, IDHwt, RTK I | Mesenchymal-like | Mesenchymal |
| T341 | LIH0337 | 75 | Female | Right temporal | primary | no prior treatment | GBM Grade 4 | GBM, IDHwt, RTK I | ++[PDGFRA, CDK6], - [3q25.3-q26.2, Chr10 ], --CDKN2A/B | ATM (chr11:g.108244000G>C, ENSP00000278616.4:p.Val182Leu)<br>COL6A3 (chr2:g.237336138T>C, ENSP00000295550.4:p.Met2988Val)<br>FGFR1 (chr8:g.38414029G>A, ENSP00000434800.1:p.Ala131Val)<br>FLT3 (chr13:g.28100491T>C, ENSP00000241453.7:p.Asp7Gly)<br>MPL (chr1:g.43338669G>A, ENSP00000361548.3:p.Val114Met)<br>MUC17 (chr7:g.101033496C>T, ENSP00000302716.4:p.Pro694Ser)<br>MUC17 (chr7:g.101035225C>A, ENSP00000302716.4:p.Thr1270Asn)<br>MYBL1 (chr8:g.66566232T>A, ENSP00000428265.1:p.Arg312Ser)<br>NOTCH2 (chr1:g.120005472C>A, ENSP00000256646.2:p.Arg91Leu)<br>PHC2 (chr1:g.33370563G>A, ENSP00000257118.5:p.Ala145Val)<br>PRKAR1A (chr17:g.68551108G>A, ENSP00000467311.2:p.Ser333Asn)<br>SMARCA2 (chr9:g.2191426G>A, ENSP00000492585.2:p.Trp270Ter)<br>SMO (chr7:g.129189225A>G, ENSP00000249373.3:p.Asp25Gly)<br>TP53 (chr17:g.7673806C>T, ENSP00000269305.4:p.Val272Met) | methylated | GBM, IDHwt, RTK I | Classic-like | NA |
| T347 | LIH0347 | 41 | Male | Left frontal | primary | no prior treatment | GBM Grade 4 | GBM, IDHwt, RTK II | ++EGFR, +[Chr7, 19p, Chr20], - [1p36.3-p34.3, 1q32.3-q44, Chr10, 13q12.11-31.1, 14q11.2-q21.3], --CDKN2A/B | AKAP9 (chr7:g.92017092G>A, ENSP00000348573.3:p.Arg1276Gln)<br>BCL11B (chr14:g.99231420G>A, ENSP00000349723.3:p.Arg189Cys)<br>BRPF1 (chr3:g.9742228del, ENSP00000507969.1:p.Cys687AlafsTer8)<br>COL6A3 (chr2:g.237387630C>T, ENSP00000295550.4:p.Val422Met)<br>CSF1R (chr5:g.150070264C>T, ENSP00000286301.3:p.Gly413Ser)<br>GSE1 (chr16:g.85654423T>G, ENSP00000253458.6:p.Val191Gly)<br>H1-1 (chr6:g.26017622T>A, ENSP00000244573.4:p.Lys37Asn)<br>H2BC1 (chr6:g.25727170C>T, ENSP00000274764.3:p.Arg88Cys)<br>JAK2 (chr9:g.5065003C>G, ENSP00000371067.4:p.Leu393Val)<br>KDM4C (chr9:g.6720968C>T, ENSP00000440656.3:p.Pro7Leu)<br>KDM5A (chr12:g.354122G>A, ENSP00000382688.2:p.Pro328Leu)<br>KDM5A (chr12:g.387273A>T, ENSP00000444251.1:p.Phe56Tyr)<br>KDR (chr4:g.55106779A>G, ENSP00000263923.4:p.Cys482Arg)<br>KMT2C (chr7:g.152180042G>T, ENSP00000262189.6:p.Pro2412Thr)<br>LRP1B (chr2:g.141229297T>G, ENSP00000374135.3:p.Met246Leu)<br>MET (chr7:g.116700208A>G, ENSP00000317272.6:p.Asn375Ser)<br>NOTCH2 (chr1:g.119925578A>T, ENSP00000256646.2:p.Leu1413His)<br>PTEN (chr10:g.87961003del, ENSP00000361021.3:p.Cys304SerfsTer3)<br>SETD7 (chr4:g.139496517G>C, ENSP00000427300.1:p.Gln309Glu) | unmethylated | GBM, IDHwt, RTK I/II | Classic-like | Classical |
| T356 | LIH0281 | 52 | Female | Left parietal, para-sagittal | recurrence 1 | radiotherapy +TMZ | GBM Grade 4 | GBM, IDHwt, mesenchymal | complex genome, +Chr7, -(partial Chr1, partial Chr2, partial Chr3, Chr4, Chr6, Chr10, Chr14, Ch17, Chr18) | FLT3 (chr13:g.28100491T>C, ENSP00000241453.7:p.Asp7Gly)<br>GABRA6 (chr5:g.161687516T>C, ENSP00000429943.1:p.Leu13Pro)<br>H2BC8 (chr6:g.26216619G>A, ENSP00000445633.2:p.Pro9Ser)<br>JMJD1C (chr10:g.63215064C>T, ENSP00000382204.2:p.Arg368Gln)<br>KDM2B (chr12:g.121524696G>A, ENSP00000444846.1:p.Ser6Leu)<br>KMT2C (chr7:g.152149133G>A, ENSP00000262189.6:p.Pro4265Leu)<br>KMT2D (chr12:g.49037164T>C, ENSP00000301067.7:p.Met3398Val)<br>LRP1B (chr2:g.140274452T>A, ENSP00000374135.3:p.Asn4372Tyr)<br>LRP1B (chr2:g.140358851C>T, ENSP00000374135.3:p.Gly3743Ser)<br>LTBP4 (chr19:g.40613152C>T, ENSP00000204005.10:p.Ala826Val)<br>LTBP4 (chr19:g.40617142C>T, ENSP00000204005.10:p.Ala1026Val)<br>NF1 (chr17:g.31327521_31327537del, ENSP00000348498.3:p.Val1743AspfsTer12)<br>PCDH8 (chr13:g.52846715T>A, ENSP00000341350.4:p.Gln574His)<br>PDGFRA (chr4:g.54274864G>C, ENSP00000257290.5:p.Trp559Cys)<br>PIK3CA (chr3:g.179234297A>G, ENSP00000263967.3:p.His1047Arg)<br>SETD2 (chr3:g.47122803C>A, ENSP00000332415.7:p.Lys483Asn)<br>SETD7 (chr4:g.139496517G>C, ENSP00000427300.1:p.Gln309Glu)<br>TRAF7 (chr16:g.2168184C>T, ENSP00000478515.1:p.Gln83Ter)<br>ZFTA (chr11:g.63765068T>C, ENSP00000482180.1:p.Asp275Gly) | unmethylated | GBM, IDHwt, RTK I/II | Mesenchymal-like | Classical |
| T361 | LIH0361 | 69 | Female | Left temporal, para-brainstem | NA | no prior treatment | GBM Grade 4 | GBM, IDHwt, RTK II | ++(EGFR, MDM4) + [Chr7, 19p, 20p13-p11.1, 20q13.1-q13.3], - [3q13.33-q21.1, 6p25.3-p21.32, 10q23.2-q26.3], --CDKN2A/B | APC (chr5:g.112838980T>C, ENSP00000257430.4:p.Leu1129Ser)<br>ARID2 (chr12:g.45776079A>G:-:)<br>CIC (chr19:g.42290992C>T, ENSP00000160740.3:p.Pro742Ser)<br>COL6A3 (chr2:g.237336138T>C, ENSP00000295550.4:p.Met2988Val)<br>DNMT3B (chr20:g.32795432G>A, ENSP00000201963.3:p.Ala376Thr)<br>EGFR (chr7:g.55143387G>A, ENSP00000275493.2:p.Arg108Lys)<br>FLT3 (chr13:g.28100491T>C, ENSP00000241453.7:p.Asp7Gly)<br>JAK2 (chr9:g.5065003C>G, ENSP00000371067.4:p.Leu393Val)<br>JMJD1C (chr10:g.63177741G>C, ENSP00000335929.5:p.Asp947Glu)<br>KDM5A (chr12:g.387273A>T, ENSP00000444251.1:p.Phe56Tyr)<br>KDR (chr4:g.55106779A>G, ENSP00000263923.4:p.Cys482Arg)<br>LRP1B (chr2:g.140356425G>A, ENSP00000374135.3:p.Ala3816Val)<br>LRP1B (chr2:g.140358878G>T, ENSP00000374135.3:p.Gln3734Lys)<br>MUC17 (chr7:g.101033503A>G, ENSP00000302716.4:p.Asn696Ser)<br>MYB (chr6:g.135202445T>C, ENSP00000326328.8:p.Leu577Pro)<br>RAD50 (chr5:g.132587981G>T, ENSP00000368100.4:p.Val315Leu)<br>SETD1B (chr12:g.121823041G>A, ENSP00000442924.1:p.Ala1445Thr)<br>SETD7 (chr4:g.139496517G>C, ENSP00000427300.1:p.Gln309Glu)<br>TET1 (chr10:g.68573641G>A, ENSP00000362748.4:p.Val435Ile)<br>TET2 (chr4:g.105234042C>T, ENSP00000265149.5:p.Leu34Phe)<br>TET2 (chr4:g.105237262C>T, ENSP00000265149.5:p.Ser1107Leu) | methylated | GBM, IDHwt, RTK II | Classic-like | Classical |

| | | | | | | | | | | | | | | |
|---|---|---|---|---|---|---|---|---|---|---|---|---|---|---|
| T363 | LIH0363 | 84 | Female | Right occipital lobe | primary | no prior treatment | GBM Grade 4 | NA | complex genome ++EGFR,-[Chr 1, 9p, Chr10, 15q, -CDKN2A/B] | APC (chr5:g.112843351G>T, ENSP00000257430.4:p.Ser2586Ile)<br>BRCA1 (chr17:g.43070959A>G, ENSP00000312236.5:p.Met510Thr)<br>CSF1R (chr5:g.150070264C>T, ENSP0000286301.3:p.Gly413Ser)<br>GSE1 (chr16:g.85656394_85656399del, ENSP00000253458.6:p.Arg351_Glu352del)<br>HDAC9 (chr7:g.18591614G>C, ENSP00000383912.1:p.Val172Leu)<br>HMCN1 (chr1:g.185922425C>A, ENSP00000271588.4:p.Thr316Asn)<br>IDO2 (chr8:g.40005374T>A, ENSP00000443432.2:p.Ser239Thr)<br>IGF1R (chr15:g.98939285G>A, ENSP00000496919.1:p.Ala1127Thr)<br>JMJD1C (chr10:g.63193075G>A, ENSP00000335929.5:p.Pro527Leu)<br>KDM2B (chr12:g.121510033G>T, ENSP0000366269.3:p.Pro363Gln)<br>KMT2C (chr7:g.152180042G>T, ENSP00000262189.6:p.Pro2412Thr)<br>L2HGDH (chr14:g.50237650G>C, ENSP00000261699.4:p.Phe424Leu)<br>LRP1B (chr2:g.140700613C>A, ENSP00000374135.3:p.Val2146Phe)<br>LTBP4 (chr19:g.40610549C>G, ENSP00000204005.10:p.Arg598Gly)<br>LTBP4 (chr19:g.40627100G>A, ENSP00000204005.10:p.Gly1401Arg)<br>MET (chr7:g.116771869C>T, ENSP00000317272.6:p.Arg988Cys)<br>MRE11 (chr11:g.94471601G>C, ENSP00000325863.4:p.Ser273Cys)<br>PCDH8 (chr13:g.52846209A>G, ENSP00000341350.4:p.Val743Ala)<br>PCDH8 (chr13:g.52847338T>C, ENSP00000341350.4:p.Thr367Ala)<br>RET (chr10:g.43128249A>G, ENSP0000347942.3:p.Met1109Val)<br>SETD7 (chr4:g.139496517G>C, ENSP00000427300.1:p.Gln309Glu)<br>ZNF292 (chr6:g.87249373G>T, ENSP00000428073.2:p.Arg344Leu) | methylated | GBM, IDHwt, RTK I/II | Classic-like | NA |
| T367 | LIH0367 | 69 | Male | Left temporal | primary | no prior treatment | GBM Grade 4 | GBM, IDHwt, RTK II | ++[EGFR, MDM4], +[Chr7, Chr19], - Chr10 | ATM (chr11:g.108272729C>G, ENSP00000278616.4:p.Pro1054Arg)<br>CDKN2A (chr9:g.21971209C>T:-:)<br>CIC (chr19:g.42293072G>A, ENSP00000160740.3:p.Ala1194Thr)<br>COL6A3 (chr2:g.237387744C>T, ENSP00000295550.4:p.Ala384Thr)<br>DDX3X (chrX:g.41346608G>A, ENSP0000382840.3:p.Arg533His)<br>DNMT3B (chr20:g.32795432G>A, ENSP00000201963.3:p.Ala376Thr)<br>GABRA6 (chr5:g.161687516T>C, ENSP00000429943.1:p.Leu13Pro)<br>IDH2 (chr15:g.90102311G>A, ENSP00000331897.4:p.Thr27Ile)<br>KDM4D (chr11:g.94998658C>G, ENSP00000334181.5:p.Pro429Arg)<br>KDR (chr4:g.55102380T>C, ENSP00000263923.4:p.Thr706Ala)<br>KMT2C (chr7:g.152177496G>C, ENSP00000262189.6:p.Leu2653Val)<br>LRP1B (chr2:g.140314950C>G, ENSP00000374135.3:p.Val4264Leu)<br>LTBP4 (chr19:g.40627020G>T, ENSP00000204005.10:p.Ser1374Ile)<br>LTBP4 (chr19:g.40627287A>T, ENSP00000204005.10:p.Tyr1463Phe)<br>MET (chr7:g.116700208A>G, ENSP00000317272.6:p.Asn375Ser)<br>MUC17 (chr7:g.101048035C>T, ENSP00000302716.4:p.Thr4152Ile)<br>NOTCH1 (chr9:g.136505767G>A, ENSP00000498587.1:p.Pro1377Ser)<br>PDGFRA (chr4:g.54267424G>A, ENSP00000257290.5:p.Val299Ile)<br>PMS2 (chr7:g.5986899C>T, ENSP00000265849.7:p.Met622Ile)<br>PTEN (chr10:g.87957873C>T, ENSP00000361021.3:p.Gln219Ter)<br>SETD2 (chr3:g.47106044G>A, ENSP00000386759.3:p.Arg1598Ter)<br>SETD7 (chr4:g.139496517G>C, ENSP00000427300.1:p.Gln309Glu)<br>TET2 (chr4:g.105275923A>T, ENSP00000369351.4:p.Asn1805Tyr)<br>ZNF814 (chr19:g.57873013_57873096del, ENSP00000410545.1:p.Phe787_Ser814del) | methylated | GBM, IDHwt, RTK I/II | Classic-like | NA |
| T384 | LIH0384 | 50 | Female | Left temporo-occipital | primary | no prior treatment | GBM Grade 4 | GBM, IDHwt, mesenchymal (no match) | ++[EGFR, 7p15.2-p11.2, 7q31.31-32.2], -[4p, 9q, 10q, partial Chr 12, partial Chr 15], --[FGFR1\TACC1, MYBL1] | ALK (chr2:g.29320870A>G, ENSP00000373700.3:p.Val476Ala)<br>ATR (chr3:g.142547733C>T, ENSP00000343741.4:p.Glu1117Lys)<br>AURKA (chr20:g.56370253T>C, ENSP00000321591.6:p.Met373Val)<br>HMCN1 (chr1:g.186038858C>T, ENSP00000271588.4:p.Leu1961Phe)<br>KDM5A (chr12:g.387273A>T, ENSP00000444251.1:p.Phe56Tyr)<br>LRP1B (chr2:g.140776142G>A, ENSP00000374135.3:p.Ser1819Phe)<br>MUC17 (chr7:g.101043496C>T, ENSP00000302716.4:p.Thr4027Met)<br>MYB (chr6:g.135198939C>T, ENSP00000326328.8:p.Ser412Leu)<br>MYBL1 (chr8:g.66566185T>C, ENSP00000428265.1:p.Asn328Ser)<br>NOTCH1 (chr9:g.136505767G>A, ENSP00000498587.1:p.Pro1377Ser)<br>SETD7 (chr4:g.139496517G>C, ENSP00000427300.1:p.Gln309Glu)<br>SMO (chr7:g.129205287G>A, ENSP00000249373.3:p.Glu208Lys)<br>SMO (chr7:g.129205670G>A, ENSP00000249373.3:p.Val270Ile)<br>TP53 (chr17:g.7674257A>T, ENSP00000269305.4:p.Tyr236Asn)<br>TSC2 (chr16:g.2060794G>A, ENSP00000219476.3:p.Arg367Gln) | methylated | GBM, IDHwt, other (no match: RTK II) | Mesenchymal-like | Mesenchymal |
| T386 | LIH0386 | 51 | Male | Left frontal | primary | no prior treatment | GBM Grade 4 | NA | +[Chr7, 19p], - [1p36.3-p34.3, Chr10, 11q12.3-q13.2, 11q24.1-q13.2, 14q11.2-q24.1, 17q11.2-q21.2, 18p11.32-p11.22, 22p ], --[CDKN2A/B, PTEN] | ALK (chr2:g.29320870A>G, ENSP00000373700.3:p.Val476Ala)<br>ATR (chr3:g.142515493T>C, ENSP00000343741.4:p.Thr1469Ala)<br>CHEK2 (chr22:g.28694073G>A, ENSP00000329012.5:p.Arg445Cys)<br>FLT3 (chr13:g.28100491T>C, ENSP00000241453.7:p.Asp7Gly)<br>HMCN1 (chr1:g.186119233G>A, ENSP00000271588.4:p.Arg3964Lys)<br>HNF1A (chr12:g.120996336G>C, ENSP00000257555.5:p.Val344Leu)<br>KMT2C (chr7:g.152252650T>C, ENSP00000262189.6:p.Ile455Met)<br>LZTR1 (chr22:g.20996745C>T, ENSP00000496779.1:p.Gln757Ter)<br>MPL (chr1:g.43338669G>A, ENSP00000361548.3:p.Val114Met)<br>MRE11 (chr11:g.94447335T>C, ENSP00000325863.4:p.Asn556Ser)<br>MUC17 (chr7:g.101041242_101041243delinsTA, ENSP00000302716.4:p.Pro3276Tyr)<br>NF1 (chr17:g.31221932A>G, ENSP00000412921.4:p.Tyr575Cys)- not pathogenic<br>PIK3R1 (chr5:g.68293801_68293803del, ENSP00000323512.8:p.Asp164_Arg165delinsGlu)<br>SETD2 (chr3:g.47103362T>C, ENSP00000386759.3:p.Asn1634Ser) | unmethylated | GBM, IDHwt, RTK I/II | Classic-like | NA |
| T394 | LIH0394 | 45 | Female | Right fronto-basal | recurrence 1 | radiotherapy | GBM Grade 4 (Secondary) | Glioma, IDHm, high-grade astrocytoma | complex genome ++[PDGFRA,FGFR3/TACC3, KIT, KDR, MET, Chr 4, 7q31.31], --CDKN2A/B | ATRX (chrX:g.77681626_77681628del, ENSP00000362441.4:p.Asp1212del)<br>ATRX (chrX:g.77682604_77682607del, ENSP00000362441.4:p.Glu886LeufsTer18)<br>HMCN1 (chr1:g.186174588C>T, ENSP00000271588.4:p.Arg5297Cys)<br>IDH1 (chr2:g.208248388C>T, ENSP00000260985.2:p.Arg132His)<br>JAK3 (chr19:g.17838345G>A, ENSP00000391676.1:p.Thr496Ile)<br>JAK3 (chr19:g.17843141G>C, ENSP00000391676.1:p.Pro151Arg)<br>MET (chr7:g.116700208A>G, ENSP00000317272.6:p.Asn375Ser)<br>MUC17 (chr7:g.101032659A>G, ENSP00000302716.4:p.Thr415Ala)<br>MYBL1 (chr8:g.66566196T>G, ENSP00000428265.1:p.Glu324Asp)<br>NTRK2 (chr9:g.84948595T>A, ENSP00000277120.3:p.Phe633Tyr)<br>NUP210L (chr1:g.153995075_153995078del:-:)<br>PIK3C2G (chr12:g.18290875C>A, ENSP00000404845.1:p.Ala261Glu)<br>PRKAR1A (chr17:g.68551108G>A, ENSP00000467311.2:p.Ser333Asn)<br>TP53 (chr17:g.7670697_7670707del, ENSP00000269305.4:p.Glu336AspfsTer7) | methylated | Glioma, IDHm, high-grade astrocytoma | K1: G-CIMP-low | Proneural |
| T407 | LIH0394 | 45 | Female | Right fronto-basal | recurrence 1 | radiotherapy | GBM Grade 4 (Secondary) | Glioma, IDHm, high grade astrocytoma | complex genome ++[PDGFRA,FGFR3/TACC3, KIT, KDR, MET, Chr 4, 7q31.31], --CDKN2A/B | ARID1B (chr6:g.156779448G>A, ENSP00000344546.5:p.Gly590Arg)<br>ATRX (chrX:g.77681626_77681628del, ENSP00000362441.4:p.Asp1212del)<br>ATRX (chrX:g.77682604_77682607del, ENSP00000362441.4:p.Glu886LeufsTer18)<br>HMCN1 (chr1:g.186174588C>T, ENSP00000271588.4:p.Arg5297Cys)<br>IDH1 (chr2:g.208248388C>T, ENSP00000260985.2:p.Arg132His)<br>JAK3 (chr19:g.17838345G>A, ENSP00000391676.1:p.Thr496Ile)<br>JAK3 (chr19:g.17843141G>C, ENSP00000391676.1:p.Pro151Arg)<br>MET (chr7:g.116700208A>G, ENSP00000317272.6:p.Asn375Ser)<br>MUC17 (chr7:g.101032659A>G, ENSP00000302716.4:p.Thr415Ala)<br>MYBL1 (chr8:g.66566196T>G, ENSP00000428265.1:p.Glu324Asp)<br>NUP210L (chr1:g.153995075_153995078del:-:)<br>OLIG2 (chr21:g.33027730G>T, ENSP00000331040.3:p.Gly290Cys)<br>PIK3C2G (chr12:g.18290875C>A, ENSP00000404845.1:p.Ala261Glu)<br>PRKAR1A (chr17:g.68551108G>A, ENSP00000467311.2:p.Ser333Asn)<br>TP53 (chr17:g.7670697_7670707del, ENSP00000269305.4:p.Glu336AspfsTer7) | methylated | Glioma, IDHm, high-grade astrocytoma | K1: G-CIMP-low | Proneural |
| T434 | LIH0304 | 54 | Male | Left frontal | recurrence 2 | radiotherapy + TMZ | GBM Grade 4 | GBM, IDHwt, RTK II | ++[EGFR, CDK4, MDM2], +[7p,7q36.1-q36.2, Chr20], -[1p36.23-p36.21, 6q26-q27, Chr10, 12q15, 14q13-qter] | ALK (chr2:g.29320870A>G, ENSP00000373700.3:p.Val476Ala)<br>ARID1A (chr1:g.26766351_26766373del:-:)<br>ARID1B (chr6:g.157207854T>C, ENSP00000344546.5:p.Val2321Ala)<br>CCND2 (chr12:g.4299904_4299952dup, ENSP00000261254.3:p.Glu272GlnfsTer2)<br>EGFR (chr7:g.55154023T>A, ENSP00000275493.2:p.Phe254Ile)<br>EPHA7 (chr6:g.93419273G>T, ENSP00000358303.1:p.His23Gln)<br>GNAS (chr20:g.58854085C>T, ENSP00000265621.6:p.Pro274Ser)<br>GSE1 (chr16:g.85672270C>T, ENSP00000410812.1:p.Thr405Ile)<br>H2BC15 (chr6:g.27839012G>C, ENSP00000475942.1:p.Lys117Asn)<br>KLF7 (chr2:g.207124205C>T, ENSP00000309570.6:p.Arg101Gln)<br>KMT2D (chr12:g.49037164T>C, ENSP00000301067.7:p.Met3398Val)<br>LRP1B (chr2:g.140950336C>A, ENSP00000374135.3:p.Arg1012Ile)<br>MIIP (chr1:g.12022189G>A, ENSP00000235332.4:p.Arg70Gln)<br>PIK3C2G (chr12:g.18609549G>C, ENSP00000404845.1:p.Asp1327His)<br>PTEN (chr10:g.87933166G>A, ENSP00000361021.3:p.Cys136Tyr)<br>SETD7 (chr4:g.139496517G>C, ENSP00000427300.1:p.Gln309Glu)<br>SIRPB1 (chr20:g.1572020C>T, ENSP00000371018.5:p.Val151Met)<br>SMARCD2 (chr17:g.63837555G>A, ENSP00000225742.9:p.Pro21Leu)<br>SMO (chr7:g.129189225A>G, ENSP00000249373.3:p.Asp25Gly)<br>TSC2 (chr16:g.2082582G>A, ENSP00000495110.1:p.Ala39Thr) | unmethylated | GBM, IDHwt, RTK II | Classic-like | Proneural |
| T470 | LIH0347 | 42 | Male | Left frontal | recurrence 1 | radiotherapy + TMZ | GBM Grade 4 | GBM, IDHwt, RTK II | ++EGFR, +[Chr7, 19p, Chr20], - [ 1p36.3-p34.3, 1p31.1-p11.21, 1q32.3-q44, Chr10, 13q12.11-31.1, 14q11.2-q21.3], --CDKN2A/B | AKAP9 (chr7:g.92017092G>A, ENSP00000348573.3:p.Arg1276Gln)<br>BCL11B (chr14:g.99231420G>A, ENSP00000349723.3:p.Arg189Cys)<br>BRPF1 (chr3:g.9742228del, ENSP00000507969.1:p.Cys687AlafsTer8)<br>COL6A3 (chr2:g.237387630C>T, ENSP00000295550.4:p.Val422Met)<br>CSF1R (chr5:g.150070264C>T, ENSP0000286301.3:p.Gly413Ser)<br>GSE1 (chr16:g.85654423T>G, ENSP00000253458.6:p.Val191Gly)<br>H1-1 (chr6:g.26017622T>A, ENSP00000244573.4:p.Lys37Asn)<br>H2BC1 (chr6:g.25727170C>T, ENSP00000274764.3:p.Arg88Cys)<br>JAK2 (chr9:g.5065003C>G, ENSP00000371067.4:p.Leu393Val)<br>KDM4C (chr9:g.6720968C>T, ENSP00000440656.3:p.Pro7Leu)<br>KDM5A (chr12:g.354122G>A, ENSP00000382688.2:p.Pro328Leu)<br>KDM5A (chr12:g.387273A>T, ENSP00000444251.1:p.Phe56Tyr)<br>KDR (chr4:g.55106779A>G, ENSP00000263923.4:p.Cys482Arg)<br>KMT2C (chr7:g.152180042G>T, ENSP00000262189.6:p.Pro2412Thr)<br>LRP1B (chr2:g.141229297T>G, ENSP00000374135.3:p.Met246Leu)<br>MET (chr7:g.116700208A>G, ENSP00000317272.6:p.Asn375Ser)<br>MUC17 (chr7:g.101039898G>A, ENSP00000302716.4:p.Gly2828Ser)<br>MUC17 (chr7:g.101040303G>A, ENSP00000302716.4:p.Ala2963Thr)<br>NOTCH2 (chr1:g.119925578A>T, ENSP00000256646.2:p.Leu1413His)<br>PTEN (chr10:g.87961003del, ENSP00000361021.3:p.Cys304SerfsTer3)<br>SETD7 (chr4:g.139496517G>C, ENSP00000427300.1:p.Gln309Glu)<br>ZNF814 (chr19:g.57874431C>T, ENSP00000410545.1:p.Gly320Glu) | unmethylated | GBM, IDHwt, RTK II | Classic-like | Classical |
| T476 | LIH0476 | 75 | Male | Left fronto-parietal | primary | no prior treatment | GBM Grade 4 | GBM, IDHwt, RTK I/II | ++[EGFR, MDM4], +[Chr 7], - [9p24.3-p21.3, Chr 10], --[CDKN2A/B, PTEN] | AKAP9 (chr7:g.92065429A>G, ENSP00000348573.3:p.Glu2059Gly)<br>CIC (chr19:g.42290992C>T, ENSP00000160740.3:p.Pro742Ser)<br>EGFR (chr7:g.55165350G>T, ENSP00000275493.2:p.Gly598Val)<br>EGFR (chr7:g.55200333G>C, ENSP00000275493.2:p.Asp956His)<br>GABRA6 (chr5:g.161687516T>C, ENSP00000429943.1:p.Leu13Pro)<br>GSE1 (chr16:g.85656394_85656399del, ENSP00000253458.6:p.Arg351_Glu352del)<br>H3C6 (chr6:g.26225537C>T, ENSP00000482271.1:p.Ala128Val)<br>MET (chr7:g.116771936C>T, ENSP00000317272.6:p.Thr1010Ile)<br>MSH6 (chr2:g.47798646A>C, ENSP00000234420.5:p.Glu221Asp)<br>NOTCH1 (chr9:g.136506781C>T, ENSP00000498587.1:p.Arg1279His)<br>PCDH8 (chr13:g.52846516C>T, ENSP00000341350.4:p.Ala641Thr)<br>SMARCA2 (chr9:g.2191426G>A, ENSP00000492585.2:p.Trp270Ter)<br>TERT (chr5:g.1293569_1293571del, ENSP00000309572.5:p.Glu441del)<br>TET2 (chr4:g.105237193A>C, ENSP00000265149.5:p.Gln1084Pro) | unmethylated | GBM, IDHwt, RTK I | Mesenchymal-like | Proneural |
| T515 | LIH0515 | 46 | Male | Right frontal | primary | no prior treatment | Anaplastic Oligodendroglioma Grade 3 | NA | NA | NA | NA | NA | NA | NA |
| T591 | LIH0384 | 52 | Female | Right temporo-occipital | recurrence 1 | radiotherapy +TMZ + Avastin + Irinotecan | GBM Grade 4 | GBM, IDHwt, RTK II/mesenchymal | ++[EGFR, 7p15.2-p11.2, 7q31.31-32.2], -[1p21.1, 3q26.1, 8p, 9p24.3, Chr 10, 11q24.2, 13q21.33-q31.1, 15q] | ALK (chr2:g.29320870A>G, ENSP00000373700.3:p.Val476Ala)<br>AURKA (chr20:g.56370253T>C, ENSP00000321591.6:p.Met373Val)<br>FLT3 (chr13:g.28100491T>C, ENSP00000241453.7:p.Asp7Gly)<br>HDAC9 (chr7:g.18954224A>G, ENSP00000383912.1:p.Ile962Val)<br>HMCN1 (chr1:g.186038858C>T, ENSP00000271588.4:p.Leu1961Phe)<br>LRP1B (chr2:g.140776142G>A, ENSP00000374135.3:p.Ser1819Phe)<br>MUC17 (chr7:g.101043496C>T, ENSP00000302716.4:p.Thr4027Met)<br>MYB (chr6:g.135198939C>T, ENSP00000326328.8:p.Ser412Leu)<br>MYBL1 (chr8:g.66566185T>C, ENSP00000428265.1:p.Asn328Ser)<br>NOTCH1 (chr9:g.136505767G>A, ENSP00000498587.1:p.Pro1377Ser)<br>SETD7 (chr4:g.139496517G>C, ENSP00000427300.1:p.Gln309Glu)<br>SMO (chr7:g.129205287G>A, ENSP00000249373.3:p.Glu208Lys)<br>SMO (chr7:g.129205670G>A, ENSP00000249373.3:p.Val270Ile)<br>TSC2 (chr16:g.2060794G>A, ENSP00000219476.3:p.Arg367Gln) | unmethylated | GBM, IDHwt, RTK I/II | Classic-like | Mesenchymal |
| T744 | LIH0744 | 69 | Male | Bilateral involvement of the hippocampus | primary | no prior treatment | GBM Grade 4 | NA | +Chr 7, -[6q, 9p, Chr10, Chr 11, 14q, 22q], --CDKN2A/B | H2BC11 (chr6:g.27132726G>C, ENSP00000342886.3:p.Pro9Ala)<br>H4C5 (chr6:g.26204726C>T, ENSP00000484789.1:p.Gln28Ter)<br>JMJD1C (chr10:g.63207223del, ENSP00000335929.5:p.Lys170SerfsTer24)<br>KDM5A (chr12:g.387273A>T, ENSP00000444251.1:p.Phe56Tyr)<br>KDM6B (chr17:g.7849900_7849902del, ENSP00000254846.5:p.Glu1174del)<br>KMT2D (chr12:g.49051644G>A, ENSP00000301067.7:p.Thr680Met)<br>LTBP4 (chr19:g.40605073G>C, ENSP00000204005.10:p.Val127Leu)<br>MUC17 (chr7:g.101041583A>G, ENSP00000302716.4:p.Ile3389Met)<br>NF1 (chr17:g.31338092C>T, ENSP00000348498.3:p.Arg2237Ter)<br>PAX7 (chr1:g.18635124C>T, ENSP00000364524.3:p.Pro112Leu)<br>PRKAR1A (chr17:g.68551108G>A, ENSP00000467311.2:p.Ser333Asn)<br>SETD7 (chr4:g.139496517G>C, ENSP00000427300.1:p.Gln309Glu)<br>ZNF814 (chr19:g.57873013_57873096del, ENSP00000410545.1:p.Phe787_Ser814del) | unmethylated | GBM, IDHwt, RTK I | Classic-like | NA |

| | | | | | | | | | | | | | | |
|---|---|---|---|---|---|---|---|---|---|---|---|---|---|---|
| T756 | LIH0556 | 46 | Male | Multifocal left | recurrence 3 | radiotherapy | GBM Grade 4 (Secondary) | Glioma, IDHm, high-grade astrocytoma | + [4q12, 10p], -[4p15.32-q35.2, 6q27, 10q, partial Chr 16, 22q], --CDKN2A/B | ATM (chr11:g.108304736A>T, ENSP00000278616.4:p.Asp1853Val)<br>D2HGDH (chr2:g.241755984G>A, ENSP00000315351.4:p.Ala426Thr)<br>DICER1 (chr14:g.95091293C>T, ENSP00000343745.3:p.Glu1813Lys)<br>FLT3 (chr13:g.28049450C>T, ENSP00000241453.7:p.Asp324Asn)<br>FLT3 (chr13:g.28100491T>C, ENSP00000241453.7:p.Asp7Gly)<br>IDH1 (chr2:g.208248388C>T, ENSP00000260985.2:p.Arg132His)<br>IDO2 (chr8:g.40005374T>A, ENSP00000443432.2:p.Ser239Thr)<br>JMJD1C (chr10:g.63214218G>A, ENSP00000382204.2:p.Thr650Ile)<br>KDM4A (chr1:g.43669216T>C, ENSP00000361473.3:p.Met427Thr)<br>KMT2D (chr12:g.49049949C>G, ENSP00000301067.7:p.Gln1213His)<br>KMT2D (chr12:g.49050218G>A, ENSP00000301067.7:p.Pro1124Ser)<br>MSH6 (chr2:g.47800651G>T, ENSP00000234420.5:p.Val890Phe)<br>MUC17 (chr7:g.101033386_101033562del, ENSP00000302716.4:p.Asn657_Thr715del)<br>NUP210L (chr1:g.154100125G>A, ENSP00000271854.3:p.Pro613Leu)<br>PIK3C2G (chr12:g.18648027G>C, ENSP00000404845.1:p.Ter1446SerextTer4)<br>PTCH1 (chr9:g.95469004G>A, ENSP00000332353.6:p.Thr666Met)<br>SETD7 (chr4:g.139496517G>C, ENSP00000427300.1:p.Gln309Glu)<br>TET2 (chr4:g.105237403A>G, ENSP00000306705.2:p.Asn1154Ser)<br>TP53 (chr17:g.7675085C>T, ENSP00000269305.4:p.Cys176Tyr)<br>TRDMT1 (chr10:g.17153494G>A, ENSP00000419625.1:p.Thr240Met)<br>VHL (chr3:g.10141921C>T, ENSP00000256474.3:p.Pro25Leu)<br>ZEB1 (chr10:g.31521751A>G, ENSP00000319248.9:p.Thr806Ala) | methylated | Glioma, IDHm, high grade astrocytoma | K1: G-CIMP-low | Proneural |
| T772 | LIH0615 | 54 | Female | Fronto-temporal left | recurrence 1 | radiotherapy + TMZ | GBM Grade 4 | GBM, IDHwt, RTK II | ++EGFR, +[7p, Chr19], -[1p36.3, 9p, Chr 10, 15q] --CDKN2A/B | AMER1 (chrX:g.64192617G>C, ENSP00000364003.4:p.Pro224Ala)<br>ARID1A (chr1:g.26762287A>G, ENSP00000320485.7:p.Tyr796Cys)<br>ARID1B (chr6:g.156778313_156778315dup, ENSP00000344546.5:p.Gln214dup)<br>ARID1B (chr6:g.157200730T>C, ENSP00000344546.5:p.Met1462Thr)<br>CDH1 (chr16:g.68738336C>A, ENSP00000261769.4:p.Pro30Thr)<br>COL6A3 (chr2:g.237336138T>C, ENSP00000295550.4:p.Met2988Val)<br>DNMT1 (chr19:g.10180797C>T, ENSP00000345739.3:p.Arg69His)<br>EGFR (chr7:g.55165350G>T, ENSP00000275493.2:p.Gly598Val)<br>FLT3 (chr13:g.28100491T>C, ENSP00000241453.7:p.Asp7Gly)<br>GNAS (chr20:g.58855063C>G, ENSP00000265621.6:p.Arg600Gly)<br>GSE1 (chr16:g.85634010A>G, ENSP00000253458.6:p.Asn35Ser)<br>H2BC4 (chr6:g.26123679C>T, ENSP00000321744.4:p.Gly76Ser)<br>IDH2 (chr15:g.90084321G>A, ENSP00000331897.4:p.Thr435Met)<br>KDM2B (chr12:g.121580856T>C, ENSP00000366271.3:p.His19Arg)<br>KDR (chr4:g.55106779A>G, ENSP00000263923.4:p.Cys482Arg)<br>KLK1 (chr19:g.50821806A>G, ENSP00000301420.1:p.Trp38Arg)<br>KMT2A (chr11:g.118481850G>A, ENSP00000374157.5:p.Ser1257Asn)<br>KMT2C (chr7:g.152180042G>T, ENSP00000262189.6:p.Pro2412Thr)<br>LRP1B (chr2:g.140526272A>G, ENSP00000374135.3:p.Ile2614Thr)<br>MSH2 (chr2:g.47512394G>A, ENSP00000384199.1:p.Arg909Gln)<br>MUC17 (chr7:g.101038698_101038723del, ENSP00000302716.4:p.Val2428IlefsTer6)<br>NBN (chr8:g.89978293T>C, ENSP00000265433.4:p.Ile171Val)<br>NCOR1 (chr17:g.16047086C>T, ENSP00000268712.2:p.Ala2182Thr)<br>NUP210L (chr1:g.154046179C>T, ENSP00000271854.3:p.Val1196Ile)<br>PHC2 (chr1:g.33370563G>A, ENSP00000257118.5:p.Ala145Val)<br>PTEN (chr10:g.87961115del, ENSP00000361021.3:p.Phe341LeufsTer3)<br>PTGFR (chr1:g.78492772T>C, ENSP00000359792.3:p.Val10Ala)<br>RET (chr10:g.43124887C>T, ENSP00000344798.4:p.Arg982Cys)<br>SETD7 (chr4:g.139496517G>C, ENSP00000427300.1:p.Gln309Glu)<br>TET2 (chr4:g.105235030C>T, ENSP00000265149.5:p.Pro363Leu)<br>TSC2 (chr16:g.2048590_2048591delinsAA, ENSP00000454487.1:p.Gly3Glu) | methylated | GBM, IDHwt, RTK I | Classic-like | NA |
| T784 | LIH0784 | 56 | Male | Left frontal | primary | no prior treatment | GBM Grade 4 | GBM, IDHwt, RTK I | +[Chr 7], -[1p36.3-22.1, 3p12.1, 10q], --CDKN2A/B | AKAP9 (chr7:g.92001112G>A, ENSP00000348573.3:p.Gly399Arg)<br>ALK (chr2:g.29320870A>G, ENSP00000373700.3:p.Val476Ala)<br>BRPF1 (chr3:g.9742228del, ENSP00000507969.1:p.Cys687AlafsTer8)<br>CREB1 (chr2:g.207575323G>A, ENSP00000236995.3:p.Gly186Glu)<br>HMCN1 (chr1:g.186055511G>A, ENSP00000271588.4:p.Met2327Ile)<br>HMCN1 (chr1:g.186122971C>T, ENSP00000271588.4:p.His4084Tyr)<br>HMCN1 (chr1:g.186144595G>A, ENSP00000271588.4:p.Ala4720Thr)<br>IGF1R (chr15:g.98916049C>T, ENSP00000456950.1:p.Arg49Trp)<br>KDM1A (chr1:g.23019732G>A, ENSP00000349049.3:p.Gly46Ser)<br>MET (chr7:g.116695757A>T, ENSP00000413857.1:p.Met1?)<br>MSH6 (chr2:g.47799169C>G, ENSP00000234420.5:p.Leu396Val)<br>MUC17 (chr7:g.101033503A>G, ENSP00000302716.4:p.Asn696Ser)<br>NOTCH1 (chr9:g.136505767G>A, ENSP00000498587.1:p.Pro1377Ser)<br>NOTCH2 (chr1:g.119915499A>T, ENSP00000256646.2:p.Leu2408His)<br>RB1 (chr13:g.48381295G>A, ENSP00000267163.4:p.Trp516Ter)<br>RET (chr10:g.43102379C>A, ENSP00000491505.1:p.Leu107Ile)<br>SETD7 (chr4:g.139496517G>C, ENSP00000427300.1:p.Gln309Glu)<br>SOX11 (chr2:g.5693411C>A, ENSP00000322568.3:p.Asp230Glu)<br>STAG2 (chrX:g.124094127dup, ENSP00000218089.9:p.Thr1230AsnfsTer21) | methylated | GBM, IDHwt, RTK I | Mesenchymal-like | NA |
| T797 | LIH0797 | 55 | Female | Left occipital | primary | no prior treatment | GBM Grade 4 | GBM, IDHwt, RTK I | complex genome ++[CDK4, MDM2], +[Chr 7, Chr 20], -[Chr 10, 13q, 15q, 22q] | ALK (chr2:g.29320870A>G,ENSP00000373700.3:p.Val476Ala)<br>BCL2 (chr18:g.63318164G>A,ENSP00000329623.3:p.Pro168Leu)<br>BRCA1 (chr17:g.43070958C>T,ENSP00000312236.5:p.Met510Ile)<br>BRCA1 (chr17:g.43092412C>T,ENSP00000326002.7:p.Ser1040Asn)<br>COL6A3 (chr2:g.237336138T>C,ENSP00000295550.4:p.Met2988Val)<br>HMCN1 (chr1:g.186055511G>A,ENSP00000271588.4:p.Met2327Ile)<br>HMCN1 (chr1:g.186122971C>T,ENSP00000271588.4:p.His4084Tyr)<br>HMCN1 (chr1:g.186144595G>A,ENSP00000271588.4:p.Ala4720Thr)<br>KDM1B (chr6:g.18171430A>C,ENSP00000297792.5:p.Gln162Pro)<br>KMT2C (chr7:g.152180042G>T,ENSP00000262189.6:p.Pro2412Thr)<br>KMT2C (chr7:g.152252650T>C,ENSP00000262189.6:p.Ile455Met)<br>MRE11 (chr11:g.94470486G>C,ENSP00000325863.4:p.Ser334Arg)<br>PIK3R1 (chr5:g.68293760_68293765del,ENSP00000323512.8:p.Glu151_Tyr152del)<br>SETD7 (chr4:g.139496517G>C,ENSP00000427300.1:p.Gln309Glu)<br>SMO (chr7:g.129189225A>G,ENSP00000249373.3:p.Asp25Gly)<br>SMO (chr7:g.129205670G>A,ENSP00000249373.3:p.Val270Ile) | methylated (wide range) | GBM, IDHwt, RTK I | Classic-like | NA |
| T831 | LIH0831 (n | 51 | Female | Bifrontal | primary (multifo | no prior treatment | GBM Grade 4 | GBM, IDHwt, RTK II | ++EGFR, +[Chr 7], -[3p12.1, 6p24.2, Chr 10, 13q, 16p12.2], --CDKN2A/B | DNMT3B (chr20:g.32798560C>T, ENSP00000201963.3:p.Arg523Cys)<br>FOXO3 (chr6:g.108561627C>T, ENSP00000339527.6:p.Ala140Val)<br>GABRA6 (chr5:g.161687516T>C, ENSP00000429943.1:p.Leu13Pro)<br>GABRA6 (chr5:g.161687516T>C, ENSP00000429943.1:p.Leu13Pro)<br>GLI1 (chr12:g.57469650C>T, ENSP00000228682.2:p.Arg510Trp)<br>KMT2C (chr7:g.152180042G>T, ENSP00000262189.6:p.Pro2412Thr)<br>KMT2D (chr12:g.49051644_49051645delinsAG, ENSP00000301067.7:p.Thr680Leu)<br>KMT2D (chr12:g.49051802_49051963del, ENSP00000301067.7:p.Glu578_Glu631del)<br>MSH4 (chr1:g.75880138A>G, ENSP00000263187.3:p.Tyr589Cys)<br>MUC17 (chr7:g.101043780A>C, ENSP00000302716.4:p.Asn4122His)<br>MYB (chr6:g.135202648G>A, ENSP00000326328.8:p.Val645Met)<br>NIPBL (chr5:g.36985674C>T, ENSP00000282516.8:p.Arg832Ter)<br>PCDH8 (chr13:g.52846209A>G, ENSP00000341350.4:p.Val743Ala)<br>PCDH8 (chr13:g.52847338T>C, ENSP00000341350.4:p.Thr367Ala)<br>PHC2 (chr1:g.33355022G>T, ENSP00000257118.5:p.Pro403His)<br>PIK3CA (chr3:g.179199048C>G, ENSP00000263967.3:p.Gln75Glu)<br>PRKAR1A (chr17:g.68551108G>A, ENSP00000467311.2:p.Ser333Asn)<br>SETD7 (chr4:g.139496517G>C, ENSP00000427300.1:p.Gln309Glu)<br>SMARCD1 (chr12:g.50086319G>T, ENSP00000370924.4:p.Gln112His) | methylated | GBM, IDHwt, RTK I/II | Classic-like | NA |
| T832 | LIH0831 (n | 51 | Female | Bifrontal | primary (multifo | no prior treatment | GBM Grade 4 | GBM, IDHwt, RTK II | ++EGFR, +[Chr 7], -[Chr 10, 13q], --CDKN2A/B | DNMT3B (chr20:g.32798560C>T,ENSP00000201963.3:p.Arg523Cys)<br>EGFR (chr7:g.55143387G>A,ENSP00000275493.2:p.Arg108Lys)<br>FOXO3 (chr6:g.108561627C>T,ENSP00000339527.6:p.Ala140Val)<br>GABRA6 (chr5:g.161687516T>C,ENSP00000429943.1:p.Leu13Pro)<br>GABRA6 (chr5:g.161687527C>T,ENSP00000430212.1:p.Thr11Ile)<br>H2AC15 (chr6:g.27837976_27837977del,ENSP00000482431.2:p.Ser123ProfsTer?)<br>KMT2C (chr7:g.152180042G>T,ENSP00000262189.6:p.Pro2412Thr)<br>MSH4 (chr1:g.75880138A>G,ENSP00000263187.3:p.Tyr589Cys)<br>MUC17 (chr7:g.101043780A>C,ENSP00000302716.4:p.Asn4122His)<br>MYB (chr6:g.135202648G>A,ENSP00000326328.8:p.Val645Met)<br>NIPBL (chr5:g.36985674C>T,ENSP00000282516.8:p.Arg832Ter)<br>PCDH8 (chr13:g.52846209A>G,ENSP00000341350.4:p.Val743Ala)<br>PCDH8 (chr13:g.52847338T>C,ENSP00000341350.4:p.Thr367Ala)<br>PHC2 (chr1:g.33355022G>T,ENSP00000257118.5:p.Pro403His)<br>PIK3CA (chr3:g.179199048C>G,ENSP00000263967.3:p.Gln75Glu)<br>PRKAR1A (chr17:g.68551108G>A,ENSP00000467311.2:p.Ser333Asn)<br>SETD7 (chr4:g.139496517G>C,ENSP00000427300.1:p.Gln309Glu)<br>SMARCD1 (chr12:g.50086319G>T,ENSP00000370924.4:p.Gln112His) | methylated | GBM, IDHwt, RTK II | Classic-like | Classical |
| T841 | LIH0841 | 38 | Male | Right temporal | primary | no prior treatment | GBM Grade 4 | GBM, IDHwt, other (no match: RTK I /Pediatric-type diffuse high-grade glioma, H3wt, IDHwt, MYCN subtype) | complex genome ++[MYCN, CDK4], +Chr 7, -[2p24.2, 6q12, Chr 10, 13q31.2, 14q12, 15q15.3] | CDKN2A (chr9:g.21970917C>T, ENSP00000307101.5:p.Ala148Thr)<br>GABRA6 (chr5:g.161687516T>C, ENSP00000429943.1:p.Leu13Pro)<br>GNAS (chr20:g.58853943T>G, ENSP00000302237.4:p.Leu228Trp)<br>HMCN1 (chr1:g.186087223G>A, ENSP00000271588.4:p.Arg3018Gln)<br>KDM5A (chr12:g.387273A>T, ENSP00000444251.1:p.Phe56Tyr)<br>MET (chr7:g.116695757A>T, ENSP00000413857.1:p.Met1?)<br>MPL (chr1:g.43338669G>A, ENSP00000361548.3:p.Val114Met)<br>MUC17 (chr7:g.101036598C>A, ENSP00000302716.4:p.Arg1728Ser)<br>PIK3C2G (chr12:g.18290875C>A, ENSP00000404845.1:p.Ala261Glu)<br>PIK3CA (chr3:g.179199156_179199161del, ENSP00000263967.3:p.Lys111_Ile112del)<br>PLCG1 (chr20:g.41172488C>T, ENSP00000468958.1:p.Thr50Met)<br>SETD7 (chr4:g.139496517G>C, ENSP00000427300.1:p.Gln309Glu) | unmethylated | GBM IDHwt, other (no match: Primit4e Neuronal component / Paediatric-type diffuse high-grade glioma, MYCN subtype) | LGm6-GBM | Proneural |
| T861 | LIH0841 | 38 | Male | Right temporal | recurrence 1 | radiotherapy + TMZ | GBM Grade 4 | NA | NA | NA | NA | NA | NA | NA |
| T899 | LIH0609 | 75 | Female | Right central | recurrence 1 | no prior treatment | GBM Grade 4 | GBM, IDHwt, other (no match: RTK I midline /Diffuse paediatric-type high grade glioma, RTK1 subtype) | complex genome ++[PDGFRA, CDK4], -[Chr10, 14q] | ABL1 (chr9:g.130884406G>A, ENSP00000323315.5:p.Gly706Ser)<br>BRCA1 (chr17:g.43092412C>T, ENSP00000326002.7:p.Ser1040Asn)<br>CDK4 (chr12:g.57749225G>A, ENSP00000257904.5:p.Ser259Leu)<br>GABRA6 (chr5:g.161687516T>C, ENSP00000429943.1:p.Leu13Pro)<br>GLI1 (chr12:g.57468181C>T, ENSP00000228682.2:p.Pro422Leu)<br>HMCN1 (chr1:g.186082892G>A, ENSP00000271588.4:p.Gly2939Ser)<br>HMCN1 (chr1:g.186144605G>A, ENSP00000271588.4:p.Arg4723Lys)<br>HMCN1 (chr1:g.186151341G>A, ENSP00000271588.4:p.Arg4917His)<br>KMT2C (chr7:g.152180042G>T, ENSP00000262189.6:p.Pro2412Thr)<br>LRP1B (chr2:g.140541066C>T, ENSP00000374135.3:p.Gly2474Ser)<br>MSH6 (chr2:g.47800616T>C, ENSP00000234420.5:p.Val878Ala)<br>MUC17 (chr7:g.101040843C>A, ENSP00000302716.4:p.His3143Asn)<br>NCOR1 (chr17:g.16073554C>T, ENSP00000268712.2:p.Arg1229Gln)<br>PIK3R1 (chr5:g.68296465A>C, ENSP00000513328.1:p.Gln235His)<br>PTCH1 (chr9:g.95468828G>A, ENSP00000332353.6:p.Pro725Ser)<br>SETD7 (chr4:g.139496517G>C, ENSP00000427300.1:p.Gln309Glu)<br>TP53 (chr17:g.7674188C>A, ENSP00000269305.4:p.Asp259Tyr)<br>ZFTA (chr11:g.63765915C>T, ENSP00000483097.1:p.Gly17Arg)<br>ZNF814 (chr19:g.57873840G>T, ENSP00000410545.1:p.Ala517Glu) | methylated (wide range) | GBM, IDHwt other (no match: Pediatric-type diffuse high grade-glioma, H3 wildtype, RTK I) | LGm6-GBM | NA |

| | | | | | | | | | | | | | | |
|---|---|---|---|---|---|---|---|---|---|---|---|---|---|---|
| T905 | LIH0609 | 76 | Female | Right central | recurrence2 | no prior treatment | GBM Grade 4 | NA | complex genome ++[PDGFRA, EGFR, CDK4], -[Chr10, 13q] | ABL1 (chr9:g.130884406G>A,ENSP00000323315.5:p.Gly706Ser)<br>BRCA1 (chr17:g.43092412C>T,ENSP00000326002.7:p.Ser1040Asn)<br>DNMT3B (chr20:g.32798560C>T,ENSP00000201963.3:p.Arg523Cys)<br>EGFR (chr7:g.55143387G>A,ENSP00000275493.2:p.Arg108Lys)<br>FOXO1 (chr13:g.40560208T>G,ENSP00000368880.4:p.Gln428Pro)<br>FOXO3 (chr6:g.108561627C>T,ENSP00000339527.6:p.Ala140Val)<br>GABRA6 (chr5:g.161687516T>C,ENSP00000429943.1:p.Leu13Pro)<br>GABRA6 (chr5:g.161687527C>T,ENSP00000430212.1:p.Thr11Ile)<br>GLI1 (chr12:g.57468181C>T,ENSP00000228682.2:p.Pro422Leu)<br>H2AC15 (chr6:g.27837976_27837977del,ENSP00000482431.2:p.Ser123ProfsTer?)<br>HMCN1 (chr1:g.186082892G>A,ENSP00000271588.4:p.Gly2939Ser)<br>HMCN1 (chr1:g.186144605G>A,ENSP00000271588.4:p.Arg4723Lys)<br>HMCN1 (chr1:g.186151341G>A,ENSP00000271588.4:p.Arg4917His)<br>KMT2C (chr7:g.152180042G>T,ENSP00000262189.6:p.Pro2412Thr)<br>LRP1B (chr2:g.140541066C>T,ENSP00000374135.3:p.Gly2474Ser)<br>MSH6 (chr2:g.47800616T>C,ENSP00000234420.5:p.Val878Ala)<br>MUC17 (chr7:g.101043780A>C,ENSP00000302716.4:p.Asn4122His)<br>NCOR1 (chr17:g.16073554C>T,ENSP00000268712.2:p.Arg1229Gln)<br>NIPBL (chr5:g.36985674C>T,ENSP00000282516.8:p.Arg832Ter)<br>PCDH8 (chr13:g.52846209A>G,ENSP00000341350.4:p.Val743Ala)<br>PCDH8 (chr13:g.52847338T>C,ENSP00000341350.4:p.Thr367Ala)<br>PHC2 (chr1:g.33355022G>T,ENSP00000257118.5:p.Pro403His)<br>PIK3CA (chr3:g.179199048C>G,ENSP00000263967.3:p.Gln75Glu)<br>PIK3R1 (chr5:g.68296465A>C,ENSP00000513328.1:p.Gln235His)<br>PRKAR1A (chr17:g.68551108G>A,ENSP00000467311.2:p.Ser333Asn)<br>PTCH1 (chr9:g.95468828G>A,ENSP00000332353.6:p.Pro725Ser)<br>SETD7 (chr4:g.139496517G>C,ENSP00000427300.1:p.Gln309Glu)<br>SMARCD1 (chr12:g.50086319G>T,ENSP00000370924.4:p.Gln112His)<br>TP53 (chr17:g.7674188C>A,ENSP00000269305.4:p.Asp259Tyr)<br>ZFTA (chr11:g.63765915C>T,ENSP00000483097.1:p.Gly17Arg)<br>ZNF587 (chr19:g.57858894G>A,ENSP00000345479.4:p.Arg161His) | methylated | GBM, IDHwt, RTK I | Mesenchymal-like | NA |
| T1020 | LIH0973 | 49 | Male | Right parietal lobe | recurrence 1 | radiotherapy + TMZ | GBM Grade 4 | GBM, IDHwt, RTK II | | | NA | NA | | NA |
| T1053 | LIH1053 | 62 | Male | Right intraaxial | primary | no prior treatment | GBM Grade 4 | GBM, IDHwt, RTK II | NA | NA | NA | NA | NA | NA |
| T1067 | LIH1067 | 84 | Female | Right parieto-opercular | primary | no prior treatment | GBM Grade 4 | GBM, IDHwt, mesenchymal | | | NA | NA | NA | NA |
| T1070 | LIH0784 | 60 | Male | Left frontal | recurrence 1 | radiotherapy + TMZ | GBM Grade 4 | GBM, IDHwt, RTK I | NA | NA | NA | NA | NA | NA |

**Table S2. List of inhibitors applied in the study.**
202 drugs are numbered in the first column. Each line represents a specific target and associated gene family per each drug.

| Number | Inhibitor | Target | Family | Source |
|---|---|---|---|---|
| 1 | MI-463 | Menin-MLL | Histone Methyltransferase complex | Custom |
| 2 | MI-503 | Menin-MLL | Histone Methyltransferase complex | Custom |
| 3 | EPZ020411 | PRMT6 | Histone Methyltransferase | Custom |
| 4 | OICR-9429 | H3 | Histone | Custom |
| 4 | OICR-9429 | MLL | Histone Methyltransferase complex | Custom |
| 4 | OICR-9429 | WDR | Histone Methyltransferase complex | Custom |
| 5 | CPI-0610 | BET bromodomain | Epigenetic Reader Domain | Custom |
| 6 | A-196 | SUV420H2 | Histone Methyltransferase | Custom |
| 6 | A-196 | SUV420H1 | Histone Methyltransferase | Custom |
| 7 | UNC0638 | GLP | Histone Methyltransferase | Custom |
| 7 | UNC0638 | G9A | Histone Methyltransferase | Custom |
| 8 | MS049 | PRMT6 | Histone Methyltransferase | Custom |
| 8 | MS049 | PRMT4 | Histone Methyltransferase | Custom |
| 9 | CPI-637 | CBP/EP300 | Epigenetic Reader Domain | Custom |
| 10 | GSK6853 | BRPF1 | Epigenetic Reader Domain | Custom |
| 11 | CPI-455 HCL | KDM5 (JARID1A) | Histone Demethylase | Custom |
| 12 | SGC2085 | CARM1 (PRMT4) | Histone Methyltransferase | Custom |
| 13 | AZD5153 | BRD4 | Epigenetic Reader Domain | Custom |
| 14 | CPI-1205 | EZH2 (KMT6) | Histone Methyltransferase | Custom |
| 15 | UNC3866 | CBX7 | Histone Methyltransferase complex | Custom |
| 15 | UNC3866 | CBX4 | Histone Methyltransferase complex | Custom |
| 16 | Mivebresib (ABBV-075) | BRDT | Epigenetic Reader Domain | Custom |
| 16 | Mivebresib (ABBV-075) | BRD4 | Epigenetic Reader Domain | Custom |
| 16 | Mivebresib (ABBV-075) | BRD2 | Epigenetic Reader Domain | Custom |
| 17 | PF-06726304 | H3K27me3 | Histone | Custom |
| 17 | PF-06726304 | EZH2 (KMT6) | Histone Methyltransferase | Custom |
| 18 | EED226 | PRC2 | Histone Methyltransferase complex | Custom |
| 19 | SF2523 | BRD4 | Epigenetic Reader Domain | Custom |
| 19 | SF2523 | DNA-PK | Kinase | Custom |
| 19 | SF2523 | mTOR | Kinase | Custom |
| 19 | SF2523 | PI3Kγ | Kinase | Custom |
| 19 | SF2523 | PI3Kα | Kinase | Custom |
| 20 | CP2 | KDM4 (JMJD2) | Histone Demethylase | Custom |
| 21 | JNJ-64619178 | PRMT5 | Histone Methyltransferase | Custom |
| 22 | AZD2461 | PARP | DNA damage Signaling | SelleckChem, L1900 |
| 23 | XL019 | JAK | Kinase | SelleckChem, L1900 |
| 24 | CX-6258 HCl | Pim-1 | Pim-Kinase | SelleckChem, L1900 |
| 24 | CX-6258 HCl | Pim-2 | Pim-Kinase | SelleckChem, L1900 |
| 24 | CX-6258 HCl | Pim-3 | Pim-Kinase | SelleckChem, L1900 |
| 24 | CX-6258 HCl | pan-Pim | Pim-Kinase | SelleckChem, L1900 |
| 25 | Pinometostat (EPZ5676) | H3K79 methylation | Histone Methyltransferase | SelleckChem, L1900 |
| 25 | Pinometostat (EPZ5676) | DOT1L | Histone Methyltransferase | SelleckChem, L1900 |
| 26 | GSK J4 HCl | UTX | Histone Demethylase | SelleckChem, L1900 |
| 26 | GSK J4 HCl | JMJD3 | Histone Demethylase | SelleckChem, L1900 |
| 27 | SGC 0946 | DOT1L | Histone Methyltransferase | SelleckChem, L1900 |
| 28 | UNC1215 | L3MBTL3 | Epigenetic Reader Domain | SelleckChem, L1900 |
| 29 | AZD1208 | Pim-1 | Pim-Kinase | SelleckChem, L1900 |
| 29 | AZD1208 | Pim-2 | Pim-Kinase | SelleckChem, L1900 |
| 29 | AZD1208 | Pim-3 | Pim-Kinase | SelleckChem, L1900 |
| 29 | AZD1208 | pan-Pim | Pim-Kinase | SelleckChem, L1900 |
| 30 | (+)-JQ1 | BRD4 | Epigenetic Reader Domain | SelleckChem, L1900 |
| 30 | (+)-JQ1 | BRD3 | Epigenetic Reader Domain | SelleckChem, L1900 |
| 30 | (+)-JQ1 | BRD2 | Epigenetic Reader Domain | SelleckChem, L1900 |
| 30 | (+)-JQ1 | BRD6 | Epigenetic Reader Domain | SelleckChem, L1900 |
| 31 | Zebularine | pan DNMTs | DNA Methyltransferase | SelleckChem, L1900 |
| 32 | 3-deazaneplanocin A (DZNeP) | EZH2 | Histone Methyltransferase | SelleckChem, L1900 |
| 33 | C646 | p300 | Histone Acetyltransferase | SelleckChem, L1900 |
| 34 | I-BET-762 | pan-BET | Epigenetic Reader Domain | SelleckChem, L1900 |
| 35 | RGFP966 | HDAC3 | Histone Deacetylase | SelleckChem, L1900 |
| 36 | GSK2801 | BAZ2B | Epigenetic Reader Domain | SelleckChem, L1900 |
| 36 | GSK2801 | BAZ2A | Epigenetic Reader Domain | SelleckChem, L1900 |
| 37 | Bromosporine | BRD2 | Epigenetic Reader Domain | SelleckChem, L1900 |
| 37 | Bromosporine | BRD4 | Epigenetic Reader Domain | SelleckChem, L1900 |
| 37 | Bromosporine | BRD9 | Epigenetic Reader Domain | SelleckChem, L1900 |
| 37 | Bromosporine | CECR2 | Epigenetic Reader Domain | SelleckChem, L1900 |
| 38 | IOX1 | 2OG oxygenases | Histone Demethylase | SelleckChem, L1900 |
| 39 | OG-L002 | LSD1 | Histone Demethylase | SelleckChem, L1900 |
| 40 | NVP-TNKS656 | PARP | DNA damage Signaling | SelleckChem, L1900 |
| 41 | SGC-CBP30 | CREBBP/EP300 | Epigenetic Reader Domain | SelleckChem, L1900 |
| 42 | MM-102 | MLL1 | Histone Methyltransferase | SelleckChem, L1900 |
| 43 | SGI-1027 | DNMT1 | DNA Methyltransferase | SelleckChem, L1900 |
| 43 | SGI-1027 | DNMT3A | DNA Methyltransferase | SelleckChem, L1900 |
| 43 | SGI-1027 | DNMT3B | DNA Methyltransferase | SelleckChem, L1900 |
| 44 | JIB-04 | JARID1A | Histone Demethylase | SelleckChem, L1900 |
| 44 | JIB-04 | JMJD2E | Histone Demethylase | SelleckChem, L1900 |
| 44 | JIB-04 | JMJD3 (KDM6B) | Histone Demethylase | SelleckChem, L1900 |
| 44 | JIB-04 | JMJD2B | Histone Demethylase | SelleckChem, L1900 |
| 44 | JIB-04 | JMJD2C | Histone Demethylase | SelleckChem, L1900 |
| 44 | JIB-04 | JMJD2D | Histone Demethylase | SelleckChem, L1900 |
| 45 | RG2833 (RGFP109) | HDAC3 | Histone Deacetylase | SelleckChem, L1900 |
| 45 | RG2833 (RGFP109) | HDAC1 | Histone Deacetylase | SelleckChem, L1900 |
| 46 | PFI-2 HCl | SETD7 | Histone Methyltransferase | SelleckChem, L1900 |
| 47 | RVX-208 | BD2 | Epigenetic Reader Domain | SelleckChem, L1900 |
| 48 | ML324 | JMJD2 (KDM4) | Histone Demethylase | SelleckChem, L1900 |
| 49 | PJ34 HCl | PARP | DNA damage Signaling | SelleckChem, L1900 |
| 50 | CPI-203 | BRD4 | Epigenetic Reader Domain | SelleckChem, L1900 |
| 51 | MS436 | BRD4 | Epigenetic Reader Domain | SelleckChem, L1900 |
| 52 | PFI-3 | SMARCA2 | Epigenetic Reader Domain | SelleckChem, L1900 |
| 52 | PFI-3 | SMARCA4 | Epigenetic Reader Domain | SelleckChem, L1900 |
| 52 | PFI-3 | PB1 | Epigenetic Reader Domain | SelleckChem, L1900 |
| 53 | TMP269 | HDAC | Histone Deacetylase | SelleckChem, L1900 |

| | | | | |
|---|---|---|---|---|
| 54 | EPZ004777 | DOT1L | Histone Methyltransferase | SelleckChem, L1900 |
| 55 | OTX015 | BRD4 | Epigenetic Reader Domain | SelleckChem, L1900 |
| 55 | OTX015 | BRD3 | Epigenetic Reader Domain | SelleckChem, L1900 |
| 55 | OTX015 | BRD2 | Epigenetic Reader Domain | SelleckChem, L1900 |
| 56 | UNC669 | MBT1 | Epigenetic Reader Domain | SelleckChem, L1900 |
| 56 | UNC669 | MBT3 | Epigenetic Reader Domain | SelleckChem, L1900 |
| 56 | UNC669 | MBT4 | Epigenetic Reader Domain | SelleckChem, L1900 |
| 57 | ME0328 | PARP | DNA damage Signaling | SelleckChem, L1900 |
| 58 | Nexturastat A | HDAC6 | Histone Deacetylase | SelleckChem, L1900 |
| 59 | MG149 | Tip60 | Histone Acetyltransferase | SelleckChem, L1900 |
| 59 | MG149 | MOF | Histone Acetyltransferase | SelleckChem, L1900 |
| 60 | Decernotinib (VX-509) | JAK | Kinase | SelleckChem, L1900 |
| 61 | 4SC-202 | HDAC | Histone Deacetylase | SelleckChem, L1900 |
| 62 | UNC0379 | SETD8 | Histone Methyltransferase | SelleckChem, L1900 |
| 63 | A-366 | G9a | Histone Methyltransferase | SelleckChem, L1900 |
| 63 | A-366 | GLP | Histone Methyltransferase | SelleckChem, L1900 |
| 64 | GSK-LSD1 2HCl | LSD1 | Histone Demethylase | SelleckChem, L1900 |
| 65 | GSK J1 | JMJD3 (KDM6B) | Histone Demethylase | SelleckChem, L1900 |
| 65 | GSK J1 | UTX (KDM6A) | Histone Demethylase | SelleckChem, L1900 |
| 66 | Anacardic Acid | p300/CBP | Histone Acetyltransferase | SelleckChem, L1900 |
| 67 | BRD4770 | G9a | Histone Methyltransferase | SelleckChem, L1900 |
| 67 | BRD4770 | GLP | Histone Methyltransferase | SelleckChem, L1900 |
| 68 | Filgotinib (GLPG0634) | JAK | Kinase | SelleckChem, L1900 |
| 69 | UNC0631 | G9a | Histone Methyltransferase | SelleckChem, L1900 |
| 69 | UNC0631 | GLP | Histone Methyltransferase | SelleckChem, L1900 |
| 70 | EI1 | EZH2 | Histone Methyltransferase | SelleckChem, L1900 |
| 71 | CPI-169 | EZH2 | Histone Methyltransferase | SelleckChem, L1900 |
| 72 | MI-2 | Menin-MLL Inhibitor | Histone Methyltransferase | SelleckChem, L1900 |
| 73 | MI-3 | Menin-MLL Inhibitor | Histone Methyltransferase | SelleckChem, L1900 |
| 74 | GSK1324726A (I-BET726) | BRD2 | Epigenetic Reader Domain | SelleckChem, L1900 |
| 74 | GSK1324726A (I-BET726) | BRD3 | Epigenetic Reader Domain | SelleckChem, L1900 |
| 74 | GSK1324726A (I-BET726) | BRD4 | Epigenetic Reader Domain | SelleckChem, L1900 |
| 75 | Niraparib (MK-4827) tosylate | PARP | DNA damage Signaling | SelleckChem, L1900 |
| 76 | Remodelin | NAT10 | Histone Acetyltransferase | SelleckChem, L1900 |
| 77 | CPI-360 | EZH1 | Histone Methyltransferase | SelleckChem, L1900 |
| 78 | SP2509 | LSD1 | Histone Demethylase | SelleckChem, L1900 |
| 79 | OF-1 | BRPF1B | Epigenetic Reader Domain | SelleckChem, L1900 |
| 79 | OF-1 | BRPF2 | Epigenetic Reader Domain | SelleckChem, L1900 |
| 80 | EPZ015666(GSK3235025) | PRMT5 | Histone Methyltransferase | SelleckChem, L1900 |
| 81 | AZ6102 | PPAR | Transcription factor activator | SelleckChem, L1900 |
| 82 | ORY-1001 (RG-6016) 2HCl | LSD1 (KDM1A) | Histone Demethylase | SelleckChem, L1900 |
| 83 | GSK2879552 2HCl | LSD1 (KDM1A) | Histone Demethylase | SelleckChem, L1900 |
| 84 | GSK503 | EZH2 | Histone Methyltransferase | SelleckChem, L1900 |
| 85 | EPZ011989 | EZH2 | Histone Methyltransferase | SelleckChem, L1900 |
| 86 | SGC707 | PRMT3 | Histone Methyltransferase | SelleckChem, L1900 |
| 87 | I-BRD9 | BRD4 | Epigenetic Reader Domain | SelleckChem, L1900 |
| 87 | I-BRD9 | BRD9 | Epigenetic Reader Domain | SelleckChem, L1900 |
| 88 | Ricolinostat (ACY-1215) | HDAC | Histone Deacetylase | SelleckChem, L1900 |
| 89 | ZM 39923 HCl | JAK | Kinase | SelleckChem, L1900 |
| 90 | SMI-4a | Pim-2 | Pim-Kinase | SelleckChem, L1900 |
| 91 | BIX 01294 | G9a | Histone Methyltransferase | SelleckChem, L1900 |
| 92 | UPF 1069 | PARP | DNA damage Signaling | SelleckChem, L1900 |
| 93 | Scriptaid | HDAC | Histone Deacetylase | SelleckChem, L1900 |
| 94 | Tubastatin A | HDAC | Histone Deacetylase | SelleckChem, L1900 |
| 95 | Lomeguatrib | O6-alkylguanine-DNA-alkyltransf | DNA Methyltransferase | SelleckChem, L1900 |
| 96 | Pacritinib (SB1518) | JAK2 | Kinase | SelleckChem, L1900 |
| 96 | Pacritinib (SB1518) | FLT3 | Kinase | SelleckChem, L1900 |
| 97 | Mirin | ATM | DNA damage Signaling | SelleckChem, L1900 |
| 97 | Mirin | ATR | DNA damage Signaling | SelleckChem, L1900 |
| 98 | GSK591 | PRMT5 | Histone Methyltransferase | SelleckChem, L1900 |
| 99 | MS023 | PRMT1 | Histone Methyltransferase | SelleckChem, L1900 |
| 99 | MS023 | PRMT3 | Histone Methyltransferase | SelleckChem, L1900 |
| 99 | MS023 | PRMT4 | Histone Methyltransferase | SelleckChem, L1900 |
| 99 | MS023 | PRMT8 | Histone Methyltransferase | SelleckChem, L1900 |
| 100 | Mitomycin C | DNA/RNA | DNA/RNA Synthesis | SelleckChem, L1900 |
| 101 | BI-7273 | BRD9 | Epigenetic Reader Domain | SelleckChem, L1900 |
| 101 | BI-7273 | BRD7 | Epigenetic Reader Domain | SelleckChem, L1900 |
| 102 | PF-CBP1 HCl | p300 | Epigenetic Reader Domain | SelleckChem, L1900 |
| 102 | PF-CBP1 HCl | CREBBP | Epigenetic Reader Domain | SelleckChem, L1900 |
| 103 | Oclacitinib | JAK | Kinase | SelleckChem, L1900 |
| 104 | HLCL-61 HCL | PRMT5 | Histone Methyltransferase | SelleckChem, L1900 |
| 105 | ITSA-1 (ITSA1) | HDAC | Histone Deacetylase | SelleckChem, L1900 |
| 106 | Veliparib (ABT-888) | PARP | DNA damage Signaling | SelleckChem, L1900 |
| 107 | Roxadustat (FG-4592) | HIF | Transcription factor | SelleckChem, L1900 |
| 108 | Panobinostat (LBH589) | HDAC | Histone Deacetylase | SelleckChem, L1900 |
| 109 | Trichostatin A (TSA) | HDAC | Histone Deacetylase | SelleckChem, L1900 |
| 110 | Vorinostat (SAHA, MK0683) | HDAC | Histone Deacetylase | SelleckChem, L1900 |
| 111 | Tozasertib (VX-680, MK-0457) | Aurora Kinase | Aurora Kinase | SelleckChem, L1900 |
| 112 | Entinostat (MS-275) | HDAC 3 | Histone Deacetylase | SelleckChem, L1900 |
| 112 | Entinostat (MS-275) | HDAC 1 | Histone Deacetylase | SelleckChem, L1900 |
| 113 | Olaparib (AZD2281, Ku-00594 | PARP | DNA damage Signaling | SelleckChem, L1900 |
| 114 | Belinostat (PXD101) | HDAC | Histone Deacetylase | SelleckChem, L1900 |
| 115 | Iniparib (BSI-201) | PARP | DNA damage Signaling | SelleckChem, L1900 |
| 116 | Abexinostat (PCI-24781) | HDAC1 | Histone Deacetylase | SelleckChem, L1900 |
| 117 | Dacinostat (LAQ824) | HDAC | Histone Deacetylase | SelleckChem, L1900 |
| 118 | Quisinostat (JNJ-26481585) 2H | HDAC | Histone Deacetylase | SelleckChem, L1900 |
| 119 | Rucaparib (AG-014699,PF-013 | PARP | DNA damage Signaling | SelleckChem, L1900 |
| 120 | MLN8054 | Aurora Kinase | Aurora Kinase | SelleckChem, L1900 |
| 121 | ZM 447439 | Aurora Kinase | Aurora Kinase | SelleckChem, L1900 |
| 122 | Danusertib (PHA-739358) | c-RET | Aurora Kinase | SelleckChem, L1900 |
| 122 | Danusertib (PHA-739358) | FGFR | Aurora Kinase | SelleckChem, L1900 |
| 122 | Danusertib (PHA-739358) | Bcr-Abl | Kinase | SelleckChem, L1900 |
| 123 | Mocetinostat (MGCD0103) | HDAC 1 | Histone Deacetylase | SelleckChem, L1900 |
| 124 | SRT1720 HCl | SIRT1 | Histone Deacetylase | SelleckChem, L1900 |
| 125 | INO-1001 (3-Aminobenzamide | PARP | DNA damage Signaling | SelleckChem, L1900 |

| | | | | |
|---|---|---|---|---|
| 126 | Alisertib (MLN8237) | Aurora A | Aurora Kinase | SelleckChem, L1900 |
| 127 | AT9283 | Bcr-Abl | Kinase | SelleckChem, L1900 |
| 127 | AT9283 | JAK | Kinase | SelleckChem, L1900 |
| 128 | AG-490 (Tyrphostin B42) | JAK | Kinase | SelleckChem, L1900 |
| 128 | AG-490 (Tyrphostin B42) | EGFR | Receptor | SelleckChem, L1900 |
| 129 | Barasertib (AZD1152-HQPA) | Aurora Kinase | Aurora Kinase | SelleckChem, L1900 |
| 130 | SNS-314 Mesylate | Aurora A | Aurora Kinase | SelleckChem, L1900 |
| 130 | SNS-314 Mesylate | Aurora B | Aurora Kinase | SelleckChem, L1900 |
| 130 | SNS-314 Mesylate | Aurora C | Aurora Kinase | SelleckChem, L1900 |
| 131 | Valproic acid sodium salt (Sodi | HDAC | Histone Deacetylase | SelleckChem, L1900 |
| 131 | Valproic acid sodium salt (Sodi | GABA Receptor | Receptor | SelleckChem, L1900 |
| 132 | CYC116 | Aurora Kinase | Aurora Kinase | SelleckChem, L1900 |
| 132 | CYC116 | VEGFR | Receptor | SelleckChem, L1900 |
| 133 | ENMD-2076 | FLT3 | Kinase | SelleckChem, L1900 |
| 133 | ENMD-2076 | VEGFR | Receptor | SelleckChem, L1900 |
| 134 | CUDC-101 | HDAC | Histone Deacetylase | SelleckChem, L1900 |
| 134 | CUDC-101 | HER2 | Receptor | SelleckChem, L1900 |
| 134 | CUDC-101 | EGFR | Receptor | SelleckChem, L1900 |
| 135 | Decitabine | pan DNA Methyltransferase | DNA Methyltransferase | SelleckChem, L1900 |
| 136 | PFI-1 (PF-6405761) | BRD4 | Epigenetic Reader Domain | SelleckChem, L1900 |
| 136 | PFI-1 (PF-6405761) | BRD2 | Epigenetic Reader Domain | SelleckChem, L1900 |
| 137 | 2-Methoxyestradiol (2-MeOE2) | HIF | Transcription factor | SelleckChem, L1900 |
| 138 | JNJ-7706621 | Aurora Kinase | Aurora Kinase | SelleckChem, L1900 |
| 138 | JNJ-7706621 | CDK1 | Kinase | SelleckChem, L1900 |
| 138 | JNJ-7706621 | CDK2 | Kinase | SelleckChem, L1900 |
| 139 | Ellagic acid | DNA/RNA | Topoisomerase | SelleckChem, L1900 |
| 140 | Ruxolitinib (INCB018424) | JAK | Kinase | SelleckChem, L1900 |
| 141 | Pirarubicin | Topoisomerase | Topoisomerase | SelleckChem, L1900 |
| 142 | Resveratrol | Sirtuin | Histone Deacetylase | SelleckChem, L1900 |
| 143 | Droxinostat | HDAC6 | Histone Deacetylase | SelleckChem, L1900 |
| 143 | Droxinostat | HDAC8 | Histone Deacetylase | SelleckChem, L1900 |
| 144 | Aurora A Inhibitor I | Aurora A | Aurora Kinase | SelleckChem, L1900 |
| 145 | PHA-680632 | pan-Aurora A | Aurora Kinase | SelleckChem, L1900 |
| 146 | Ofloxacin | Topoisomerase | Topoisomerase | SelleckChem, L1900 |
| 147 | MC1568 | HDAC | Histone Deacetylase | SelleckChem, L1900 |
| 148 | Norfloxacin | Topoisomerase | Topoisomerase | SelleckChem, L1900 |
| 149 | Pracinostat (SB939) | HDAC | Histone Deacetylase | SelleckChem, L1900 |
| 150 | Hesperadin | Aurora B | Aurora Kinase | SelleckChem, L1900 |
| 151 | Selisistat (EX 527) | Sirt1 | Histone Deacetylase | SelleckChem, L1900 |
| 151 | Selisistat (EX 527) | Sirtuin | Histone Deacetylase | SelleckChem, L1900 |
| 152 | Azacitidine | pan DNA Methyltransferase | DNA Methyltransferase | SelleckChem, L1900 |
| 153 | PCI-34051 | HDAC 8 | Histone Deacetylase | SelleckChem, L1900 |
| 153 | PCI-34051 | HDAC | Histone Deacetylase | SelleckChem, L1900 |
| 154 | ENMD-2076 L-(+)-Tartaric acid | Aurora Kinase | Aurora Kinase | SelleckChem, L1900 |
| 154 | ENMD-2076 L-(+)-Tartaric acid | FLT3 | Kinase | SelleckChem, L1900 |
| 154 | ENMD-2076 L-(+)-Tartaric acid | VEGFR | Receptor | SelleckChem, L1900 |
| 155 | KW-2449 | Aurora Kinase | Aurora Kinase | SelleckChem, L1900 |
| 155 | KW-2447 | Bcr-Abl | Kinase | SelleckChem, L1900 |
| 155 | KW-2448 | FLT3 | Kinase | SelleckChem, L1900 |
| 156 | AZD1480 | JAK | Kinase | SelleckChem, L1900 |
| 157 | Givinostat (ITF2357) | HDAC | Histone Deacetylase | SelleckChem, L1900 |
| 158 | AG-14361 | PARP | DNA damage Signaling | SelleckChem, L1900 |
| 159 | Gandotinib (LY2784544) | JAK 2 | Kinase | SelleckChem, L1900 |
| 160 | SGI-1776 free base | Pim 1 | Kinase | SelleckChem, L1900 |
| 160 | SGI-1776 free base | Pim | Kinase | SelleckChem, L1900 |
| 161 | AZ 960 | JAK 2 | Kinase | SelleckChem, L1900 |
| 161 | AZ 960 | JAK | Kinase | SelleckChem, L1900 |
| 162 | Momelotinib (CYT387) | JAK1 | Kinase | SelleckChem, L1900 |
| 162 | Momelotinib (CYT387) | JAK2 | Kinase | SelleckChem, L1900 |
| 163 | AR-42 | HDAC | Histone Deacetylase | SelleckChem, L1900 |
| 164 | Quercetin | Sirtuin | Histone Deacetylase | SelleckChem, L1900 |
| 164 | Quercetin | PI3K | Kinase | SelleckChem, L1900 |
| 164 | Quercetin | PKC | Kinase | SelleckChem, L1900 |
| 164 | Quercetin | Src | Kinase | SelleckChem, L1900 |
| 165 | Daphnetin | PKC | Kinase | SelleckChem, L1900 |
| 165 | Daphnetin | PKA | Kinase | SelleckChem, L1900 |
| 165 | Daphnetin | EGFR | Receptor | SelleckChem, L1900 |
| 166 | Tubastatin A HCl | HDAC6 | Histone Deacetylase | SelleckChem, L1900 |
| 166 | Tubastatin A HCl | HDAC | Histone Deacetylase | SelleckChem, L1900 |
| 167 | NVP-BSK805 2HCl | JAK | Kinase | SelleckChem, L1900 |
| 168 | TG101209 | FLT3 | Kinase | SelleckChem, L1900 |
| 168 | TG101209 | JAK | Kinase | SelleckChem, L1900 |
| 168 | TG101209 | c-RET | Kinase | SelleckChem, L1900 |
| 169 | Resminostat | HDAC1 | Histone Deacetylase | SelleckChem, L1900 |
| 169 | Resminostat | HDAC6 | Histone Deacetylase | SelleckChem, L1900 |
| 169 | Resminostat | HDAC3 | Histone Deacetylase | SelleckChem, L1900 |
| 169 | Resminostat | HDAC | Histone Deacetylase | SelleckChem, L1900 |
| 170 | TAK-901 | Aurora B | Aurora Kinase | SelleckChem, L1900 |
| 170 | TAK-901 | Aurora A | Aurora Kinase | SelleckChem, L1900 |
| 171 | AMG-900 | Aurora A | Aurora Kinase | SelleckChem, L1900 |
| 171 | AMG-900 | Aurora B | Aurora Kinase | SelleckChem, L1900 |
| 171 | AMG-900 | Aurora C | Aurora Kinase | SelleckChem, L1900 |
| 171 | AMG-900 | Aurora Kinase | Aurora Kinase | SelleckChem, L1900 |
| 172 | Fedratinib (SAR302503, TG10 | JAK | Kinase | SelleckChem, L1900 |
| 173 | GSK1070916 | Aurora Kinase | Aurora Kinase | SelleckChem, L1900 |
| 173 | GSK1070916 | Aurora B | Aurora Kinase | SelleckChem, L1900 |
| 173 | GSK1070916 | Aurora C | Aurora Kinase | SelleckChem, L1900 |
| 174 | CUDC-907 | HDAC1 | Histone Deacetylase | SelleckChem, L1900 |
| 174 | CUDC-907 | HDAC2 | Histone Deacetylase | SelleckChem, L1900 |
| 174 | CUDC-907 | HDAC3 | Histone Deacetylase | SelleckChem, L1900 |
| 174 | CUDC-907 | HDAC10 | Histone Deacetylase | SelleckChem, L1900 |
| 174 | CUDC-907 | HDAC | Histone Deacetylase | SelleckChem, L1900 |
| 174 | CUDC-907 | PI3Kα | Kinase | SelleckChem, L1900 |
| 175 | MK-5108 (VX-689) | Aurora A | Aurora Kinase | SelleckChem, L1900 |
| 175 | MK-5108 (VX-689) | Aurora Kinase | Aurora Kinase | SelleckChem, L1900 |

| | | | | |
|---|---|---|---|---|
| 176 | M344 | HDAC | Histone Deacetylase | SelleckChem, L1900 |
| 177 | Tofacitinib (CP-690550,Tasocit | JAK | Kinase | SelleckChem, L1900 |
| 178 | WP1066 | JAK2 | Kinase | SelleckChem, L1900 |
| 178 | WP1066 | JAK | Kinase | SelleckChem, L1900 |
| 178 | WP1066 | STAT3 | Transcription factor | SelleckChem, L1900 |
| 179 | Sirtinol | Sirt1 | Histone Deacetylase | SelleckChem, L1900 |
| 179 | Sirtinol | Sirtuin | Histone Deacetylase | SelleckChem, L1900 |
| 180 | CEP-33779 | JAK 2 | Kinase | SelleckChem, L1900 |
| 181 | Tacedinaline (CI994) | HDAC1 | Histone Deacetylase | SelleckChem, L1900 |
| 181 | Tacedinaline (CI994) | HDAC2 | Histone Deacetylase | SelleckChem, L1900 |
| 181 | Tacedinaline (CI994) | HDAC3 | Histone Deacetylase | SelleckChem, L1900 |
| 181 | Tacedinaline (CI994) | HDAC | Histone Deacetylase | SelleckChem, L1900 |
| 182 | RG108 | Transferase | DNA Methyltransferase | SelleckChem, L1900 |
| 183 | Baricitinib (LY3009104, INCB0 | JAK2 | Kinase | SelleckChem, L1900 |
| 183 | Baricitinib (LY3009104, INCB0 | JAK 1 | Kinase | SelleckChem, L1900 |
| 184 | WHI-P154 | JAK3 | Kinase | SelleckChem, L1900 |
| 184 | WHI-P154 | EGFR | Receptor | SelleckChem, L1900 |
| 185 | PJ34 | PARP | DNA damage Signaling | SelleckChem, L1900 |
| 186 | S-Ruxolitinib (INCB018424) | JAK1 | Kinase | SelleckChem, L1900 |
| 186 | S-Ruxolitinib (INCB018424) | JAK2 | Kinase | SelleckChem, L1900 |
| 186 | S-Ruxolitinib (INCB018424) | JAK | Kinase | SelleckChem, L1900 |
| 187 | IOX2 | HIF | Transcription factor | SelleckChem, L1900 |
| 188 | Clevudine | DNA/RNA | DNA/RNA Synthesis | SelleckChem, L1900 |
| 189 | Entacapone | COMT | Histone Methyltransferase | SelleckChem, L1900 |
| 190 | Sodium Phenylbutyrate | HDAC | Histone Deacetylase | SelleckChem, L1900 |
| 191 | Tranylcypromine (2-PCPA) HC | MAO | Histone Deacetylase | SelleckChem, L1900 |
| 191 | Tranylcypromine (2-PCPA) HC | CYP2A6 | Monooxygenase | SelleckChem, L1900 |
| 192 | Procainamide HCl | DNA Methyltransferase | DNA Methyltransferase | SelleckChem, L1900 |
| 193 | Tofacitinib (CP-690550) Citrate | JAK | Kinase | SelleckChem, L1900 |
| 194 | Gemcitabine | DNA/RNA | DNA/RNA Synthesis | SelleckChem, L1900 |
| 195 | Carboplatin | DNA/RNA | DNA/RNA Synthesis | SelleckChem, L1900 |
| 196 | Daptomycin | DNA/RNA | DNA/RNA Synthesis | SelleckChem, L1900 |
| 197 | Mizorribine | DNA/RNA | DNA/RNA Synthesis | SelleckChem, L1900 |
| 198 | Cytarabine | DNA/RNA | DNA/RNA Synthesis | SelleckChem, L1900 |
| 199 | Nedaplatin | DNA/RNA | DNA/RNA Synthesis | SelleckChem, L1900 |
| 200 | Procarbazine | DNA/RNA | DNA/RNA Synthesis | SelleckChem, L1900 |
| 201 | Blasticidin | DNA/RNA | DNA/RNA Synthesis | SelleckChem, L1900 |
| 202 | APTSTAT3 | STAT3 | Transcription factor | SelleckChem, L1900 |

**Table S3. Drug efficacy across the PDO cohort.**
3rd-4th column: median drug viability across the PDO cohort at 72h and 144h per each individual compound
5th-6th column: the number of PDO models achieving ≤25% and ≤50% viability at 144h

| Inhibitor | Family | Median %Viability | | Viability threshold 144h | |
|---|---|---|---|---|---|
| | | 72h | 144h | 25% | 50% |
| (+)-JQ1 | Epigenetic Reader Domain | 58.21430288 | 27.15005667 | 0 | 4 |
| 2-Methoxyestradiol (2-MeOE2) | Transcription factor | 88.19521767 | 86.62548209 | 0 | 3 |
| 3-deazaneplanocin A (DZNeP) HCl | Histone Methyltransferase | 58.59509179 | 53.26709131 | 1 | 4 |
| 4SC-202 | Histone Deacetylase | 71.02859198 | 70.13535667 | 9 | 19 |
| A-196 | Histone Methyltransferase | 40.10677411 | 33.1967994 | 0 | 6 |
| A-366 | Histone Methyltransferase | 71.40946899 | 61.99691039 | 0 | 0 |
| Abexinostat (PCI-24781) | Histone Deacetylase | 83.54238461 | 75.95846347 | 5 | 20 |
| AG-14361 | DNA damage Signaling | 91.87009334 | 98.70611854 | 0 | 1 |
| AG-490 (Tyrphostin B42) | Kinase | 86.45307251 | 76.23314396 | 0 | 1 |
| Alisertib (MLN8237) | Aurora Kinase | 86.2081904 | 86.11340667 | 0 | 2 |
| AMG-900 | Aurora Kinase | 93.59976208 | 92.31795494 | 0 | 6 |
| Anacardic Acid | Histone Acetyltransferase | 72.55624953 | 63.33250381 | 0 | 0 |
| APTSTAT3 | Transcription factor | 104.4647711 | 108.9266 | 0 | 6 |
| AR-42 | Histone Deacetylase | 92.82446482 | 83.72320591 | 11 | 21 |
| AT9283 | Kinase | 86.44080605 | 81.76535567 | 0 | 11 |
| Aurora A Inhibitor I | Aurora Kinase | 88.96828403 | 81.30786385 | 1 | 7 |
| AZ 960 | Kinase | 92.49584261 | 91.68463667 | 2 | 15 |
| AZ6102 | Transcription factor activator | 75.61136293 | 72.98468811 | 6 | 20 |
| Azacitidine | DNA Methyltransferase | 90.88371772 | 77.48930085 | 0 | 0 |
| AZD1208 | Pim-Kinase | 58.01154525 | 43.12659333 | 0 | 1 |
| AZD1480 | Kinase | 91.4055296 | 87.53939667 | 1 | 6 |
| AZD2461 | DNA damage Signaling | 51.41863287 | 24.86103192 | 0 | 2 |
| AZD5153 | Epigenetic Reader Domain | 43.47702236 | 33.24427924 | 1 | 6 |
| Barasertib (AZD1152-HQPA) | Aurora Kinase | 86.6559282 | 79.77719333 | 0 | 3 |
| Baricitinib (LY3009104, INCB028050) | Kinase | 96.52875871 | 95.644691 | 2 | 8 |
| Belinostat (PXD101) | Histone Deacetylase | 82.9796515 | 85.34874667 | 1 | 12 |
| BI-7273 | Epigenetic Reader Domain | 79.97394289 | 77.54815084 | 0 | 0 |
| BIX 01294 | Histone Methyltransferase | 77.40978114 | 55.12947718 | 1 | 13 |
| Blasticidin | DNA/RNA Synthesis | 104.2604206 | 106.6311456 | 7 | 18 |
| BRD4770 | Histone Methyltransferase | 72.66240424 | 66.20716173 | 0 | 0 |
| Bromosporine | Epigenetic Reader Domain | 63.23281822 | 56.28634723 | 0 | 5 |
| C646 | Histone Acetyltransferase | 58.70185048 | 45.20988333 | 0 | 0 |
| Carboplatin | DNA/RNA Synthesis | 100.5356353 | 103.70989 | 0 | 6 |
| CEP-33779 | Kinase | 95.89385843 | 90.44774742 | 0 | 0 |
| Clevudine | DNA/RNA Synthesis | 97.13584616 | 100.4079597 | 0 | 0 |
| CP2 | Histone Demethylase | 47.81927311 | 38.27142256 | 1 | 4 |
| CPI-0610 | Epigenetic Reader Domain | 37.42192463 | 14.90205451 | 0 | 9 |
| CPI-1205 | Histone Methyltransferase | 44.09417246 | 44.6103658 | 1 | 3 |
| CPI-169 | Histone Methyltransferase | 73.85857418 | 73.3624674 | 0 | 0 |
| CPI-203 | Epigenetic Reader Domain | 68.01194441 | 51.83048576 | 0 | 0 |
| CPI-360 | Histone Methyltransferase | 74.42346393 | 66.74208173 | 0 | 2 |
| CPI-455 HCL | Histone Demethylase | 41.25956906 | 32.25386 | 0 | 0 |
| CPI-637 | Epigenetic Reader Domain | 40.80475471 | 33.38467864 | 1 | 6 |
| CUDC-101 | Histone Deacetylase | 87.43190988 | 84.75833681 | 0 | 15 |
| CUDC-907 | Histone Deacetylase | 94.12340304 | 91.09481803 | 10 | 23 |
| CX-6258 HCl | Pim-Kinase | 53.29202418 | 41.70454462 | 5 | 20 |
| CYC116 | Aurora Kinase | 87.164476 | 77.60090063 | 0 | 1 |
| Cytarabine | DNA/RNA Synthesis | 102.0407822 | 97.85525497 | 0 | 3 |
| Dacinostat (LAQ824) | Histone Deacetylase | 83.9656404 | 72.03402257 | 14 | 23 |
| Danusertib (PHA-739358) | Aurora Kinase | 84.90600991 | 82.10646029 | 2 | 5 |
| Daphnetin | Kinase | 92.9162603 | 86.24003792 | 0 | 4 |
| Daptomycin | DNA/RNA Synthesis | 100.5517752 | 107.297835 | 0 | 1 |
| Decernotinib (VX-509) | Kinase | 70.78791018 | 53.87418712 | 0 | 1 |
| Decitabine | DNA Methyltransferase | 87.95065709 | 89.82501039 | 0 | 2 |
| Droxinostat | Histone Deacetylase | 88.8912831 | 85.86423185 | 0 | 12 |
| EED226 | Histone Methyltransferase complex | 46.36613695 | 41.87110821 | 0 | 1 |
| EI1 | Histone Methyltransferase | 73.79069051 | 78.61470298 | 0 | 0 |
| Ellagic acid | Topoisomerase | 88.51705524 | 80.88375232 | 0 | 2 |
| ENMD-2076 | Kinase | 87.42450794 | 90.62021507 | 1 | 4 |
| ENMD-2076 L-(+)-Tartaric acid | Aurora Kinase | 91.07301249 | 92.16523894 | 1 | 7 |
| Entacapone | Histone Methyltransferase | 98.15418514 | 100.7609307 | 0 | 1 |
| Entinostat (MS-275) | Histone Deacetylase | 82.52785507 | 82.55791675 | 1 | 6 |
| EPZ004777 | Histone Methyltransferase | 68.894647 | 69.531815 | 0 | 0 |
| EPZ011989 | Histone Methyltransferase | 76.39773467 | 82.7033042 | 0 | 1 |
| EPZ015666(GSK3235025) | Histone Methyltransferase | 75.47178906 | 70.71703729 | 0 | 1 |
| EPZ020411 | Histone Methyltransferase | 33.24251328 | 29.58079658 | 0 | 3 |
| Fedratinib (SAR302503, TG101348) | Kinase | 93.98343041 | 91.35828747 | 5 | 17 |
| Filgotinib (GLPG0634) | Kinase | 73.28560089 | 47.53763062 | 1 | 4 |
| Gandotinib (LY2784544) | Kinase | 91.92855802 | 84.49814694 | 0 | 3 |
| Gemcitabine | DNA/RNA Synthesis | 99.78595762 | 100.0052102 | 1 | 3 |
| Givinostat (ITF2357) | Histone Deacetylase | 91.69636977 | 86.10810716 | 2 | 17 |
| GSK J1 | Histone Demethylase | 72.51396728 | 53.46672413 | 0 | 0 |
| GSK J4 HCl | Histone Demethylase | 54.8455468 | 45.7304664 | 2 | 18 |
| GSK1070916 | Aurora Kinase | 94.12078637 | 92.43466402 | 5 | 11 |
| GSK1324726A (I-BET726) | Epigenetic Reader Domain | 74.08898577 | 79.32611388 | 0 | 2 |
| GSK2801 | Epigenetic Reader Domain | 62.6178125 | 33.97351902 | 0 | 0 |
| GSK2879552 2HCl | Histone Demethylase | 75.99337741 | 74.99765751 | 0 | 0 |
| GSK503 | Histone Methyltransferase | 76.01382616 | 75.25584527 | 0 | 1 |
| GSK591 | Histone Methyltransferase | 79.36193039 | 80.22657903 | 0 | 1 |
| GSK6853 | Epigenetic Reader Domain | 40.83718302 | 31.51723113 | 0 | 1 |
| GSK-LSD1 2HCl | Histone Demethylase | 71.66204234 | 71.75116207 | 0 | 0 |
| Hesperadin | Aurora Kinase | 90.71979557 | 90.86177876 | 7 | 19 |
| HLCL-61 HCL | Histone Methyltransferase | 81.03583516 | 82.96211618 | 0 | 0 |
| I-BET-762 | Epigenetic Reader Domain | 58.96602277 | 51.24067681 | 0 | 2 |
| I-BRD9 | Epigenetic Reader Domain | 76.86326133 | 71.28561274 | 0 | 1 |
| Iniparib (BSI-201) | DNA damage Signaling | 83.14170222 | 72.71954599 | 0 | 4 |
| INO-1001 (3-Aminobenzamide) | DNA damage Signaling | 85.34208273 | 83.49112866 | 0 | 2 |
| IOX1 | Histone Demethylase | 63.28667923 | 60.53349017 | 0 | 0 |
| IOX2 | Transcription factor | 97.07018331 | 95.66978127 | 0 | 2 |
| ITSA-1 (ITSA1) | Histone Deacetylase | 81.06786894 | 80.41799919 | 0 | 0 |
| JIB-04 | Histone Demethylase | 65.77849443 | 63.15509667 | 0 | 3 |
| JNJ-64619178 | Histone Methyltransferase | 48.83127555 | 52.52148 | 6 | 15 |
| JNJ-7706621 | Aurora Kinase | 88.37371833 | 83.41106 | 0 | 5 |
| KW-2449 | Aurora Kinase | 91.16799205 | 76.05927424 | 0 | 7 |
| Lomeguatrib | DNA Methyltransferase | 78.48119621 | 74.01412206 | 0 | 0 |

| | | | | | |
|---|---|---|---|---|---|
| M344 | Histone Deacetylase | 94.60154059 | 86.38010665 | 1 | 17 |
| MC1568 | Histone Deacetylase | 89.88265298 | 87.68071333 | 0 | 2 |
| ME0328 | DNA damage Signaling | 70.23123594 | 72.47106763 | 0 | 0 |
| MG149 | Histone Acetyltransferase | 70.71710769 | 67.35194861 | 0 | 0 |
| MI-2 | Histone Methyltransferase | 73.94969901 | 76.2643203 | 0 | 0 |
| MI-3 | Histone Methyltransferase | 74.08798093 | 74.9927487 | 0 | 0 |
| MI-463 | Histone Methyltransferase complex | 29.21898242 | 27.43467563 | 5 | 22 |
| MI-503 | Histone Methyltransferase complex | 31.90505736 | 24.70325773 | 11 | 23 |
| Mirin | DNA damage Signaling | 79.30641978 | 76.78927357 | 0 | 0 |
| Mitomycin C | DNA/RNA Synthesis | 79.81895469 | 81.12711845 | 7 | 24 |
| Mivebresib (ABBV-075) | Epigenetic Reader Domain | 46.23373456 | 45.40885 | 2 | 13 |
| Mizorribine | DNA/RNA Synthesis | 101.9274161 | 100.9261096 | 0 | 1 |
| MK-5108 (VX-689) | Aurora Kinase | 94.55041828 | 97.48292 | 1 | 11 |
| ML324 | Histone Demethylase | 67.30225834 | 57.04445333 | 1 | 1 |
| MLN8054 | Aurora Kinase | 84.41101557 | 77.63244334 | 0 | 0 |
| MM-102 | Histone Methyltransferase | 64.6311085 | 48.67717848 | 0 | 3 |
| Mocetinostat (MGCD0103) | Histone Deacetylase | 84.95781247 | 88.07341667 | 20 | 22 |
| Momelotinib (CYT387) | Kinase | 92.65751013 | 88.43131109 | 0 | 2 |
| MS023 | Histone Methyltransferase | 79.49922545 | 71.26358086 | 0 | 0 |
| MS049 | Histone Methyltransferase | 40.54636932 | 40.21741419 | 0 | 3 |
| MS436 | Epigenetic Reader Domain | 68.38700327 | 63.82249212 | 0 | 0 |
| Nedaplatin | DNA/RNA Synthesis | 102.7589508 | 100.9657604 | 3 | 13 |
| Nexturastat A | Histone Deacetylase | 70.36009517 | 77.47279746 | 0 | 2 |
| Niraparib (MK-4827) tosylate | DNA damage Signaling | 74.16001177 | 72.65554046 | 0 | 0 |
| Norfloxacin | Topoisomerase | 90.0636898 | 87.32503291 | 0 | 0 |
| NVP-BSK805 2HCl | Kinase | 93.05418164 | 96.48061177 | 4 | 19 |
| NVP-TNKS656 | DNA damage Signaling | 63.45025529 | 62.429847 | 0 | 2 |
| Oclacitinib | Kinase | 80.72113724 | 79.92759145 | 0 | 3 |
| OF-1 | Epigenetic Reader Domain | 75.12529515 | 76.84119757 | 0 | 1 |
| Ofloxacin | Topoisomerase | 89.77123868 | 83.76818667 | 0 | 0 |
| OG-L002 | Histone Demethylase | 63.38163156 | 62.7314502 | 0 | 0 |
| OICR-9429 | Histone | 35.6258818 | 32.02309667 | 0 | 0 |
| Olaparib (AZD2281, Ku-0059436) | DNA damage Signaling | 82.80646949 | 75.14485432 | 0 | 1 |
| ORY-1001 (RG-6016) 2HCl | Histone Demethylase | 75.78816199 | 68.31543333 | 0 | 0 |
| OTX015 | Epigenetic Reader Domain | 69.38778989 | 39.19929667 | 0 | 2 |
| Pacritinib (SB1518) | Kinase | 78.66402409 | 65.13878353 | 1 | 13 |
| Panobinostat (LBH589) | Histone Deacetylase | 81.81206826 | 84.02231326 | 4 | 18 |
| PCI-34051 | Histone Deacetylase | 90.94139496 | 81.68830333 | 0 | 2 |
| PF-06726304 | Histone | 46.34813713 | 43.4827777 | 2 | 6 |
| PF-CBP1 HCl | Epigenetic Reader Domain | 80.54714016 | 79.87442582 | 0 | 1 |
| PFI-1 (PF-6405761) | Epigenetic Reader Domain | 87.95615197 | 90.07353485 | 0 | 1 |
| PFI-2 HCl | Histone Methyltransferase | 66.43531831 | 62.16048501 | 0 | 0 |
| PFI-3 | Epigenetic Reader Domain | 68.5206127 | 46.13798231 | 0 | 0 |
| PHA-680632 | Aurora Kinase | 89.18510666 | 79.61251262 | 0 | 0 |
| Pinometostat (EPZ5676) | Histone Methyltransferase | 53.80044208 | 51.37962333 | 0 | 2 |
| Pirarubicin | Topoisomerase | 88.6205403 | 78.57578356 | 2 | 16 |
| PJ34 | DNA damage Signaling | 96.6827703 | 90.59411049 | 0 | 0 |
| PJ34 HCl | DNA damage Signaling | 68.00745635 | 61.86996509 | 0 | 1 |
| Pracinostat (SB939) | Histone Deacetylase | 90.15306708 | 88.59545512 | 2 | 18 |
| Procainamide HCl | DNA Methyltransferase | 99.57702157 | 105.5338977 | 0 | 0 |
| Procarbazine | DNA/RNA Synthesis | 103.9336533 | 101.1318461 | 0 | 3 |
| Quercetin | Histone Deacetylase | 92.840814 | 90.64924 | 0 | 0 |
| Quisinostat (JNJ-26481585) 2HCl | Histone Deacetylase | 84.06837403 | 78.91287333 | 13 | 23 |
| Remodelin | Histone Acetyltransferase | 74.27015325 | 80.11981537 | 0 | 0 |
| Resminostat | Histone Deacetylase | 93.29113 | 91.77726336 | 4 | 17 |
| Resveratrol | Histone Deacetylase | 88.64083357 | 88.11051242 | 0 | 4 |
| RG108 | DNA Methyltransferase | 96.35754627 | 99.6070683 | 0 | 1 |
| RG2833 (RGFP109) | Histone Deacetylase | 65.88622109 | 43.43754093 | 0 | 1 |
| RGFP966 | Histone Deacetylase | 62.01656765 | 41.45273894 | 0 | 3 |
| Ricolinostat (ACY-1215) | Histone Deacetylase | 76.90507894 | 64.72183 | 0 | 1 |
| Roxadustat (FG-4592) | Transcription factor | 81.57940606 | 75.22434 | 0 | 0 |
| Rucaparib (AG-014699,PF-01367338) | DNA damage Signaling | 84.35314603 | 68.45086626 | 0 | 2 |
| Ruxolitinib (INCB018424) | Kinase | 88.53111743 | 87.22328684 | 0 | 0 |
| RVX-208 | Epigenetic Reader Domain | 66.59993581 | 62.05800872 | 0 | 0 |
| Scriptaid | Histone Deacetylase | 77.93934167 | 82.04385112 | 0 | 1 |
| Selisistat (EX 527) | Histone Deacetylase | 90.7960918 | 82.26261049 | 0 | 1 |
| SF2523 | Epigenetic Reader Domain | 47.63092507 | 39.94602042 | 1 | 9 |
| SGC 0946 | Histone Methyltransferase | 55.00492146 | 60.26823107 | 0 | 3 |
| SGC2085 | Histone Methyltransferase | 41.76586996 | 29.41174333 | 1 | 2 |
| SGC707 | Histone Methyltransferase | 76.45289705 | 71.679275 | 0 | 0 |
| SGC-CBP30 | Epigenetic Reader Domain | 63.57287279 | 73.13257746 | 0 | 9 |
| SGI-1027 | DNA Methyltransferase | 65.71781499 | 41.00794667 | 1 | 19 |
| SGI-1776 free base | Kinase | 92.2896049 | 92.42529233 | 4 | 19 |
| Sirtinol | Histone Deacetylase | 95.59159743 | 93.12723125 | 0 | 1 |
| SMI-4a | Pim-Kinase | 77.40102703 | 70.39268315 | 0 | 0 |
| SNS-314 Mesylate | Aurora Kinase | 86.73404908 | 70.92765766 | 1 | 2 |
| Sodium Phenylbutyrate | Histone Deacetylase | 98.22082603 | 103.116119 | 0 | 0 |
| SP2509 | Histone Demethylase | 74.48758682 | 60.7612844 | 2 | 21 |
| SRT1720 HCl | Histone Deacetylase | 85.00782772 | 78.02768362 | 15 | 23 |
| S-Ruxolitinib (INCB018424) | Kinase | 96.75099643 | 92.02556643 | 0 | 3 |
| Tacedinaline (CI994) | Histone Deacetylase | 96.02054092 | 76.23476151 | 1 | 7 |
| TAK-901 | Aurora Kinase | 93.39256614 | 92.09521401 | 8 | 26 |
| TG101209 | Kinase | 93.07147333 | 80.64104663 | 6 | 17 |
| TMP269 | Histone Deacetylase | 68.8069284 | 63.77315743 | 0 | 0 |
| Tofacitinib (CP-690550) Citrate | Kinase | 99.66578082 | 99.56833563 | 0 | 1 |
| Tofacitinib (CP-690550,Tasocitinib) | Kinase | 94.61783941 | 90.56576447 | 0 | 0 |
| Tozasertib (VX-680, MK-0457) | Aurora Kinase | 82.50361414 | 82.4001966 | 0 | 4 |
| Tranylcypromine (2-PCPA) HCl | Histone Deacetylase | 99.0818497 | 109.0974911 | 0 | 0 |
| Trichostatin A (TSA) | Histone Deacetylase | 81.9345051 | 81.51365667 | 7 | 21 |
| Tubastatin A | Histone Deacetylase | 77.99991178 | 82.46397585 | 0 | 0 |
| Tubastatin A HCl | Histone Deacetylase | 93.05384415 | 98.81852333 | 0 | 3 |
| UNC0379 | Histone Methyltransferase | 71.17884765 | 74.75041333 | 10 | 24 |
| UNC0631 | Histone Methyltransferase | 73.34979337 | 71.02901232 | 1 | 11 |
| UNC0638 | Histone Methyltransferase | 40.259482 | 36.15840996 | 6 | 25 |
| UNC1215 | Epigenetic Reader Domain | 56.0985858 | 50.94476484 | 0 | 0 |
| UNC3866 | Histone Methyltransferase complex | 45.59027463 | 49.60401394 | 2 | 3 |
| UNC669 | Epigenetic Reader Domain | 69.65223344 | 65.75892906 | 0 | 0 |
| UPF 1069 | DNA damage Signaling | 77.67860805 | 56.53945628 | 2 | 2 |
| Valproic acid sodium salt (Sodium valp | Histone Deacetylase | 86.87697594 | 78.4945712 | 0 | 0 |
| Veliparib (ABT-888) | DNA damage Signaling | 81.54566303 | 73.72162333 | 0 | 0 |
| Vorinostat (SAHA, MK0683) | Histone Deacetylase | 82.44097983 | 77.33260667 | 0 | 12 |
| WHI-P154 | Kinase | 96.59454598 | 96.63999503 | 0 | 2 |
| WP1066 | Kinase | 94.66091973 | 99.58184181 | 0 | 1 |
| XL019 | Kinase | 51.47335463 | 51.42750644 | 0 | 0 |
| Zebularine | DNA Methyltransferase | 58.34015568 | 32.62005547 | 0 | 0 |
| ZM 39923 HCl | Kinase | 77.12580891 | 84.36797793 | 0 | 0 |
| ZM 447439 | Aurora Kinase | 84.84831969 | 77.30339055 | 0 | 9 |

**Table S4. Loading scores of the variables identified in the genomic, epigenomic, transcriptomic, and drug layers in the LF1, LF2, and LF4.**
The features with loading scores above 0.75 are displayed, except for the epigenomics layer, which shows markers with the highest loading scores >0.9.

| Latent Factor 1 | | | | Latent Factor 2 | | | | Latent Factor 4 | | | |
|---|---|---|---|---|---|---|---|---|---|---|---|
| **Feature** | **Loading score** | **Annotated gene/ Drug family** | **MOFA layer** | **Feature** | **Loading score** | **Annotated gene/ Drug family** | **MOFA layer** | **Feature** | **Loading score** | **Annotated gene Drug family** | **MOFA layer** |
| PDGFRA | 1.000 | PDGFRA | Genomics | MYCN | 1.000 | MYCN | Genomics | Chr7 | -1.000 | Chr7 | Genomics |
| cg00916884 | 1.000 | MT1M | Epigenomics | Chr2 | 0.748 | Chr2 | Genomics | cg02861441 | 1.000 | LINC00620 | Epigenomics |
| cg20611882 | 1.000 | WLS | Epigenomics | cg19601293 | -1.000 | MAD1L1 | Epigenomics | cg19305111 | 0.984 | PTP4A3 | Epigenomics |
| cg26008007 | 0.997 | NA | Epigenomics | cg16876647 | 0.999 | ZFP36L2 | Epigenomics | cg14480858 | 0.976 | SPATA6L | Epigenomics |
| cg16322929 | 0.996 | SNORA74A | Epigenomics | cg15810758 | 0.983 | NA | Epigenomics | cg11118396 | 0.975 | PTP4A3 | Epigenomics |
| cg17213402 | 0.992 | NA | Epigenomics | cg08878368 | 0.978 | SALL1 | Epigenomics | cg02307165 | 0.972 | NA | Epigenomics |
| cg25745651 | 0.991 | UBB | Epigenomics | cg14344261 | 0.962 | PAX5 | Epigenomics | cg18400845 | 0.969 | RCN1 | Epigenomics |
| cg23167218 | 0.987 | MATR3 | Epigenomics | cg08193650 | 0.933 | ZFP36L2 | Epigenomics | cg10552126 | 0.957 | DPP6 | Epigenomics |
| cg22388634 | 0.987 | VSX1 | Epigenomics | cg12002303 | 0.932 | SKOR1 | Epigenomics | cg18547942 | 0.953 | POU2AF3 | Epigenomics |
| cg03671191 | 0.986 | SPACA6 | Epigenomics | cg14094409 | -0.919 | DIABLO | Epigenomics | cg00467244 | 0.951 | PTP4A3 | Epigenomics |
| cg11549953 | 0.985 | ANXA2R-OT1 | Epigenomics | cg15355952 | 0.918 | SLC1A3 | Epigenomics | cg21794428 | 0.942 | LATS2 | Epigenomics |
| cg24846594 | 0.982 | LGALS8 | Epigenomics | cg08199953 | 0.914 | SALL1 | Epigenomics | cg14791866 | 0.941 | PRXL2A | Epigenomics |
| cg18945842 | 0.979 | SHISA8 | Epigenomics | cg04646987 | 0.913 | TRIO | Epigenomics | cg19099833 | 0.935 | CSRP1 | Epigenomics |
| cg20668644 | 0.977 | RAPGEFL1 | Epigenomics | cg17303756 | 0.911 | POU3F2 | Epigenomics | cg02285477 | 0.929 | LNX2 | Epigenomics |
| cg13912673 | 0.975 | QNG1 | Epigenomics | cg10809560 | 0.909 | NA | Epigenomics | cg13554744 | 0.927 | CSRP1 | Epigenomics |
| cg01507044 | 0.975 | SACS | Epigenomics | cg05342467 | -0.909 | MAD1L1 | Epigenomics | cg26470101 | 0.919 | NA | Epigenomics |
| cg05298677 | 0.974 | ETV7 | Epigenomics | cg01663603 | 0.908 | KCNB1 | Epigenomics | cg25771041 | 0.905 | WWTR1 | Epigenomics |
| cg07309102 | 0.974 | KLF4 | Epigenomics | cg18322569 | -0.904 | BARHL2 | Epigenomics | ENSG00000188732 | -1.000 | FAM221A | Transciptomics |
| cg04134240 | 0.973 | NA | Epigenomics | cg13660126 | 0.903 | DIP2C-AS1 | Epigenomics | ENSG00000250317 | -0.978 | SMIM20 | Transciptomics |
| cg07226193 | 0.970 | SWAP70 | Epigenomics | cg02380531 | 0.902 | POU3F2 | Epigenomics | ENSG00000147592 | -0.972 | LACTB2 | Transciptomics |
| cg17246382 | 0.967 | DNAJA4 | Epigenomics | ENSG00000177807 | -1.000 | KCNJ10 | Transciptomics | ENSG00000130340 | -0.966 | SNX9 | Transciptomics |
| cg08184652 | 0.967 | TBR1 | Epigenomics | ENSG00000162407 | -0.858 | PLPP3 | Transciptomics | ENSG00000198585 | -0.954 | NUDT16 | Transciptomics |
| cg07454803 | 0.965 | MYADM | Epigenomics | ENSG00000079215 | -0.841 | SLC1A3 | Transciptomics | ENSG00000154059 | -0.946 | IMPACT | Transciptomics |
| cg04089426 | 0.965 | LTBP3 | Epigenomics | ENSG00000167614 | -0.819 | TTYH1 | Transciptomics | ENSG00000168214 | -0.929 | RBPJ | Transciptomics |
| cg07147599 | 0.965 | NA | Epigenomics | ENSG00000100427 | -0.798 | MLC1 | Transciptomics | ENSG00000147065 | -0.916 | MSN | Transciptomics |
| cg17976576 | 0.965 | LIMS2 | Epigenomics | CX-6258 HCl | 1.000 | Pim-Kinase | Drugs | ENSG00000127125 | -0.909 | PPCS | Transciptomics |
| cg03603977 | 0.962 | DLGAP3 | Epigenomics | CPI-1205 | -0.831 | Histone Methyltransferase | Drugs | ENSG00000044446 | -0.887 | PHKA2 | Transciptomics |
| cg22321237 | 0.961 | SORBS3 | Epigenomics | EED226 | -0.767 | Histone Methyltransferase complex | Drugs | ENSG00000111879 | -0.883 | FAM184A | Transciptomics |
| cg24934561 | 0.959 | GFOD3P | Epigenomics | XL019 | 0.757 | Kinase | Drugs | ENSG00000103335 | -0.861 | PIEZO1 | Transciptomics |
| cg12794421 | 0.958 | SH2D4A | Epigenomics | | | | | ENSG00000159228 | -0.847 | CBR1 | Transciptomics |
| cg03018796 | 0.957 | NA | Epigenomics | | | | | ENSG00000172965 | -0.829 | MIR4435-2HG | Transciptomics |
| cg03214297 | 0.957 | PICK1 | Epigenomics | | | | | ENSG00000170315 | 0.801 | UBB | Transciptomics |
| cg17631196 | 0.956 | CPQ | Epigenomics | | | | | ENSG00000130813 | -0.784 | SHFL | Transciptomics |
| cg15490944 | 0.955 | FHIP1A | Epigenomics | | | | | ENSG00000122378 | -0.777 | PRXL2A | Transciptomics |
| cg07201456 | 0.955 | CHST6 | Epigenomics | | | | | ENSG00000101460 | -0.771 | MAP1LC3A | Transciptomics |
| cg23220551 | 0.955 | MFSD2B | Epigenomics | | | | | ENSG00000196756 | -0.770 | SNHG17 | Transciptomics |
| cg16705383 | 0.955 | BIRC3 | Epigenomics | | | | | ENSG00000152455 | 0.765 | SUV39H2 | Transciptomics |
| cg08884368 | 0.954 | NME2 | Epigenomics | | | | | ENSG00000188643 | -0.755 | S100A16 | Transciptomics |
| cg05036032 | 0.954 | GRAMD2A | Epigenomics | | | | | ML324 | -1.000 | Histone Demethylase | Drugs |
| cg19070139 | 0.954 | GNMT | Epigenomics | | | | | Zebularine | -0.852 | DNA Methyltransferase | Drugs |
| cg14279121 | 0.953 | OSBPL9 | Epigenomics | | | | | Decernotinib (VX-509) | -0.846 | Kinase | Drugs |
| cg18345115 | 0.952 | PDE4A | Epigenomics | | | | | I-BRD9 | -0.839 | Epigenetic Reader Domain | Drugs |
| cg07843056 | 0.952 | FKBP5 | Epigenomics | | | | | Ricolinostat (ACY-1215) | -0.814 | Histone Deacetylase | Drugs |
| cg05581701 | 0.951 | MT1M | Epigenomics | | | | | RVX-208 | -0.808 | Epigenetic Reader Domain | Drugs |
| cg07141824 | 0.949 | SBF2-AS1 | Epigenomics | | | | | SF2523 | -0.797 | Epigenetic Reader Domain, Kinase | Drugs |
| cg13730743 | 0.949 | SPACA6 | Epigenomics | | | | | RG2833 (RGFP109) | -0.772 | Histone Deacetylase | Drugs |
| cg00873050 | 0.949 | FAM229A | Epigenomics | | | | | TMP269 | -0.766 | Histone Deacetylase | Drugs |
| cg08369872 | 0.947 | DERL3 | Epigenomics | | | | | Nexturastat A | -0.751 | Histone Deacetylase | Drugs |
| cg10456203 | 0.947 | ARHGEF28 | Epigenomics | | | | | | | | |
| cg25221442 | 0.947 | NA | Epigenomics | | | | | | | | |
| cg23833588 | 0.946 | RUBCNL | Epigenomics | | | | | | | | |
| cg03087610 | 0.946 | AIFM2 | Epigenomics | | | | | | | | |
| cg03113878 | 0.945 | NYAP1 | Epigenomics | | | | | | | | |
| cg02009088 | 0.945 | NRG2 | Epigenomics | | | | | | | | |
| cg23725321 | 0.945 | SEC31B | Epigenomics | | | | | | | | |
| cg07207099 | 0.944 | C3orf62 | Epigenomics | | | | | | | | |
| cg05580655 | 0.944 | IQCJ-SCHIP1 | Epigenomics | | | | | | | | |
| cg04976780 | 0.943 | MAGI2 | Epigenomics | | | | | | | | |
| cg02380531 | 0.943 | POU3F2 | Epigenomics | | | | | | | | |
| cg26487948 | 0.943 | ERGIC1 | Epigenomics | | | | | | | | |
| cg17498803 | 0.942 | NA | Epigenomics | | | | | | | | |
| cg16938490 | 0.942 | B3GNT5 | Epigenomics | | | | | | | | |
| cg01591431 | 0.942 | JHY | Epigenomics | | | | | | | | |
| cg12975230 | 0.941 | NA | Epigenomics | | | | | | | | |
| cg26508844 | 0.941 | NME2 | Epigenomics | | | | | | | | |
| cg26458072 | 0.941 | SEC31B | Epigenomics | | | | | | | | |
| cg21972318 | 0.940 | NA | Epigenomics | | | | | | | | |
| cg24465469 | 0.940 | PTPN14 | Epigenomics | | | | | | | | |
| cg14709691 | 0.939 | PIK3R3 | Epigenomics | | | | | | | | |
| cg23878564 | 0.938 | TNK1 | Epigenomics | | | | | | | | |
| cg02687055 | 0.938 | MFSD2B | Epigenomics | | | | | | | | |
| cg17792315 | 0.938 | ARHGEF28 | Epigenomics | | | | | | | | |
| cg27205904 | 0.937 | EID3 | Epigenomics | | | | | | | | |
| cg10664112 | 0.937 | KIF5C | Epigenomics | | | | | | | | |
| cg19867649 | 0.937 | DKK3 | Epigenomics | | | | | | | | |
| cg25042239 | 0.936 | CUL7 | Epigenomics | | | | | | | | |
| cg18222083 | 0.936 | TMEM106A | Epigenomics | | | | | | | | |
| cg23693289 | 0.936 | PTK2B | Epigenomics | | | | | | | | |
| cg25835669 | 0.935 | TUBA8 | Epigenomics | | | | | | | | |
| cg12918457 | 0.934 | LINC00900 | Epigenomics | | | | | | | | |
| cg15603424 | 0.934 | BMAL1 | Epigenomics | | | | | | | | |
| cg09939831 | 0.932 | LGALS3 | Epigenomics | | | | | | | | |
| cg14932794 | 0.932 | TOM1L1 | Epigenomics | | | | | | | | |
| cg23243652 | 0.932 | ARHGEF28 | Epigenomics | | | | | | | | |
| cg11593482 | 0.931 | ADPRH | Epigenomics | | | | | | | | |
| cg22295787 | 0.931 | ADRA1A | Epigenomics | | | | | | | | |
| cg04098866 | 0.930 | NA | Epigenomics | | | | | | | | |
| cg24525457 | 0.930 | NYAP1 | Epigenomics | | | | | | | | |
| cg21211480 | 0.930 | TMEM106A | Epigenomics | | | | | | | | |
| cg17824240 | 0.930 | HNRNPF | Epigenomics | | | | | | | | |
| cg25525687 | 0.930 | TGFBR3L | Epigenomics | | | | | | | | |
| cg21010450 | 0.929 | NA | Epigenomics | | | | | | | | |
| cg26163368 | 0.929 | MAPT | Epigenomics | | | | | | | | |
| cg02147681 | 0.928 | RAI1 | Epigenomics | | | | | | | | |
| cg16358679 | 0.928 | DNAJA4 | Epigenomics | | | | | | | | |
| cg22105332 | 0.928 | RNF39 | Epigenomics | | | | | | | | |
| cg21819468 | 0.927 | PDPN | Epigenomics | | | | | | | | |
| cg10063637 | 0.927 | HMOX2 | Epigenomics | | | | | | | | |
| cg15991262 | 0.927 | FAM221A | Epigenomics | | | | | | | | |
| cg05852231 | 0.927 | ACTC1 | Epigenomics | | | | | | | | |
| cg23018092 | 0.927 | MEGF10 | Epigenomics | | | | | | | | |
| cg12025522 | 0.926 | MAGI2 | Epigenomics | | | | | | | | |
| cg25624927 | 0.926 | SLC25A20 | Epigenomics | | | | | | | | |
| cg09234567 | 0.926 | GRAMD2A | Epigenomics | | | | | | | | |
| cg08540953 | 0.926 | NAT16 | Epigenomics | | | | | | | | |
| cg12928379 | 0.925 | DRD4 | Epigenomics | | | | | | | | |
| cg23613219 | 0.925 | ERBB2 | Epigenomics | | | | | | | | |
| cg24662107 | 0.925 | NA | Epigenomics | | | | | | | | |
| cg07019438 | 0.925 | EGLN3 | Epigenomics | | | | | | | | |
| cg12497564 | 0.924 | RBP1 | Epigenomics | | | | | | | | |
| cg13096820 | 0.924 | EFNA3 | Epigenomics | | | | | | | | |
| cg08662757 | 0.923 | SSH3 | Epigenomics | | | | | | | | |
| cg04407470 | 0.923 | NR2E1 | Epigenomics | | | | | | | | |
| cg24016627 | 0.923 | RNF39 | Epigenomics | | | | | | | | |
| cg16016176 | 0.923 | ADPRH | Epigenomics | | | | | | | | |
| cg06285590 | 0.922 | MCAM | Epigenomics | | | | | | | | |
| cg05019168 | 0.922 | TRIP4 | Epigenomics | | | | | | | | |
| cg13506653 | 0.922 | GSX2 | Epigenomics | | | | | | | | |
| cg22580372 | 0.921 | TMEM177 | Epigenomics | | | | | | | | |
| cg18686527 | 0.921 | RAB34 | Epigenomics | | | | | | | | |
| cg05724965 | 0.921 | JHY | Epigenomics | | | | | | | | |
| cg23238119 | 0.921 | C6orf147 | Epigenomics | | | | | | | | |
| cg02863856 | 0.920 | STARD13 | Epigenomics | | | | | | | | |
| cg20409466 | 0.920 | NA | Epigenomics | | | | | | | | |
| cg13883696 | 0.920 | NA | Epigenomics | | | | | | | | |
| cg17091361 | 0.919 | TEKT3 | Epigenomics | | | | | | | | |
| cg14621053 | 0.918 | ADAM12 | Epigenomics | | | | | | | | |
| cg21512644 | 0.918 | NPY2R | Epigenomics | | | | | | | | |
| cg06118384 | 0.918 | KIF5C | Epigenomics | | | | | | | | |
| cg14159304 | 0.918 | CALHM2 | Epigenomics | | | | | | | | |
| cg10913456 | 0.917 | FKBP5 | Epigenomics | | | | | | | | |
| cg15026277 | 0.917 | TMEM106A | Epigenomics | | | | | | | | |
| cg27294816 | 0.917 | MYADM | Epigenomics | | | | | | | | |
| cg23939808 | 0.917 | RNF39 | Epigenomics | | | | | | | | |
| cg13259296 | 0.917 | MFSD2B | Epigenomics | | | | | | | | |
| cg05790451 | 0.916 | APBA2 | Epigenomics | | | | | | | | |
| cg13185030 | 0.915 | LGALS3 | Epigenomics | | | | | | | | |
| cg23520075 | 0.915 | MYADM | Epigenomics | | | | | | | | |
| cg25172604 | 0.915 | NA | Epigenomics | | | | | | | | |
| cg08859095 | 0.915 | TSSK3 | Epigenomics | | | | | | | | |
| cg02352240 | 0.915 | SALL1 | Epigenomics | | | | | | | | |
| cg23332699 | 0.915 | ELL3 | Epigenomics | | | | | | | | |

| | | | |
|---|---|---|---|
| cg01567239 | 0.915 | ANXA2R-OT1 | Epigenomics |
| cg03608167 | 0.915 | CXCR4 | Epigenomics |
| cg11401866 | 0.914 | HSPB1 | Epigenomics |
| cg19936912 | 0.914 | FHIP1A | Epigenomics |
| cg11800832 | 0.913 | CCNP | Epigenomics |
| cg26381313 | 0.913 | SEC31B | Epigenomics |
| cg09093388 | 0.913 | RBPMS2 | Epigenomics |
| cg27403810 | 0.913 | MYCBPAP | Epigenomics |
| cg16831085 | 0.912 | ALOX5 | Epigenomics |
| cg19489797 | 0.912 | DNMT3A | Epigenomics |
| cg03342530 | 0.912 | NA | Epigenomics |
| cg04786141 | 0.912 | NA | Epigenomics |
| cg27637930 | 0.911 | NA | Epigenomics |
| cg26336594 | 0.911 | FHIP1A | Epigenomics |
| cg11889692 | 0.911 | NA | Epigenomics |
| cg02436788 | 0.911 | TEKT3 | Epigenomics |
| cg04178858 | 0.910 | RAPGEFL1 | Epigenomics |
| cg18190187 | 0.910 | DRD1 | Epigenomics |
| cg19681956 | 0.909 | RAB32 | Epigenomics |
| cg18691564 | 0.909 | TTBK1 | Epigenomics |
| cg15684962 | 0.909 | NA | Epigenomics |
| cg00927231 | 0.909 | NA | Epigenomics |
| cg13839457 | 0.909 | LINC00900 | Epigenomics |
| cg25660386 | 0.908 | IQSEC1 | Epigenomics |
| cg24677744 | 0.908 | FAR2 | Epigenomics |
| cg04829853 | 0.908 | HAPLN3 | Epigenomics |
| cg22230604 | 0.908 | NA | Epigenomics |
| cg08149333 | 0.908 | SLC25A24 | Epigenomics |
| cg00129651 | 0.907 | RAPGEFL1 | Epigenomics |
| cg06561044 | 0.907 | CTRL | Epigenomics |
| cg10171063 | 0.906 | FAR2 | Epigenomics |
| cg17303756 | 0.906 | POU3F2 | Epigenomics |
| cg02322203 | 0.905 | HSD11B2 | Epigenomics |
| cg05347878 | 0.905 | LINC00092 | Epigenomics |
| cg20623601 | 0.904 | EPAS1 | Epigenomics |
| cg20308679 | 0.904 | FRZB | Epigenomics |
| cg01806295 | 0.904 | ARHGEF28 | Epigenomics |
| cg00339300 | 0.904 | NA | Epigenomics |
| cg06241292 | 0.904 | C3orf62 | Epigenomics |
| cg14744537 | 0.903 | RAB27B | Epigenomics |
| cg14870461 | 0.903 | EOGT | Epigenomics |
| cg09541379 | 0.903 | CALHM2 | Epigenomics |
| cg01663603 | 0.903 | KCNB1 | Epigenomics |
| cg11967654 | 0.903 | SALL1 | Epigenomics |
| cg12191938 | 0.902 | AGAP11 | Epigenomics |
| cg25032595 | 0.901 | CLDN10 | Epigenomics |
| cg13850625 | 0.901 | TMEM144 | Epigenomics |
| cg01593751 | 0.900 | TOM1L1 | Epigenomics |
| cg00275232 | 0.900 | ZNF296 | Epigenomics |
| cg16556145 | 0.900 | CLDN10 | Epigenomics |
| ENSG00000103671 | -1.000 | TRIP4 | Transciptomics |
| ENSG00000178537 | -0.990 | SLC25A20 | Transciptomics |
| ENSG00000170315 | -0.946 | UBB | Transciptomics |
| ENSG00000006756 | -0.903 | ARSD | Transciptomics |
| ENSG00000147065 | -0.903 | MSN | Transciptomics |
| ENSG00000198715 | -0.864 | GLMP | Transciptomics |
| ENSG00000161267 | -0.850 | BDH1 | Transciptomics |
| ENSG00000112874 | -0.817 | NUDT12 | Transciptomics |
| ENSG00000189337 | -0.814 | KAZN | Transciptomics |
| ENSG00000150456 | -0.797 | EEF1AKMT1 | Transciptomics |
| ENSG00000165912 | -0.774 | PACSIN3 | Transciptomics |
| ENSG00000116977 | -0.770 | LGALS8 | Transciptomics |
| EPZ011989 | -1.000 | Histone Methyltransferase | Drugs |
| RGFP966 | -0.825 | Histone Deacetylase | Drugs |
| CPI-1205 | -0.774 | Histone Methyltransferase | Drugs |
| Sirtinol | -0.766 | Histone Deacetylase | Drugs |
| BRD4770 | -0.758 | Histone Methyltransferase | Drugs |

**Table S5. Gene ontology functional annotation of differentially expressed genes.**

Differential gene expression analysis was performed between
(a) IDH1m (T394, T407, T756) versus IDH1wt models;
(b) within IDH1m models (T756 versus T394/T407);
(c) T841 MYCN amplified GBM model versus other models in the cohort.
Differentially expressed genes were defined at threshold: FDR ≤0.01 and |log2FC|≥1.
Gene ontology (GO) terms are shown separately for up and down-regulated genes. Top 30 GO terms are shown if p-value ≤0.01

| | | | |
|---|---|---|---|
| IDH1m (T394, T407, T756) versus IDH1wt models | | | |
| Upregulated genes (n=264) | | Downregulated genes (n=1683) | |
| GO term | P -value | GO term | P -value |
| GO:0030527~structural constituent of chromatin | 1.37E-14 | GO:0005576~extracellular region | 3.51E-20 |
| GO:0000786~nucleosome | 7.38E-13 | GO:0062023~collagen-containing extracellular matrix | 7.50E-19 |
| GO:0046982~protein heterodimerization activity | 5.58E-10 | GO:0005886~plasma membrane | 8.84E-16 |
| GO:0008494~translation activator activity | 5.98E-06 | GO:0005615~extracellular space | 7.68E-14 |
| GO:0006334~nucleosome assembly | 8.69E-06 | GO:0007155~cell adhesion | 5.06E-13 |
| GO:0045948~positive regulation of translational initiation | 6.85E-04 | GO:0045087~innate immune response | 9.92E-12 |
| GO:0070935~3'-UTR-mediated mRNA stabilization | 7.88E-04 | GO:0009986~cell surface | 1.48E-11 |
| GO:0032200~telomere organization | 0.001451929 | GO:0005201~extracellular matrix structural constituent | 2.86E-11 |
| GO:0002227~innate immune response in mucosa | 0.003628193 | GO:0005788~endoplasmic reticulum lumen | 4.60E-09 |
| GO:0006325~chromatin organization | 0.005626473 | GO:0009615~response to virus | 4.78E-09 |
| GO:0030154~cell differentiation | 0.007040391 | GO:0005604~basement membrane | 2.22E-08 |
| GO:0043505~CENP-A containing nucleosome | 0.009222832 | GO:0045071~negative regulation of viral genome replication | 3.56E-08 |
| GO:0061644~protein localization to CENP-A containing chromatin | 0.009752526 | GO:0031012~extracellular matrix | 5.04E-08 |
| | | GO:0006954~inflammatory response | 8.89E-08 |
| | | GO:0051607~defense response to virus | 1.46E-07 |
| | | GO:0016324~apical plasma membrane | 2.50E-07 |
| | | GO:0070062~extracellular exosome | 3.66E-07 |
| | | GO:0048471~perinuclear region of cytoplasm | 1.11E-06 |
| | | GO:0009897~external side of plasma membrane | 1.23E-06 |
| | | GO:0015293~symporter activity | 5.85E-06 |
| | | GO:0007165~signal transduction | 6.13E-06 |
| | | GO:0071260~cellular response to mechanical stimulus | 7.82E-06 |
| | | GO:0042383~sarcolemma | 8.17E-06 |
| | | GO:0005515~protein binding | 9.70E-06 |
| | | GO:0016020~membrane | 1.00E-05 |
| | | GO:0010628~positive regulation of gene expression | 1.03E-05 |
| | | GO:0048018~receptor ligand activity | 1.05E-05 |
| | | GO:0042803~protein homodimerization activity | 1.15E-05 |
| | | GO:0030198~extracellular matrix organization | 1.19E-05 |
| | | GO:0005102~signaling receptor binding | 1.19E-05 |
| T756 versus T394/T407 | | | |
| Upregulated genes (n=1689) | | Downregulated genes (n=2065) | |
| GO term | P -value | GO term | P -value |
| GO:0009952~anterior/posterior pattern specification | 2.00E-08 | GO:0098978~glutamatergic synapse | 1.96E-20 |
| GO:1990837~sequence-specific double-stranded DNA binding | 2.42E-08 | GO:0045211~postsynaptic membrane | 2.62E-17 |
| GO:0030527~structural constituent of chromatin | 2.75E-08 | GO:0016020~membrane | 5.01E-13 |
| GO:0005829~cytosol | 3.40E-08 | GO:0098982~GABA-ergic synapse | 5.16E-13 |
| GO:0009954~proximal/distal pattern formation | 4.50E-08 | GO:0005886~plasma membrane | 7.57E-13 |
| GO:0005515~protein binding | 4.68E-08 | GO:0043005~neuron projection | 9.26E-13 |
| GO:0000785~chromatin | 2.83E-07 | GO:0014069~postsynaptic density | 3.04E-11 |
| GO:0005737~cytoplasm | 4.32E-07 | GO:0043197~dendritic spine | 1.69E-10 |
| GO:0000786~nucleosome | 3.98E-06 | GO:0030425~dendrite | 3.60E-10 |
| GO:0007411~axon guidance | 6.21E-06 | GO:0005604~basement membrane | 2.14E-09 |
| GO:0045944~positive regulation of transcription by RNA polymerase II | 7.31E-06 | GO:0042734~presynaptic membrane | 2.58E-09 |
| GO:0030182~neuron differentiation | 8.90E-06 | GO:0045202~synapse | 1.38E-08 |
| GO:0003700~DNA-binding transcription factor activity | 1.23E-05 | GO:0007268~chemical synaptic transmission | 1.61E-07 |
| GO:0060065~uterus development | 1.96E-05 | GO:0042803~protein homodimerization activity | 1.74E-07 |
| GO:0070062~extracellular exosome | 2.38E-05 | GO:0043204~perikaryon | 2.06E-07 |
| GO:0001657~ureteric bud development | 2.69E-05 | GO:0030672~synaptic vesicle membrane | 3.83E-07 |
| GO:0005634~nucleus | 3.42E-05 | GO:0098839~postsynaptic density membrane | 4.61E-07 |
| GO:0009653~anatomical structure morphogenesis | 4.03E-05 | GO:0060078~regulation of postsynaptic membrane potential | 4.97E-07 |
| GO:0070161~anchoring junction | 4.46E-05 | GO:0062023~collagen-containing extracellular matrix | 5.45E-07 |
| GO:0032200~telomere organization | 5.34E-05 | GO:0007155~cell adhesion | 6.93E-07 |
| GO:1902895~positive regulation of miRNA transcription | 5.63E-05 | GO:0005509~calcium ion binding | 7.53E-07 |
| GO:0030154~cell differentiation | 1.12E-04 | GO:0032590~dendrite membrane | 7.74E-07 |
| GO:0032991~protein-containing complex | 1.33E-04 | GO:0007616~long-term memory | 1.56E-06 |
| GO:0072659~protein localization to plasma membrane | 1.52E-04 | GO:0007214~gamma-aminobutyric acid signaling pathway | 2.31E-06 |
| GO:1900025~negative regulation of substrate adhesion-dependent cell spreading | 2.02E-04 | GO:0005267~potassium channel activity | 2.87E-06 |
| GO:0000978~RNA polymerase II cis-regulatory region sequence-specific DNA binding | 2.90E-04 | GO:0007411~axon guidance | 6.16E-06 |
| GO:0120163~negative regulation of cold-induced thermogenesis | 3.52E-04 | GO:0030424~axon | 8.99E-06 |
| GO:0003677~DNA binding | 3.97E-04 | GO:0043025~neuronal cell body | 1.06E-05 |
| GO:0000981~DNA-binding transcription factor activity, RNA polymerase II-specific | 3.99E-04 | GO:0044325~transmembrane transporter binding | 1.08E-05 |
| GO:0051015~actin filament binding | 4.33E-04 | GO:0048786~presynaptic active zone | 1.14E-05 |
| T841 versus other models | | | |
| Upregulated genes (n=458) | | Downregulated genes (n=1473) | |
| GO term | P -value | GO term | P -value |
| GO:0005739~mitochondrion | 5.53E-08 | GO:0007155~cell adhesion | 3.56E-36 |
| GO:0005829~cytosol | 1.45E-07 | GO:0005886~plasma membrane | 3.65E-30 |
| GO:0003723~RNA binding | 1.68E-06 | GO:0009986~cell surface | 7.72E-24 |
| GO:0000149~SNARE binding | 5.78E-06 | GO:0062023~collagen-containing extracellular matrix | 5.62E-22 |
| GO:0010807~regulation of synaptic vesicle priming | 2.76E-05 | GO:0007399~nervous system development | 1.92E-17 |
| GO:0099502~calcium-dependent activation of synaptic vesicle fusion | 8.96E-05 | GO:0007156~homophilic cell adhesion via plasma membrane adhesion molecules | 1.52E-14 |
| GO:0005515~protein binding | 1.29E-04 | GO:0045944~positive regulation of transcription by RNA polymerase II | 2.71E-13 |
| GO:0005654~nucleoplasm | 1.84E-04 | GO:0005615~extracellular space | 3.52E-13 |
| GO:0017158~regulation of calcium ion-dependent exocytosis | 2.80E-04 | GO:0005925~focal adhesion | 5.21E-13 |
| GO:0005737~cytoplasm | 3.19E-04 | GO:0043235~receptor complex | 5.49E-13 |
| GO:0019905~syntaxin binding | 3.82E-04 | GO:0005509~calcium ion binding | 7.01E-13 |
| GO:0030424~axon | 4.99E-04 | GO:0005604~basement membrane | 1.21E-12 |
| GO:0061891~calcium ion sensor activity | 7.34E-04 | GO:0005576~extracellular region | 1.31E-12 |
| GO:0043204~perikaryon | 9.58E-04 | GO:0031012~extracellular matrix | 1.47E-12 |
| GO:0030672~synaptic vesicle membrane | 0.001082934 | GO:0001525~angiogenesis | 2.39E-12 |
| GO:0001889~liver development | 0.001382898 | GO:0007411~axon guidance | 2.43E-12 |
| GO:0031045~dense core granule | 0.001663927 | GO:0007165~signal transduction | 6.71E-12 |
| GO:0006417~regulation of translation | 0.001809839 | GO:1902895~positive regulation of miRNA transcription | 2.54E-11 |
| GO:0045956~positive regulation of calcium ion-dependent exocytosis | 0.001835428 | GO:0030198~extracellular matrix organization | 1.92E-10 |
| GO:0016192~vesicle-mediated transport | 0.001855973 | GO:0005201~extracellular matrix structural constituent | 2.16E-10 |
| GO:0032922~circadian regulation of gene expression | 0.002123327 | GO:0030335~positive regulation of cell migration | 4.37E-10 |
| GO:0003774~cytoskeletal motor activity | 0.002692805 | GO:0005178~integrin binding | 8.59E-10 |
| GO:0005524~ATP binding | 0.002934223 | GO:0002052~positive regulation of neuroblast proliferation | 1.27E-09 |
| GO:0005741~mitochondrial outer membrane | 0.004087159 | GO:0098839~postsynaptic density membrane | 2.22E-09 |
| GO:0030970~retrograde protein transport, ER to cytosol | 0.004118033 | GO:0005515~protein binding | 3.65E-09 |
| GO:0048471~perinuclear region of cytoplasm | 0.004576441 | GO:0016020~membrane | 5.29E-09 |
| GO:0000836~Hrd1p ubiquitin ligase complex | 0.005081699 | GO:0005912~adherens junction | 5.41E-09 |
| GO:0043022~ribosome binding | 0.005139742 | GO:0007219~Notch signaling pathway | 6.76E-09 |
| GO:0060612~adipose tissue development | 0.005178298 | GO:0098609~cell-cell adhesion | 6.87E-09 |
| GO:0006446~regulation of translational initiation | 0.005178298 | GO:0009897~external side of plasma membrane | 1.70E-08 |

**Table S6. Primary antibodies for Immunohistochemistry (IHC)**

| Antibody | Vendor | Catalog number | Dilution |
|---|---|---|---|
| Nestin | Abcam | ab6320 | 1:500 |
| Vimentin | Milipore | MAB3400 | 1:200 |
| Sox2 | Cell Signaling | 23064 | 1:100 |
| S100 | Chemicon | MAB079-1 | 1:100 |
| Fibronectin 1 | Abcam | ab2413 | 1:100 |
| Collagen VI | Abcam | ab182744 | 1:250 |
| GFAP | Aligent | Z033429-2 | 1:500 |
